# A Pan-Cancer Ex Vivo Drug Screen Atlas for Functional Precision Oncology

**DOI:** 10.64898/2026.02.14.705918

**Authors:** Karl Pichotta, Jessica B. White, Jeffrey F. Quinn, Anneliese Markus, Christopher Tosh, Antoine De Mathelin, Erin Coyne, Feiyang Huang, Wesley Tansey

## Abstract

Compared to immortalized cell lines, patient-derived organoids and other ex vivo models have been shown to better recapitulate patient responses to therapy. High cost and technical complexity have prevented the creation of pan-cancer ex vivo datasets, limiting comprehensive analyses and predictive modeling for ex vivo drug response. We present the Pan-PreClinical (PPC) project: a drug screen atlas of 2.1M experiments across 1,982 ex vivo samples and 3,100 drugs spanning 134 cancer indications tested across 26 studies. We develop a contrastive Bayesian model to harmonize across studies, identifying 303 tissue-specific drug sensitivities and demonstrating drug sensitivities are predictive of clinically-relevant molecular profiles. Integrating established cell line databases reveals systematic biases across 55 cancer subtypes, with cell line screens favoring drugs targeting highly proliferative cells and undervaluing cell-cell communication targets. We leverage PPC to establish an ex vivo foundation model and computational platform for scalable ex vivo cancer biology and predictive oncology.

---

Cancer patients with rare or treatment-resistant tumors are often left without any effective standard of care. The need to identify new treatments for these patients has led to a rapid acceleration of ex vivo model development and high throughput drug screening to identify functional vulnerabilities in these tumors arising from mechanisms not detectable through classical sequencing^1,2^. Compared to immortalized cell lines, ex vivo models like patient-derived cells (PDCs), organoids (PDOs), xenografts (PDXs), and xenograft-derived cells (PDXCs) maintain a higher fidelity match to their original human tissues^3–6^. Thus, positive drug screen hits in the ex vivo setting may have a higher likelihood of translating to successful clinical therapies.

Ex vivo screens in triple negative breast cancer^7^, fibrolamellar carcinoma^8^, glioblastoma^9,10^, pancreatic adenocarcinoma^1,4,11^, acute myeloid leukemia^12,13^, and epithelial ovarian cancers^14–16^ have revealed novel vulnerabilities and new potential personalized therapies for patients. The importance and feasibility of this discovery opportunity is increasing rapidly as large scale efforts to generate ex vivo models for drug screens accelerate within expert PI labs, in institutional organoid biobank programs^17^, and across multi-institutional consortia (e.g. the Pediatric Preclinical In Vivo Testing (PIVOT) consortium^18^). Ideally, these efforts could be streamlined to generate pan-cancer atlases analogous to the Cancer Dependency Map^19^ for immortalized cell lines. Unfortunately, ex vivo models tend to require bespoke protocols for each tissue or disease, limiting streamlined pan-cancer screen databases to date.

The preclinical success of ex vivo screens has also begun to translate into clinical trial success. The short distance between an ex vivo model and its patient of origin enables drug screens on ex vivo models to provide a personalized, functional characterization of a patient’s tumor in terms of vulnerabilities to candidate therapies. The INFORM^20^, TARGET/ZERO^21^, and NCH-FPM trials^22^, all on high-risk pediatric cancers, integrated ex vivo drug screens into the clinical decision making process and demonstrated that screen-guided treatments can improve patient outcomes. These successes have led to the rise of the new field of functional precision oncology (FPO), which posits that ex vivo screens of patient tumors can be used to guide treatment in the clinic^23^. While these studies have yielded substantial evidence for the FPO thesis, systematic analyses have been lacking due to the focused nature of each ex vivo study.

We reasoned that large-scale data integration of ex vivo drug screens would enable predictive modeling and quantitative evaluation of ex vivo drug responses. To this end, we established the Pan-PreClinical (PPC) project, a large open-resource effort to compile drug-response data from ex vivo studies. The PPC dataset currently comprises data from 26 different ex vivo studies, along with genomic, transcriptomic, chemoinformatic, and clinical data where available. All PPC data were integrated into a unified data model with a common cancer subtyping nomenclature, primary/metastasis label, and site-of-sample annotation. We developed a contrastive machine learning algorithm and statistical postprocessing pipeline to harmonize across studies and experimental conditions, enabling pan-cancer analyses. Using the harmonized results, we conducted statistical analyses that reveal tissue-specific differential drug responses, as well as site-specific variation in drug sensitivities across metastatic samples. Integrating 12 large pan-cancer cell line datasets, we found systematic differences in drug sensitivities between ex vivo models and disease-matched cell lines. We leveraged the integrated dataset to train a transformer-based foundation model capable of guiding future studies and showed that drug screen embeddings reliably predict genomic and transcriptomic properties of ex vivo samples. The PPC dataset, model results, and analyses can be explored on an interactive web portal: https://www.panpreclinical.org/

We conducted a literature review of publications involving ex vivo drug screens. Study inclusion criteria were that at least five patient-derived ex vivo samples were screened with at least ten unique compounds. We excluded studies involving only immortalized cell lines or animal models of disease. We identified and obtained data from 26 studies across three categories of patient-derived ex vivo biological modalities, namely assays of patient-derived cells and spheroids (PDCs)^9,10,14–16,20,24–29^, patient-derived organoids^1,4,11,30–36^, patient-derived xenograft-derived cells (PDXCs)^37^, and studies with data from multiple modalities^7,38–41^. For 9 of the studies, raw fluorescence or viability measurements were publicly deposited with the original publication. For the remaining 18 studies, we communicated with the original authors to retrieve and verify the raw data. All data were provided by study authors with consent for sharing previously published data. In one case, additional pre-publication data were provided without the ability to disentangle published from non-published samples. The public version of the PPC dataset omits this study, but we include it here in our analyses.

Through further communication with study authors, we retrieved clinical diagnoses, site-of-disease annotations, and, where available, patient history and tumor molecular information. Each ex vivo sample was matched to a standardized OncoTree cancer type^42^ to give fine-grained disease typing information, with distinctions made between primary and metastatic sample status. We labeled metastatic samples with the same 21-site conventions as the MSK-MET dataset^43^. For metastatic samples, we annotated both the original primary tissue site and the location of the sampled metastasis. Each ex vivo sample was annotated with available genomic, transcriptomic, and clinical information such as sex, treatment history, and overall survival. Studies varied in whether they gathered each additional modality. Regulatory restrictions further prevented us from obtaining and integrating gathered molecular information in some studies. All genomic and transcriptomic data were reprocessed using a standardized pipeline with a common reference genome (see Methods).

## The PPC project unifies ex vivo studies into a single pan-cancer atlas

In total, we collected data on 3,072 ex vivo models, of which 1,982 have drug viability measurements. These represent 130 distinct cancer types (110 with viability measurements) (Fig. 1a,c), from 24 different anatomical sites (23 with viabilities), along with 53 healthy samples (40 with viabilities). Among the non-healthy samples, 2,324 are primary samples (1,834 with viabilities) and 155 are metastatic samples (108 with viabilities). The most frequent ex vivo construct type is PDCs (Fig. 1a), with 2,355 samples (1,574 with viability measurements). We also compiled 595 PDO samples (368 with viabilities) and 104 PDXC samples (37 with viabilities). Drug responses varied across cancer types and drug classes (Fig. 1b).

**Figure 1:**
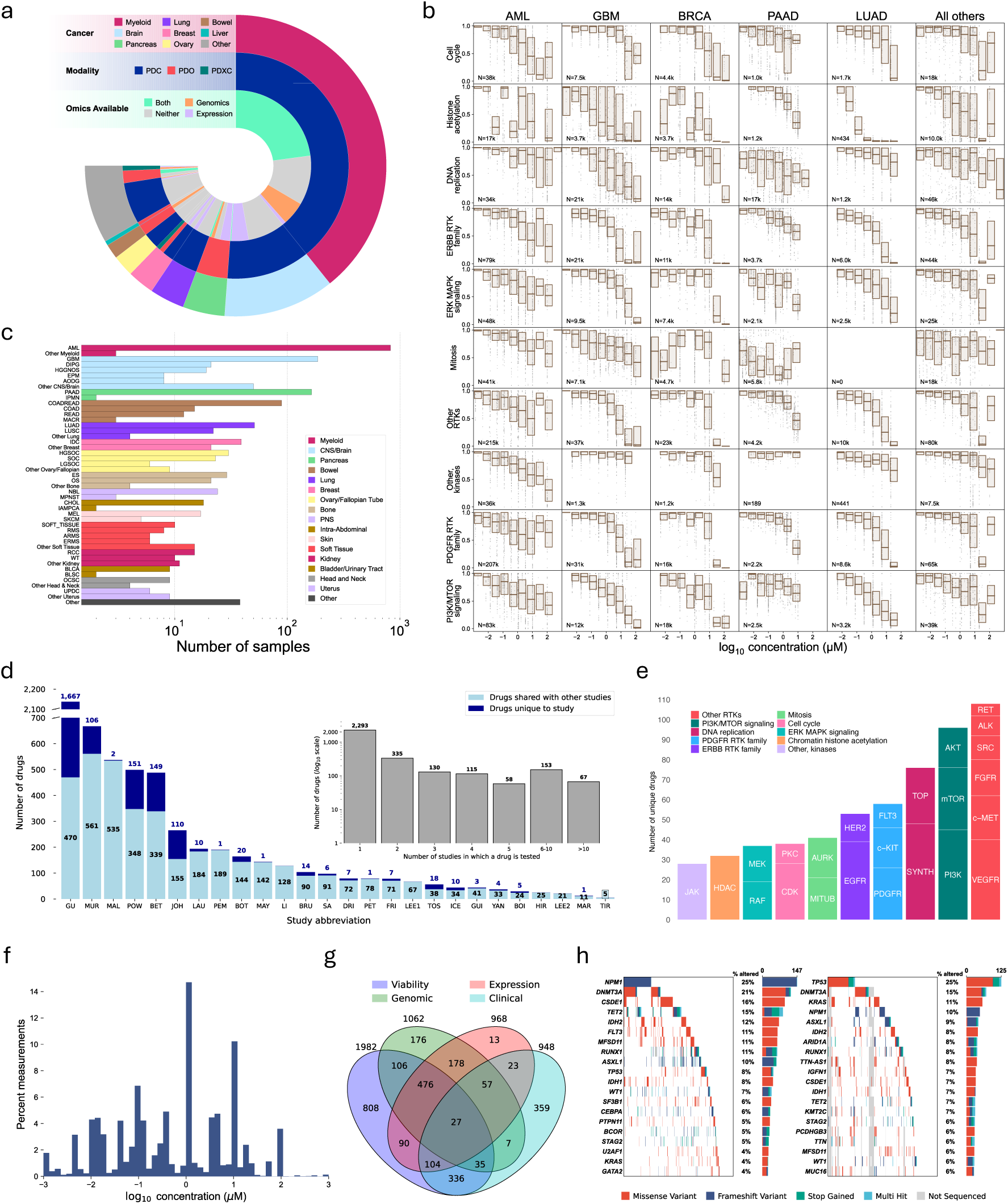
The PPC dataset represents a pan-cancer atlas of ex vivo experiments across diseases, omics, and drugs. (a) Characteristics of samples in the PPC dataset. Width of wedges represents the number of samples. The outer ring represents primary cancer site, the middle ring represents the ex vivo cell culture modality used on the sample, and the innermost ring shows whether transcriptomics and/or genomics results were available for the sample. (b) Viability measurements by OncoTree code (table S2) for top drug targets. (c) Number of samples broken down by primary diagnosis. Diagnoses are abbreviated using their OncoTree codes (table S2). (d) Number of drugs per component study. Studies are abbreviated as specified in table S1. (e) Description of the drugs in the PPC dataset tested in >1% of all samples with the first level grouping by high-level drug category and the second level grouping by specific target. SYNTH=DNA/RNA Synthesis, TOP=Topoisomerase, MITUB=Microtubule Associated, AURK=Aurora Kinase, HEDGE=Hedgehog/Smoothened, all other target abbreviations correspond to Selleckchem mechanism of action labels (Methods). (f) Distribution of drug concentrations across experiments in the PPC dataset. (g) Number of samples annotated with viability, genomic, expression, and clinical data. Clinical data includes patient sex, approximate age at diagnosis, or overall survival. (h) OncoPrint describing mutations for the liquid (left) and solid (right) tumor samples. Excludes genes for which variants were only detected in whole genome or exome sequencing.

Using a custom iterative graph algorithm (see Methods), we harmonized drug names across studies and compiled measurements for 3,151 unique drugs and 2,133,246 individual viability measurements representing 327,335 unique dose-response curves. The median number of doses in each single-drug curve was five, though some studies employed an experimental design involving large libraries of compounds screened at a single dose; 54,351 dose-response curves comprise data at a single drug concentration (fig. S3). Each compound was annotated with FDA approval status, along with annotations for mechanisms of action (fig. S4). Drug concentrations tested in viability measurements spanned from nanomolar to millimolar ranges with a plurality taken at micromolar concentrations (Fig. 1f). Out of 26 ex vivo studies, 12 screened 100 or more compounds (Fig. 1d). Overall, 2,293 drugs (72.7%) were screened in a single study and 858 (27.2%) were screened in two or more studies (Fig. 1d); a majority of compounds tested in each study were also screened in other studies. Using a custom algorithm for drug target annotation (see Methods), we leveraged publicly available resources^44–46^ to annotate 2,219 of the 3,100 drugs with at least one target. The harmonized drug set includes a wide variety of small molecule drugs targeting many common cancer-associated proteins and biological processes and tested in >1% of samples in the PPC dataset (Fig. 1e).

We compiled and harmonized genomics data for 1,062 samples and transcriptomics data for 968 samples; 476 samples had genomics, transcriptomics, and viabilities (Fig. 1g). Five studies used whole genome or exome assays, while six studies used targeted gene panels. Four of the six studies that use gene panels assess alterations in 50 or fewer genes. Demographic and clinical annotations were comparatively limited: 948 samples had patient age, sex, overall survival, or prior treatment history; 336 had clinical annotations and viabilities (Fig. 1g). Constituent studies vary both in coverage across disease type (fig. S1) and drugs (fig. S2).

Both solid and hematological samples are available with genomic alteration data. All hematological samples with annotated genomic alterations (Fig. 1h, left) are derived from AML patients and were assessed using a 5K gene panel. The proportion of these AML samples with the most highly recurrent mutations are consistent with the levels at which they are observed in this cancer type in general, including *NPM1* (25%) and *DNMT3* (21%)^47,48^. For the remaining ten ex vivo studies with genomic alteration data for solid tumor samples, we account for differences in the genes evaluated for mutations in the underlying genomic assays (Fig. 1h, right). Most samples are evaluated for alterations in the most frequently mutated genes, including *T P*53, *DN MT* 3*A*, and *KRAS*.

## Bayesian nonparametric modeling integrates PPC drug screens and generalizes to held out experiments

We sought to integrate across all the PPC component studies to produce a harmonized dataset for comparative analyses. We focused only on integrating drugs used in at least 3 studies, leaving 504 compounds. Heterogeneity in experimental designs across studies makes computational modeling and statistical analyses of the PPC dataset challenging. Overlap in drug libraries between studies varied from 0% to 94%, with an average overlap of 17%, leading to missing measurements across studies. Studies that tested drugs at only a single concentration are not compatible with a simple Hill model curve fitting approach, making full dose-response estimation, as well as summary IC_50_ or area under the curve (AUC) calculation, impossible without statistical modeling. However, while many machine learning models have been developed to predict drug responses from omics^49–52^, the majority of PPC samples do not have omics available. Further, the studies in PPC used different incubation times, culture media, and other experimental factors that lead to batch effects which existing predictive models are not designed to handle. To enable dataset-wide analyses, we developed a custom probabilistic dose-response model.

The PPC dose-response model performs a contrastive tensor factorization on the samples × drugs × concentrations tensor (Fig. 2a). Each ex vivo sample *i* is modeled as a real-valued embedding vector, *v*_*i*_ ∈ R^*d*^. For each drug *j*, the concentration space is divided into *K* discrete points and the drug is modeled as a collection (*u_j_*^(1)^, *u_j_*^(2)^,…, *u_j_*^(*K*)^) of embeddings also in R^*d*^; concentrations falling between grid points are interpolated. The inner product of a specific drug-dose embedding and a sample embedding then represents the log-odds of increasing in viability from the previous grid point; this enforces monotonicity in the inferred curve. To reduce batch effects and inject biological knowledge into the model, the training objective includes contrastive penalties^53^ on the embeddings to encourage samples from the same tissue site to have similar embeddings and drugs with the same mechanism of action to have similar embeddings (see Methods). Once trained, the model was used to impute full monotone-down dose-response curves across the entire samples × drugs space, including sample-drug combinations unobserved during training.

**Figure 2:**
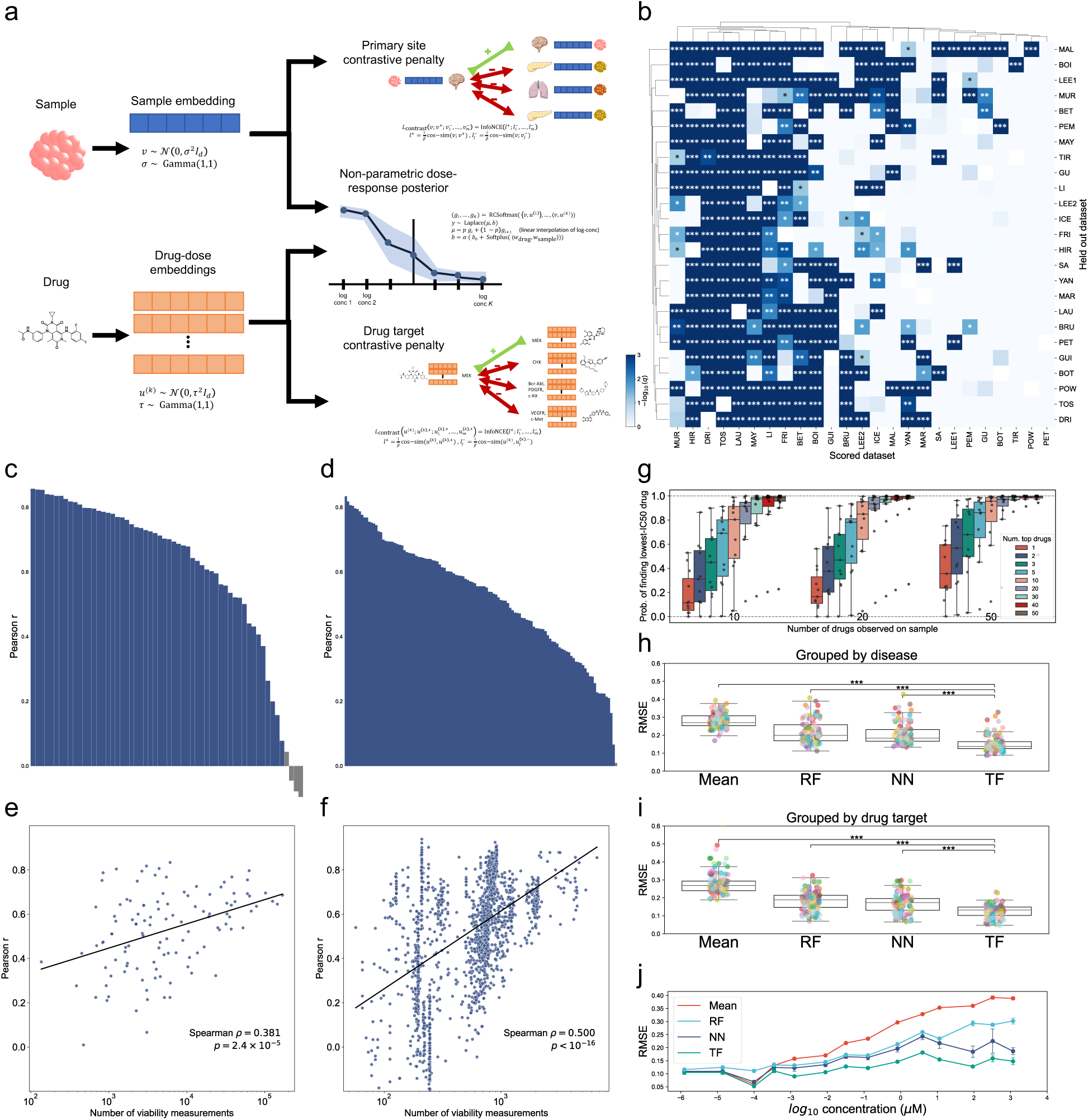
The PPC dose-response model integrates across multiple studies and generalizes to held out experiments. (a) The PPC model performs a constrained tensor factorization to predict curves and impute missing measurements. Primary site and drug mechanism annotations are used as contrastive learning labels to regularize the model. (b) Assessment of which component studies improve generalization on other component studies. Whole-drug-holdout cross-validation on each column dataset was compared with and without having each row dataset in the training set; improvement significance calculated via one-sided binomial test and BH correction. ∗∗∗: *q* < 0.001; ∗∗: *q* < 0.01; ∗: *q* < 0.1 (study abbreviations: table S1). (c,d) Cross-validation performance (Pearson correlation) stratified by (c) disease subtype and (d) drug mechanism; gray: *q* > 0.05 after BH correction (two-sided Pearson correlation t-test); abbreviations: table S2. (e,f) Cross-validation prediction by (Pearson correlation) vs. number viability measurements stratified by (e) drug mechanism and (f) sample; (*p*: two-sided Spearman rank-correlation test). (g) Power analysis benchmark evaluating the number of drugs needed in both a pilot round and a hit prediction round to achieve a target level of power to find the most efficacious drug. (h–j) Benchmarks against baseline machine learning methods, with error stratified by (h) disease, (i) drug target, and (j) log-concentration (two-sided Mann-Whitney U test with BH correction; ***: *q* < 1 × 10^−9^; Mean: bucketed mean; RF: random forest; NN: neural network; TF: PPC Bayesian tensor factorization method (Methods)); (bars: 1 standard deviation).

Given the heterogeneity of the PPC studies, with different experimental designs and labs conducting each experiment, we first asked if data integration was even beneficial. To test this, we conducted a series of ablation studies where we held out a study and evaluated whether the removal of that study harmed or improved cross-validation performance on every other study (see Methods). We found that all datasets provide statistically significant (one-sided binomial test) empirical improvement for at least 50% of other datasets (Fig. 2b). Only one study, PET^20^, saw no statistically significant benefit from any other study though we note that four studies did contribute small improvements that were not significant after multiple testing correction (fig. S6). Studies that were found to be most helpful tended to have larger drug libraries (MAL^26^, MUR^16^) and more diverse samples (LEE1^9^, MAY^29^). The most frequently helpful studies were often the majority of the studies which benefited the least from other studies, suggesting that data integration improvements diminish as studies become broader and more expansive. However, we were only able to evaluate studies on drugs that were measured in those studies. Predicting drug responses for compounds not in the study panel still requires data integration even for the more expansive studies in the PPC dataset.

We next evaluated the model on a series of performance tasks on held-out data to test its robustness. In each task, we performed five-fold cross-validation, holding out entire dose-response curves in the test fold (see Methods). Stratifying by disease subtype, we found variable model performance, ranging from Pearson’s *r* = −0.10 to *r* = 0.86, with median performance of *r* = 0.70 (Fig. 2c, fig. S9). Across all indications, 84% (49 out of 58) had *r* > 0.4, with none having statistically significantly negative correlations (Pearson *r*). Among the diseases on which the model performed best (fig. S7) are endometrial carcinoma (UCEC), uterine sarcoma (USARC), ovarian carcinoma (OVT), and breast carcinoma (BRCA); this reflects differences in data variation, dose-concentration coverage within experiments (fig. S3), and drug coverage within constituent datasets (fig. S2).

Stratifying by mechanism of action (Fig. 2d, fig. S7), we found model performance ranged from Pearson’s *r* = 0.01 to *r* = 0.83, with median *r* = 0.56. Of 116 targets measured, 87 (75%) had *r* > 0.4. Drug classes with fewer measurements had significantly lower predictive accuracy as measured by Pearson *r* (*p* = 2.4 × 10^−5^, two-sided Spearman rank-correlation, Fig. 2e, fig. S10–fig. S13). Further, we found that regions of tighter uncertainty bounds corresponded to regions of higher predictive accuracy (fig. S8), suggesting predictive uncertainty in the model is well-calibrated. For rare disease subtypes and uncommon drugs, we therefore recommend evaluating the PPC model uncertainty estimates to assess the likely fidelity of imputations. Stratifying by sample revealed that samples with more viability measurements had significantly better model accuracy as measured by Pearson *r* (Fig. 2f, *p* < 10^−1^^6^, two-sided Spearman rank-correlation).

To assess the number of measurements needed to obtain a fixed performance threshold, we designed a power analysis benchmark (see Methods). Briefly, samples were held out from the PPC model during initial training, then the model was given a random subset of 10, 20, or 50 drugs as a virtual initial screen. The model was then given a budget of between 1 and 50 more drugs to predict a top hit, defined as the lowest-IC_50_ drug in the panel. We found no clear patterns emerged to suggest an optimal way to balance the initial drug budget versus the top hit prediction budget (Fig. 2g). The model obtained 80% power using two rounds of 10 drugs, 95% power using two rounds of 20 drugs, and > 99% power using two rounds of 50 drugs.

Finally, we sought to assess if the PPC model was exceeding the performance of traditional machine learning methods. We benchmarked the PPC model against both a naive baseline mean estimator and two machine learning methods: random forests and neural networks (see Methods). Evaluating across disease subtype and drug target, we found that the PPC model consistently outperformed all three baselines (Fig. 2h,i). This result was robust when evaluated at different concentration points on the dose-response curve (Fig. 2j), with the PPC model outperforming the baselines at every concentration level.

## Latent embedding spaces capture meaningful relationships between drug mechanisms and tumor sub-types

Using the trained PPC model, we fully harmonized and imputed the PPC dataset then assessed it for biological, chemical, and clinical coherence. Projecting drug and sample embeddings via UMAP^54^ indicated clear separation by both disease subtype (Fig. 3b) and drug mechanism (Fig. 3c). Related disease subtypes like BLSC and BLCA, HCC and IPN, and PAAD and PACT, all cluster together, capturing hierarchical biological structure in diseases. Similarly, drugs inhibiting components within the same pathway, including MEK/ERK and PI3K/Akt/mTOR, cluster together. Therapeutics targeting sex hormones, including androgens and estrogens, are in close proximity which are in turn near PARP inhibitors that are most often used to treat ovarian, breast, prostate, and pancreatic cancer sub-types^55^. Mitotic targets like microtubules and PLK are also clustered near metabolic targets suggesting that the model learns to differentiate agents most effective in rapidly dividing constructs.

**Figure 3:**
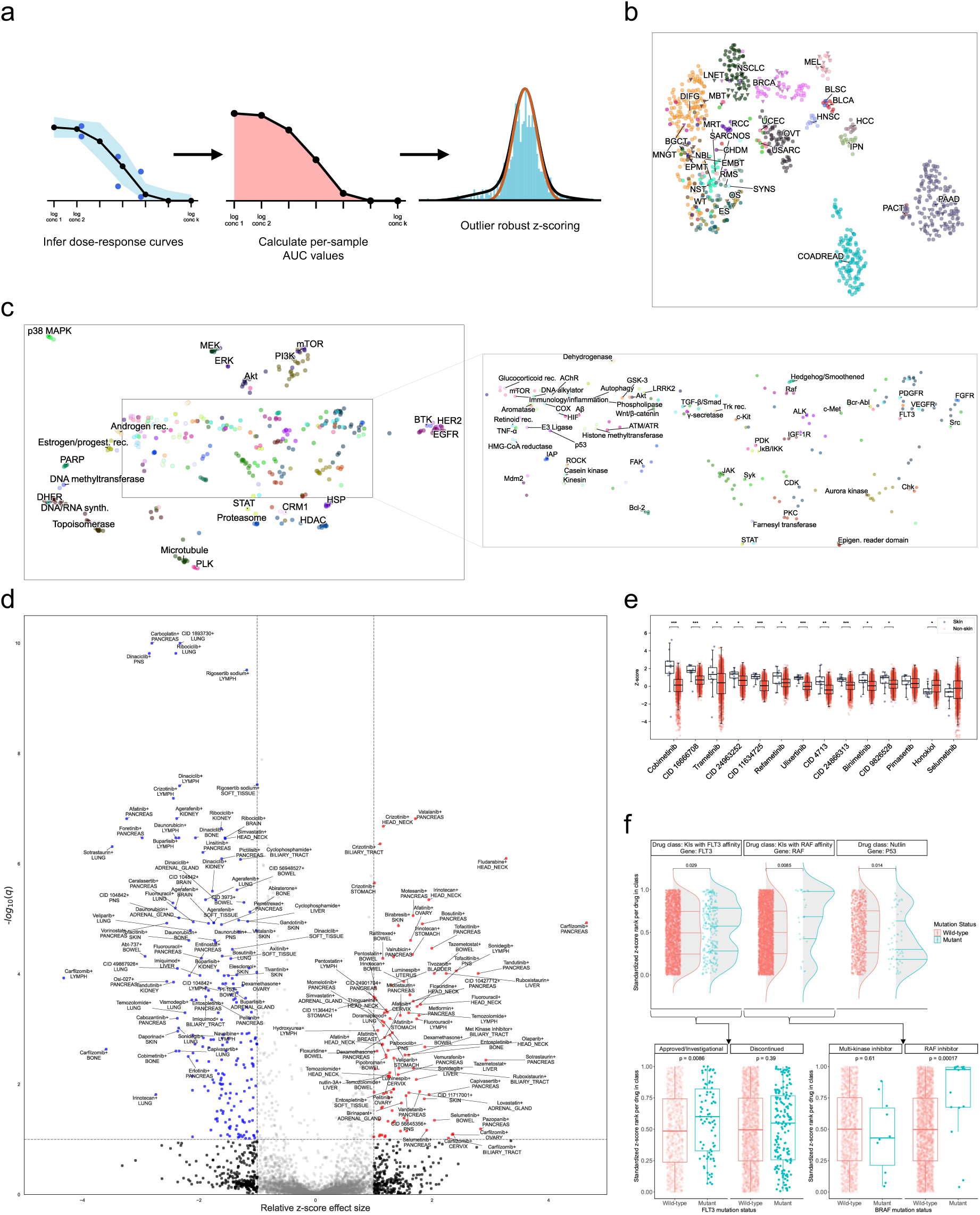
The PPC model learns biologically rational latent spaces and predicts clinically rational drug responses. (a) Batch effect-corrected z-scores are calculated on area under the dose-response curve (AUC) values using a robust null distribution procedure (see Methods). (b) UMAP of solid tumor sample embeddings colored by disease subtype. (c) UMAP of concatenated drug embeddings colored by most-common drug target. (d) Volcano plot showing top hits for primary site-grouped z-scores on each drug (x-axis: relative z-score difference, y-axis: negative *log*_10_ q-value from BH corrected t-test p-values). (e) Comparison of z-scores on drugs labeled as MEK/ERK inhibitors for both skin (left) and non-skin (right) primary tumor samples. (BH-corrected p-values from two-sided Mann-Whitney U tests, *: *p* < 0.1; **: *p* < 0.01; ***: *p* < 0.001. Boxes: first and third quartiles; line: median; whiskers: 1.5 × IQR). (f) Fully-imputed and standardized per drug z-score ranks for all drugs with given mechanisms of action in the Beat AML cohort. Top: Kinase inhibitors (KIs) with FLT3 and RAF affinity and Nutlin-derived drugs in samples with and without deleterious *FLT3*, *RAF*, and *TP53* mutations, respectively. Bottom, left: FLT3 KIs by stage of clinical development. Bottom, right: RAF KIs by kinase selectivity. (*p*-value calculated via two-sided Mann-Whitney U test; Boxes: first and third quartiles; line: median; whiskers: 1.5 × IQR).

## De-batched drug sensitivities show systematic differences across tissue sites

Assessing drug sensitivities across studies first required handling batch effects due to heterogeneity in experimental conditions between labs. The predicted dose-response curves, and corresponding AUCs, from the PPC model do not alleviate batch effects. The model seeks to minimize error for each sample in its specific study and thus recapitulates batch effects in the predicted response curves. In preliminary analyses, we identified that standard global z-scoring of AUCs fails to remove study-specific batch effects (fig. S15). To remove batch effects, we adapted an empirical Bayesian procedure^56,57^ to assign a de-batched z-score to each dose-response curve (Fig. 3a). For each sample, the procedure performs a robust estimation of the null distribution of the AUC histogram, ignoring outliers representing drugs for which the sample is truly resistant or sensitive which would skew the z-score distribution (see Methods). Grouping drugs by mechanism of action, we found global per-study batch effects are reduced using these robust z-scores compared to uncorrected AUCs (fig. S14). The empirical Bayes z-scoring procedure also produced z-scores that tended to be centered around zero, whereas global z-scoring leads to skewed median z-scores for each sample (fig. S17). Stratifying z-scores by drug target revealed broad differences in drug efficacy across disease types (fig. S19). Proteasome and IAP inhibitors are more cytotoxic than average aggregating across all tissue types. On the other hand, many traditional chemotherapeutic agents (e.g. DNA/RNA synthesis inhibitors like gemcitabine, 5-fluorouracil, and temozolomide) are less efficacious on average, possibly due partially to treatment-induced resistance in some samples. Regardless of the mean rank of a drug class, we observed considerable variance in average efficacy among disease types. For instance, proteasome inhibitors have the highest mean but a range of average z-scores from −2 (SARCNOS) to 3.3 (CEAD), whereas the lowest mean drug z-score across all disease subtypes is DHFR with an average z-score of −1.75. The large variability suggests that, while there is some toxicity bias to drug classes, substantial room remains for sample-specific sensitivities to emerge.

We hypothesized that using the de-batched z-scores as a measure of drug sensitivity may reveal systematic differences across tissue sites. We conducted a differential sensitivity analysis across the PPC dataset by filtering to primary samples and grouping by primary tissue. For each tissue site, we compared the average response to each drug to the average responses on the same drug aggregated across the other 22 annotated tissue sites with viabilities (see Methods). Overall, we found 303 tissue-specific drug sensitivities and resistances (Fig. 3d).

Several of the statistically significant sensitive drug-tissue type associations concord with findings from either molecular assays or clinical studies. VEGFR inhibitor vatalinib demonstrated favorable progression-free survival compared to historical controls in a Phase 2 trial in post-gemcitabine pancreatic adenocarcinoma patients^58^. Cytotoxic purine analog fludarabine has long been studied in head and neck cancer both as systemic monotherapy and in combination, either with radiotherapy or with fludarabine administered intratumorally^59–61^. Recent cell line and organoid screens identified transcriptionally defined sub-populations of pancreatic adenocarcinoma samples susceptible to proteasome inhibitor carfilzomib^62^. Thymidylate synthase inhibitor raltitrexed demonstrates comparable efficacy to 5-fluorouracil-based regimens in colorectal or bowel cancer^63^. EZH2 inhibitors promote cell death via ferroptosis in hepatocellular carcinoma^64^, and we find liver samples are significantly sensitive to EZH2 inhibitor tazemetostat. In combination with immunotherapy, tazemetostat has also shown disease control clinically in colorectal cancer^65^. Purine analog pentostatin shows sensitivity in lymph tissue, has been studied extensively in lymphoid malignancies, and is approved for use in the B-lymphocyte malignancy hairy cell leukemia^66,67^.

In general, drugs targeting the MAPK pathway, which includes MEK or ERK inhibitors, are more efficacious on primary samples from skin tumors than on samples from other sites in the PPC dataset (Fig. 3e). Drugs approved by the FDA for metastatic or unresectable BRAF V600E/K mutant melanoma (Cobimetinib, Trametinib, Binimetinib) were all significantly more effective in skin compared to non-skin samples^68–70^. In contrast, MEK inhibitor selumetinib is only approved for use in a rare, genetically predisposed peripheral nerve sheath tumor and is not significantly more effective in skin compared to non-skin samples^71,72^. Across drugs labeled as targeting MEK or ERK, z-scores representing drugs with measurements are well-mixed with those from fully imputed curves (fig. S18).

## Model imputations predict clinically rational drug responses

To assess the biological and clinical rationale of the model predictions, we conducted an analysis of the BeatAML^13^ study. We focused on BeatAML as it is unique among the PPC datasets for having a large sample size, a broad drug screen panel, and comprehensive genomic profiling for nearly every sample. Using the fully-imputed responses for AML, we identified samples amenable to treatment with agents for biomarker-defined sub-types, despite not explicitly incorporating genomic information in the PPC model (Fig. 3f, top). AML samples with deleterious mutations in *FLT3* and *RAF* genes are associated with improved responses to kinase inhibitors (KIs) with affinity for FLT3 and RAF, respectively, compared to their wild-type counterparts. This is consistent with the fact that FLT3 and RAF inhibitor approved indications are generally restricted to mutant patient populations^73,74^. In contrast, AML samples with deleterious *TP53* mutations were predicted to be less responsive to Nutlin-derived drugs. This underscores the mechanism of action of such drugs which abrogate the interaction between p53 and MDM2 and thereby reduce proteasomal degradation of the former by the latter; this has been demonstrated in wild-type constructs but may be more complicated in mutant *TP53* samples^75^.

Though FLT3 and RAF KIs were generally predicted to be more efficacious in *FLT3*- and *RAF*-mutant samples, respectively, the distributions of predicted z-scores are heterogeneous (Fig. 3f, top). In evaluating which drugs contributed to this phenomenon, we observed that approved or investigational FLT3 drugs (quizartinib, pacritinib, dovitinib) showed greater efficacy than those whose clinical development was discontinued (amuvatinib, tandutinib, CID 16041424, CID 11427553) (Fig. 3f, bottom left)^76–81^. Similarly, RAF KIs adjudicated by Selleckchem to be solely RAF-targeting agents were significantly more effective in *RAF*-mutant samples than those annotated as more broad-spectrum KIs with multiple kinase targets, including RAF (Fig. 3f, bottom right).

## Metastatic samples exhibit systematic differences in absolute and relative drug sensitivities from primary samples

The PPC dataset contains detailed site annotations for both primary and metastatic samples. The metastatic samples are annotated with both the primary site and the tissue site where the metastasis was sampled (Fig. 4a). None of the 26 studies in PPC performed longitudinal sampling of patients to allow for direct primary-to-metastasis comparisons within the same patient. However, the primary disease and site annotations enabled us to perform a population-level assessment of primary and metastatic samples that both originated from the same disease or site.

**Figure 4:**
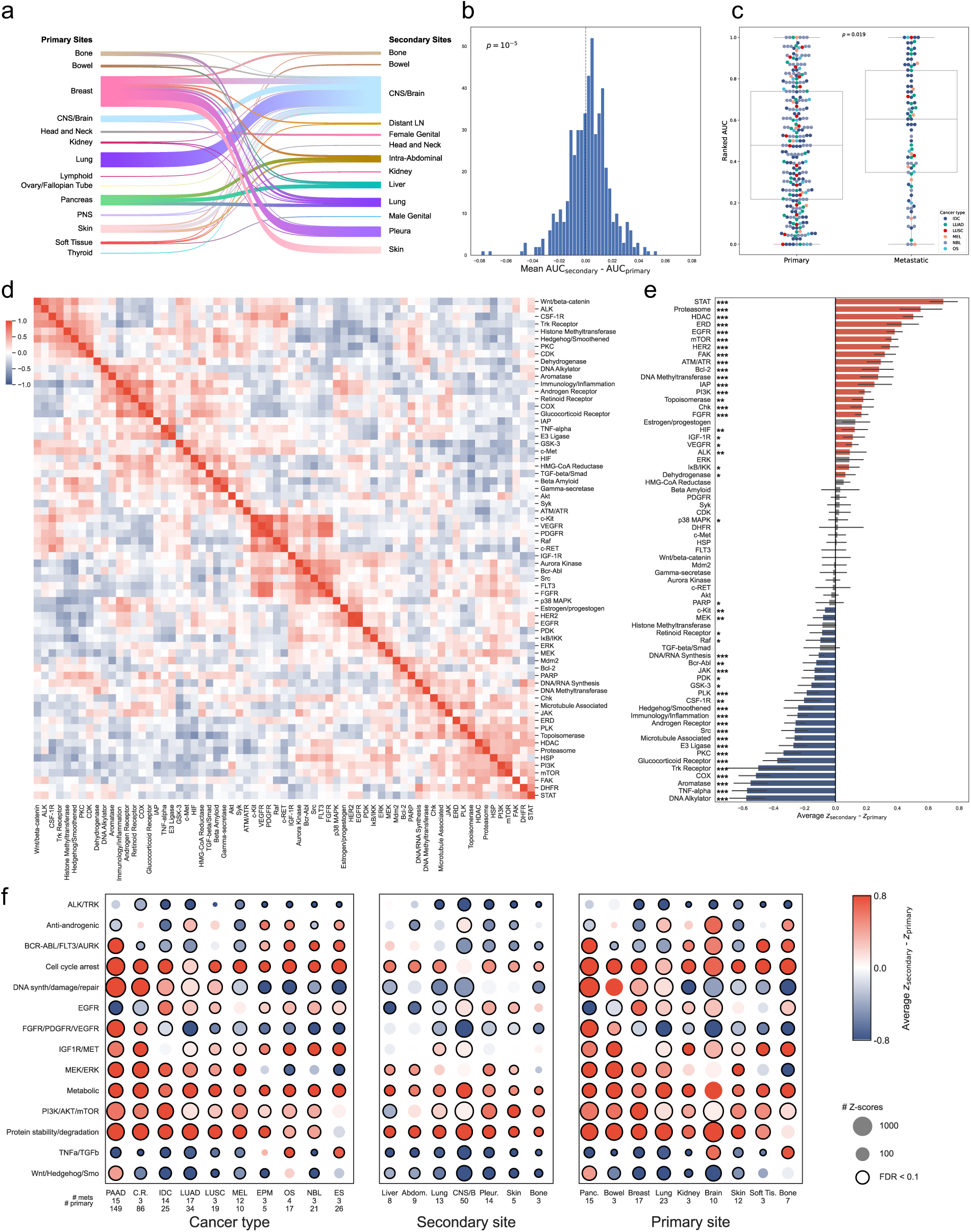
Metastatic samples possess differential drug sensitivities after correcting for intrinsic resistance. (a) Organotropism patterns reflected in the dataset. (b) Distribution of perdrug, per-disease differences in mean AUCs between primary and metastatic samples (left: metastasis more sensitive; right: metastasis more resistant; p-value: one-sided binomial test). (c) Restricting to per-disease standard-of-care drugs and ranking samples by AUC reveals that these drugs are on average less cytotoxic to metastatic ex vivo samples than to primary samples (Methods; *p*-value calculated via two-sided Mann-Whitney U test). (d) Correlations of differences in primary versus metastatic z-scores across disease types. (e) Average difference in z-score between primary and metastatic samples (left: metastasis more resistant; right: metastasis more sensitive; ∗∗∗: *q* < 0.001; ∗∗: *q* < 0.01; ∗: *q* < 0.1, Mann-Whitney U test on BH-corrected *q*-values; error bars: 90th percentile bootstrap intervals; red/blue: *q* < 0.1 and 95th/5th percentile has appropriate sign). (f) Grouped drug signature (vertical axis) z-score differences across top drugs for primary and metastatic samples, grouped by disease (left), primary site (center), and metastatic site (right). Color: mean z-score difference (red/higher: metastatic samples more sensitive; blue/lower: primary samples more sensitive; size: number of samples; solid outlines: significance at FDR=0.1; q-values: two-sided one-sample t-test against 0, with BH correction; C.R.: COADREAD; abbreviations: table S2).

Across all ex vivo samples of the same disease, metastatic samples were on average more resistant to the same cytotoxic agent (Fig. 4b, *p* = 1 × 10^−5^, one-sided binomial test). Metastatic samples remained more resistant on average when analysis was restricted to drugs designated as disease standard-of-care in the NCCN Guidelines (Fig. 4c, *p* = 0.019, two-sided Mann-Whitney U test)^82^. Drug resistance in metastatic disease is a well-described property of metastatic cancer^83^.

We reasoned that using the PPC z-scores instead of raw AUCs would remove the intrinsic resistance bias in metastatic samples and reveal patterns in relative drug sensitivity. Samples were stratified by disease type and primary status (primary or metastasis). In total, there were 20 disease groups with at least 1 sample in each stratum. Within each disease group, z-scores were averaged and the difference between the metastatic and primary average was used as a measure of relative increase in sensitivity for the metastatic samples (see Methods).

Hierarchical clustering on the z-score differences correlation matrix revealed functional subgroups of drug targets that see similar difference-in-response, across diseases (Fig. 4d). Targets upstream and downstream of one another in the same pathway, including PI3K/mTOR and MEK/ERK, are shown to cluster together. Several tyrosine kinase targets cluster closely in the center of the heatmap, including c-Kit, VEGFR, PDGFR, IGF-1R, Bcr-Abl, Src, and FLT3. We attributed this to the relative promiscuity of kinase inhibitors that leads single agents to target multiple kinases. A number of cell division-related targets cluster on the bottom right of the heatmap with emergent sub-clusters, including DNA damage repair and synthesis (PARP, DNA/RNA synthesis) and mitosis-related targets (Chk, microtubule associated, PLK, and topoisomerase).

Across diseases with both primary and metastatic ex vivo samples, comparative efficacy of labeled drug targets varies (Fig. 4e). Specifically, drugs labeled as targeting STAT, ATM/ATR, HER2, EGFR, FAK, HIF, and HDAC are found to have higher relative cytotoxicity on average in metastatic samples compared to primary samples across diseases. Conversely, drugs labeled as targeting TNF-*α*, DNA alkylators, and COX are found to be less relatively cytotoxic on average in metastatic samples compared to primary samples. Drugs labeled as targeting STAT were found to differ from those targeting JAK in comparative primary-metastatic sensitivity. JAK-targeting drugs were more effective on primary samples, while STAT-targeting drugs were more effective on metastatic samples. Differential phosphorylation of STAT3 has been previously observed between matched primary and metastatic lung cancer samples^84^. Targeting STAT3 activation has been observed to inhibit both tumor growth and metastasis in vitro and in vivo^85^, while targeting JAK alone has been observed to be insufficient in blocking growth of some late-stage ovarian cancer models^86^.

## Drug sensitivity differs based on metastatic sample site for some drug classes

Tissue- and disease-specific patterns of organotropism have been shown to be statistically associated with potential metastatic driver mutations^43^. Corresponding patterns in functional responses to drug treatments have not been investigated at scale. We stratified samples by primary and metastatic tissue sites, as well as disease subtype. Diseases and sites with fewer than three metastatic samples were excluded. Drug response z-scores were aggregated into higher level mechanistic groups based on the correlation structure among mechanism of action annotations (Fig. 4d). Differences between average z-scores were calculated and tested for significance using a two-sided t-test (see Methods). With some exceptions, average z-score differences between metastatic and primary samples are generally directionally consistent across cancer types, metastatic sites, and primary sites for the top drug classes (Fig. 4f). TNFa/TGFb targeting therapies are significantly more effective in metastatic compared to primary samples for OS and ES and for samples with primary sites in brain and bone, in line with preclinical reports regarding the importance of TGFb in local growth and metastatic progression in these tumor types^87–89^. Wnt/Hedgehog/Smo targeting therapies are significantly more effective in metastatic compared to primary samples for PAAD and samples whose primary site is the pancreas, possibly reflecting the importance of non-canonical Wnt signaling in pancreatic metastasis through effects of epithelial-to-mesenchymal (EMT) transition and cancer stemness^90^. Drugs targeting DNA synthesis, damage, and repair exhibited the greatest degree of variation across cancer types and sites. Such drugs exhibited significantly increased efficacy in metastatic compared to primary samples for PAAD, COADREAD, IDC, LUSC, and LUAD and significantly increased efficacy in primary compared to metastatic samples for MEL, EPM, OS, NBL, and ES. Overall, sample sizes for the subtype- and site-specific analyses were limited: only two site categories had more than 20 met samples and most cancer subtype categories had fewer than 15 mets. We therefore caution that these results in particular are preliminary and warrant expanded data collection to draw robust conclusions.

## Cell line drug responses are systematically different from ex vivo drug responses

The emergence of organoids and other ex vivo models has led to a shift in the drug screening field away from traditional immortalized cell lines and towards ex vivo models, in the expectation that results will better replicate patient responses to therapy^91^. Though one study in the PPC dataset^11^ performed limited comparisons to traditional 2D cell lines, no study to-date has had the breadth of samples and screen results to perform pan-cancer analyses comparing cell line and ex vivo drug responses. We sought to address this gap by integrating existing cell line drug screen atlases^24,92–101^ into the PPC dataset. We hypothesized that comparing drug responses between disease-matched model subpopulations would reveal distinct, recurrent differences between the two modalities.

Cell line drug responses were integrated into the PPC dataset using the same preprocessing pipeline previously described for ex vivo samples (see Methods). Primary diagnosis and metastasis status were obtained from Cellosaurus^102^ where available. In total, we integrated 2,790 cell lines, 21,005 drugs, and 50.3M viability measurements. We refit the PPC dose-response model jointly to all cell line and ex vivo samples. The same contrastive penalties were used in the joint model as in the model trained purely on ex vivo samples. Drug responses were normalized using the same robust z-scoring procedure used for ex vivo samples.

We first inspected the ability of the PPC dose-response model to learn integrated representations of the cell line samples. We found that cell line samples did not integrate with ex vivo samples when projected using UMAP (Fig. 5a). Classifying samples by disease type, we found that the sample embeddings on the jointly trained dose-response model qualitatively exhibit substantially more clustering by disease type than the cell line models (Fig. 5a). Comparing sample embeddings by nearest neighbors in Euclidean space confirmed that ex vivo models are significantly more likely than cell line samples to cluster with samples from the same disease type (Fig. 5b; Methods). We reasoned that immortalized cell lines may have lost many of the properties of their cell of origin and instead become driven by their molecular alterations due to selection for growth in culture. We found that cell line embeddings with TP53, KRAS, and BRAF mutations are significantly more like to cluster near other sample embeddings with the same mutation than wild type sample embeddings are (Fig. 5c), suggesting that mutation status of these genes is important in characterizing sample drug response. Similarly, cell line embeddings with high tumor mutational burden (TMB), as measured by the absolute number of mutations found, are significantly more likely to have neighbors with higher TMB (Fig. 5d). Using bootstrap data resampling, we observed that this trend was robust with an estimated 95% confidence interval of *r* = [0.225, 0.425] (fig. S22).

**Figure 5:**
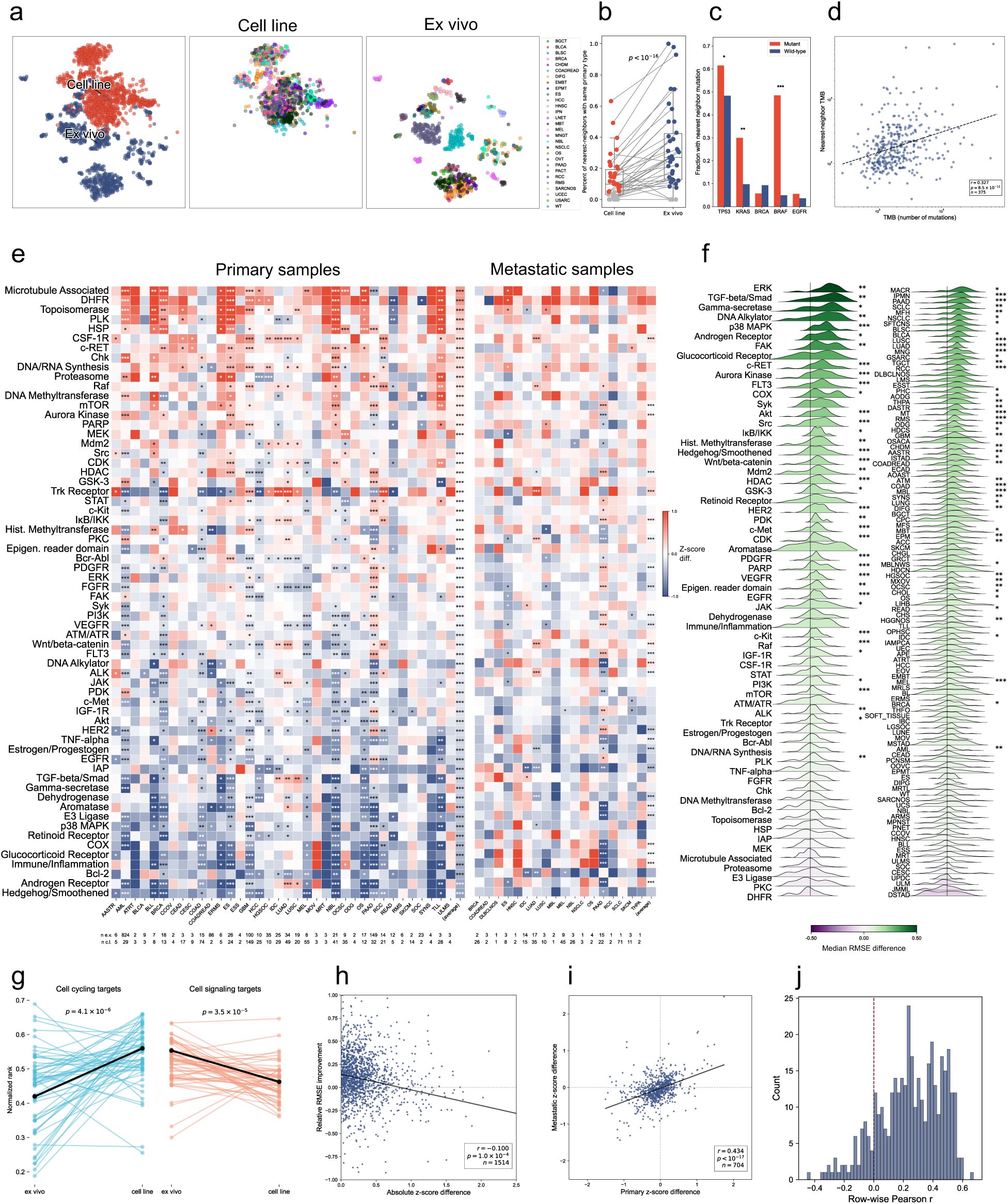
Cell line and ex vivo constructs differ systematically in drug response. (a) UMAP embeddings for cell line and ex vivo sample embeddings from the dose-response model trained jointly on all data (left). Ex vivo samples (right) cluster by primary cancer type, whereas cell lines (middle) do not. (b) Percentage of samples, grouped by disease, for which the aggregated 5 nearest-neighbors (in sample embedding space) is of the same disease. In general, ex vivo embeddings cluster more by disease than cell line embeddings; gray points: *q* > 0.1 after BH correction on a two-sided binomial test; global p-value via two-sided Fisher’s exact test (boxes: quartiles 1 and 3; line: median; whiskers: 1.5 IQR). (c) Proportion of cell line sample embeddings (with genomics) whose nearest-embedding neighbor has at least one mutation for that gene (*: *q* < 0.1; ***: *q* < 0.001; q-values: two-sided Fisher’s exact test with BH correction). (d) Tumor mutational burden (TMB) of samples nearest neighbor versus sample TMB, for cell line samples with genomics data (*r*: Pearson correlation; p-value: Pearson correlation test). (e) (Left) Relative efficacy differences stratified by disease types (horizontal axis) and annotated drug targets (vertical axis) for primary samples (left) and metastatic samples (right); red/higher: cell lines are more sensitive, blue/lower: ex vivo samples are more sensitive; *q*-values derived from BH-corrected permutation test (Methods); disease-average p-values calculated via Fisher’s method (∗∗∗: *q* < 0.001; ∗∗: *q* < 0.01; ∗: *q* < 0.1; BH corrected). (f) Distribution of per-disease relative change in model cross-validation RMSE when cell line data is added to model training set (Methods), stratified by drug target (left) and disease (right) (∗∗∗: *p* < 0.001; ∗∗: *p* < 0.01; ∗: *p* < 0.05); purple/lower: adding cell lines samples decreases performance (higher RMSE); green/higher: the addition increases performance. (g) Per-disease comparison of normalized rankings of cell-cycling and cell-signaling targeting drugs across model construct type. Higher rankings represent greater comparative drug efficacy on samples (p-values: two-tailed paired t-test; target groupings: table S4). (h) Relative RMSE improvement (as aggregated in (f) above) vs absolute z-score difference (as aggregated in (e) above) per target/disease pair (*r*: Pearson correlation; *p*: Pearson correlation test). (i) Primary versus metastatic z-score difference scores for matched diseases at least two samples of each group shows moderate concordance between primary and metastatic samples (*r*: Pearson correlation; *p*: Pearson correlation test). (j) Histogram of Pearson correlations per distinct pair of drugs across mean efficaciousness z-scores per-disease (fig. S25).

To assess whether cell lines have systematically different drug responses, we next compared z-scores between disease-matched model subpopulations. Cell line and ex vivo models were stratified by primary disease annotation and primary or metastatic sample status. Drug response z-scores were grouped by drug class and averaged to get the typical cell line and ex vivo relative efficacy scores for each class within a disease subgroup. The difference between the average cell line and ex vivo z-score were then calculated as a measure of deviation between modalities; results were tested for statistical significance via a nonparametric permutation test (see Methods).

Remarkably, we observed systematic deviations between cell line and ex vivo z-scores across disease types when grouping by drug class (Fig. 5e). Among 36 disease categories and 64 drug classes, we found 700 significant differences between disease-matched subpopulations (621 primary, 79 metastatic). Of these, 234 (33%) were more effective in cell lines and 466 (67%) were more effective in ex vivo samples. Across disease types, 63 of 64 drug classes were significantly different in primary samples and 24 classes were significantly different in metastatic samples, though only 5 exhibited differences greater than 0.5 z-score standard deviations (on primary samples, microtubule-associated, DHFR, and Hedgehog/Smoothened; on metastatic samples, DHFR and Bcl-2). Disease-matched differences were concordant between primary and metastatic indications (Fig. 5i,j), suggesting differences were robust to tumor stage.

Drugs targeting rapidly cycling cells (e.g., microtubule targeting drugs, DHFR inhibitors, topoisomerase inhibitors) were found to be significantly more effective on cell lines compared to ex vivo samples in 20/36 (56%) disease types. These include broad-spectrum chemotherapeutics like 5-FU, irinotecan, and oxaliplatin, which have previously been shown in limited experiments to be more resistant under 3D culture than in 2D monolayers^103^. Similarly, the chemotherapeutic Paclitaxel, which disrupts microtubule dynamics, has been found to be non-cytotoxic in some ex vivo models^104,105^ while inducing cell death in immortalized cell lines^106,107^. Aggregating across diseases, cell lines are significantly more sensitive to cell-cycling drugs compared to ex vivo samples and less comparatively sensitive to drugs targeting a broad class of intercellular signaling pathways (Fig. 5g). We observed a moderate to strong correlation (Pearson *r* = 0.434; *p* < 10^−1^^7^) between the primary and metastatic sample scores (Fig. 5i), suggesting the differences are robust to the stage of the underlying tumor, though more detailed clinical annotations would be needed to assess this systematically.

Drug classes that were relatively more effective on ex vivo samples were enriched for autocrine and paracrine signaling-related pathways. These include receptor tyrosine kinases (e.g., EGFR, IGF-1R, FGFR, VEGFR, PDGFR), migration and remodeling signaling (Wnt/beta-catenin, TGF*β*/Smad), and enzyme complexes (e.g., gamma-secretase, COX). Grouping by cell cycling or cell signaling related drugs showed significant trends for both polarized findings (fig. S20).

Among the less consistently polarized differences, TRK receptor drugs displayed outsized differences for either ex vivo or cell lines, depending on the specific disease subtype. Given the rarity of NTRK fusions, which occur at frequencies below 1% in common cancers like lung and colorectal cancer^108^, the relatively higher magnitude of differences across disease types for TRK receptor-targeting drugs observed in Fig. 5c is not explainable by on-target fusion activity. This may be due in part to the promiscuity of first-generation TRK inhibitors like entrectinib, which have been observed to inhibit multiple off-target kinases like ROS1 and ALK^109,110^.

We considered that differences in media conditions may have led to confounding that accounts for these results. To test this, we extracted all additives used in the media of the ex vivo and cell line studies (see Methods); this yielded 42 unique additives used in at least three studies. We one-hot encoded the presence or absence of each additive and whether the sample was a cell line or ex vivo sample, then ran two-covariate regressions for each drug target category and additive. Results showed that some additives do explain part of the variance in scores, but none affect the directionality or general systematic bias attributable to the differences between ex vivo and cell lines (fig. S23(a), fig. S24). We further tested this by running ℓ_1_-regularized multiple regressions controlling for all media additives, study ID, and ex vivo status. The high dimensional, collinear nature of the features led to high variation in the predicted coefficients, but we nonetheless observed the same trend as in the marginal and two-covariate analyses (fig. S23(b)).

## Integrating cell line drug screens improves ex vivo drug prediction accuracy

Despite the systematic differences between cell line and ex vivo drug responses, we found integration of cell lines still benefited predictive performance on ex vivo samples. Training the PPC model on cell line data combined with ex vivo data, compared with training on the ex vivo dataset alone, significantly improved empirical held-out prediction performance on ex vivo data for 38/64 (59%) drug classes and did not significantly reduce performance for any class (Fig. 5f, left). Similarly, model predictive performance improved significantly across 48/109 (44%) disease subtypes and did not significantly reduce performance for any class (Fig. 5f, right).

We observed a weak association between the size of cell line response discrepancy and integration improvement across drug classes (Fig. 5h). Leveraging bootstrap resampling, we observed that this trend was robust with an estimated 90% confidence interval of *r* = [−0.163, −0.079] (fig. S21). We further observed similar behavior across inter-drug correlation among both cell line and ex vivo samples (Fig. 5j). Drugs more often than not exhibit positively correlated behavior when comparing individual rows of the drug correlation matrices between ex vivo and cell line samples (fig. S25). These results suggest that, while cell lines are biased relative to ex vivo viabilities, the similar viability correlation structure appears to be sufficient to improve predictive ex vivo performance of the PPC model.

In aggregate, the results suggest that cell line drug responses should be interpreted and utilized with caution. Directly translating cell line responses to ex vivo responses is likely to be error prone and biased. However, when used as a data augmentation tool to better learn relational structure between drug responses, they consistently boosted the predictive performance of the PPC model. We therefore anticipate that future foundation models for ex vivo drug response would likely benefit from incorporating cell line drug responses.

## A Foundation Model for Functional Precision Oncology

Clinical application of functional precision oncology is often limited by low tissue availability and the need for biologically rational explanations of drug response^23^. Insufficient material for large scale ex vivo screens constrains the size and diversity of the drug library, reducing the range of possible therapies for patients. Molecular profiling also faces numerous clinical challenges^111^, making it difficult to generate orthogonal genomic and transcriptomic data needed to support molecular tumor board recommendations for drug screen hits. While notable successes exist, such as the INFORM trial^20^, it remains prohibitively challenging to scale simultaneous functional and molecular profiling across diverse cancer indications, patient cohorts, and disease stages.

We reasoned that a computational platform based on the PPC dataset could ameliorate these challenges. We sought to leverage the PPC database to build a foundation model for ex vivo drug screens that enables efficient and accurate “few-shot” prediction for new studies, using only a small number of drug response observations to predict a much larger range of doses and compounds. Unlike existing cell line dose-response models that assume comprehensive genomic and transcriptomic profiling^49,52,112–116^, we sought to build a model capable of predicting full ex vivo dose-response curves and molecular profiles entirely based on a small drug screen over a handful of concentrations. We also required that the model worked with novel patient samples, preventing the adaptation of cell line foundation models that rely on large language models to generate literature-informed embeddings about previously-studied samples^117^.

The PPC foundation model uses a transformer-based architecture inspired by BERT^118^ and TabPFN^119^ to perform few-shot multimodal prediction from a small number of drug screen results on a sample (Fig. 6a). The model is made of 8 transformer blocks with hidden layer size of 384, intermediate hidden size of 512 and 8 attention heads in each transformer. The foundation model directly takes as input a small labeled input set and an unlabeled query set, and outputs predictions for the query samples. The input set consists of a small number of labeled triplets (drug, dose, viability) measured on a new biological sample, while the query set contains unlabeled pairs (drug^′^, dose^′^) for which the model predicts viability. This setup allows the model to infer viability responses for unseen drug–dose combinations based solely on a few measured examples from the same biological sample. Both input and query sets are encoded using sequence of tokens for the drug, dose and viability. The drug tokens are given by an embedding layer, which is a dictionary retuning a vector of size 128 for each drug. For the dose and viability embedding, we also use vectors of size 128. To encode the continuity of these values, the embedding is given by a linear combination of Fourier features with learnable parameters. Finally, the three embeddings corresponding to the drug, dose, and viability are concatenated to form a 384-dimensional embedded token for each (drug, dose, viability) triplet in the input sequence. For the query sequence, a placeholder token is used for the unknown viability value, resulting in query tokens with the same 384-dimensional representation. The input and query sequences are then processed by eight Transformer blocks, which perform cross-attention between the two sequences. The resulting query token representations are passed through a linear prediction head to estimate viability.

**Figure 6:**
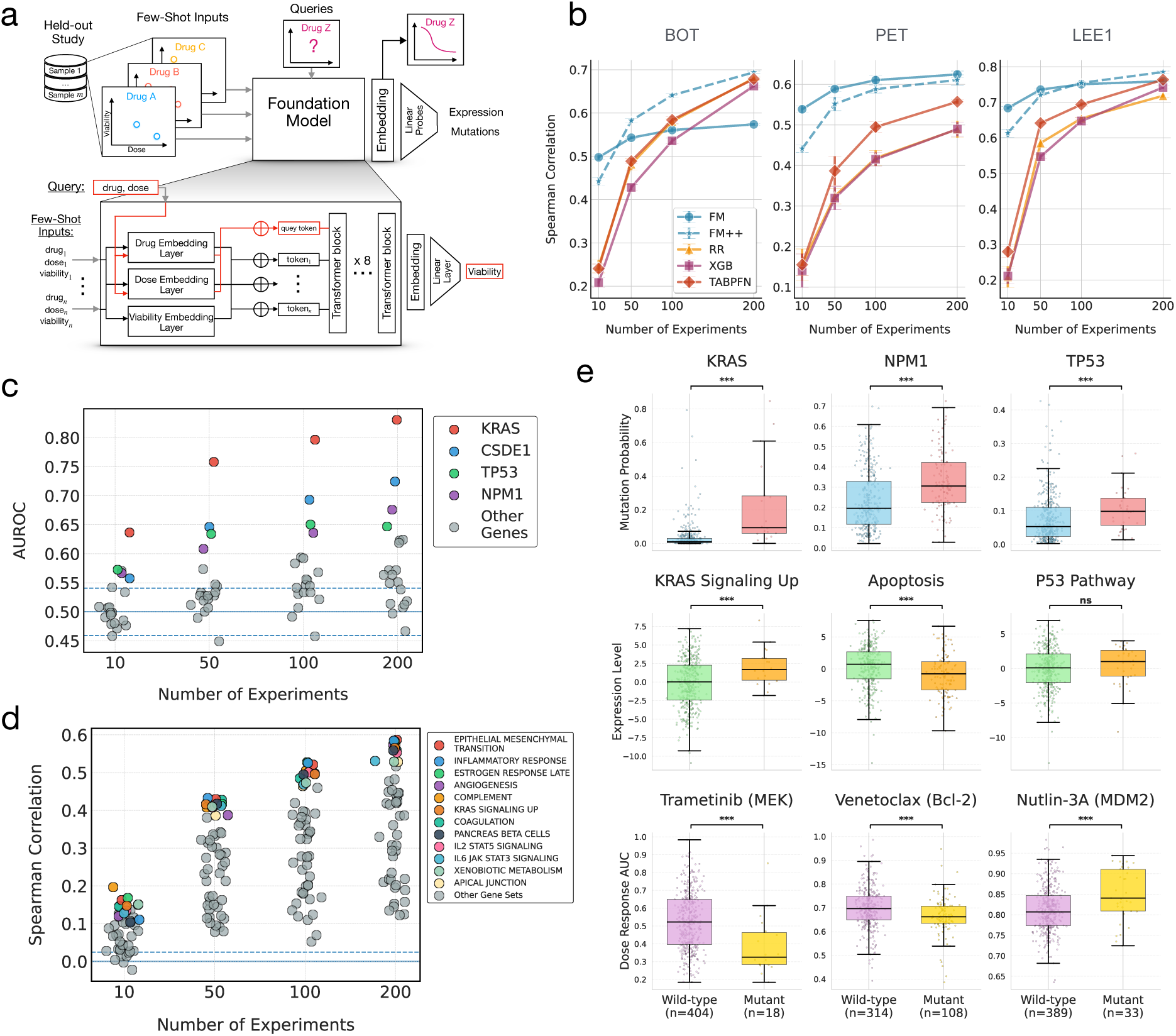
Foundation model evaluation and multi-omics analysis. **(a)** Overview of the experimental setup. The foundation model (FM) is trained on PPC with one study held out at a time, enabling few-shot dose–response inference for unseen samples. **(b)** Few-shot drug response performance. Spearman Correlation for dose response prediction as a function of the number of experiments, i.e. the number of (drug, dose, viability) observations (*n*_few-shots_ ∈ {10, 50, 100, 200}). The three held-out studies are BOT ^13,28^, PET ^20^ and LEE1^9^. Error bars show two standard deviations over five random repetitions. The Foundation Model (FM) and its finetuned version (FM++) is compared against TabPFN, XGBoost (XGB), and Ridge regression (RR) baselines. **(c–e)** Multi-Omics analysis conducted on the Beat AML cohort (BOT). **(c)** Prediction of gene mutation status from sample embeddings. Linear probes are trained to predict binary mutation labels using embeddings derived from the FM. Shown are mean AUROCs across five cross-validation folds and five random few-shot selections, highlighting the 4 most accurately predicted genes. **(d)** Prediction of pathway-level gene expression. Ridge regression models predict aggregated Hallmark pathway activity scores from FM embeddings. Results are averaged over five cross-validation folds and five random few-shot selections; the 12 best-predicted pathways are displayed. **(e)** Multi-omics consistency analysis across mutation, pathway activity, and drug response. Rows show mutation logits, pathway scores, and predicted drug AUCs for *n*_few-shots_ = 100. Boxplots compare wild-type and mutant groups. One-sided Mann–Whitney U tests were repeated 5 times; p-values were combined using Fisher’s method. Significance: * p < 0.05, ** p < 0.01, *** p < 0.001.

## The PPC foundation model reduces the number of experiments needed to accurately predict drug responses

To assess few-shot performance, we evaluated how accurately the model can recover the dose–response for a new sample in a held-out study when only a few observations are available. We conducted independent benchmark experiments using the three PPC studies (BOT, PET, LEE1). The three studies chosen represent the largest drug libraries, broadest range of concentrations, and the largest ex vivo sample sizes, enabling us to test a range of potential down-scaled experimental designs. Each benchmark experiment used a leave-one-study-out design, holding out one study at a time while training on the rest of the PPC dataset, including cell lines. The held-out study was then treated as a new domain, where only a few-shot subset of labeled observations is available. These few-shot examples were used as input to the foundation model to produce dose–response curve predictions and sample embeddings.

We evaluated the viability predictions and embeddings to assess the capacity of the model to generalize to unseen studies and efficiently adapt to new drug screen experiments. Performance on viability predictions was compared against linear (ridge regression), machine learning (XGBoost^120^), and generalist foundation models (TabPFN^119^). For the PPC foundation model, we considered two use cases: (i) a static model (FM) that makes predictions purely using attention over the inputs, and (ii) a fine-tuned model (FM++) that updates the model weights to the new data before making predictions.

The PPC foundation model consistently outperformed all baseline models across the three held-out studies (Fig. 6b). Baseline models approached FM performance when the number of experiments was large (≥ 200). When data was more limited, as might the case with low tissue availability, the PPC foundation model correlation was 100-400% higher than the baselines. Fine-tuning (FM++) slightly underperformed FM when only 10 experiments are available but achieved similar results after 50 experiments for two studies. For one of the three studies, the fine-tuned model provided a clear improvement, suggesting that this study is either more challenging or less similar to the other PPC studies. We therefore generally recommend fine-tuning when possible to ensure the most robust performance. When access to GPUs are unavailable, the static foundation model still represents a high performing model available for rapid prediction.

## Foundation model probes provide rational genomic and transcriptomic interpretability for drug predictions

A key benefit of foundation model architectures is their ability to generate embeddings that serve as general purpose covariates for predicting a wide range of related tasks for which the model was not originally trained^121^. We reasoned that the PPC foundation model embeddings may enable accurate imputation of genomic and transcriptomic profiles, particularly those driving phenotypic response to drugs. If accurate, imputing molecular profiles would enable a layer of biological interpretability to the foundation model predictions. Imputed profiles could also serve as biological rationale for molecular tumor boards in the setting where comprehensive multimodal profiling is infeasible.

To evaluate the embeddings, we used the Beat AML cohort (study BOT), which gathered both whole exome sequencing and bulk RNA sequencing on matched patient tissues, in addition to performing ex vivo drug screens. For each sample, the foundation model produced embeddings derived from dose–response predictions using *n*_few-shots_ ∈ {10, 50, 100, 200}. We then trained simple linear probes on these embeddings to assess how well they captured underlying molecular features. Binary mutation status was predicted using logistic regression. Gene expression data was normalized and log-transformed, then aggregated at the pathway level using the Hallmark gene sets^122^; the first principal component across member genes was used as the pathway activity score. Performance on each modality was evaluated using 5-fold cross-validation.

Prediction accuracy varied across targets but was particularly high for targets directly related to drug response and resistance (Fig. 6c-d). Mutations in key oncogenic and resistance drivers, namely KRAS, TP53, CSDE1 and NPM1 were among the most accurately inferred (Fig. 6c) with AUROC 0.65-0.83 with 200 observations. Drugs in the Beat AML library, like idasanutlin and trametinib, target upstream or downstream targets of these genes. In the case of KRAS, this structure is further reflected in the embedding space (fig. S26), where mutant and wild-type samples form distinct clusters. Pathway prediction showed variability but consistently improved in accuracy as the number of observations increased (Fig. 6d). Pathway predictions were also most accurate when they related to disease severity (e.g. epithelial mesenchymal transition) or were connected to drug targets in the panel (e.g. KRAS signaling up).

We further analyzed the molecular imputations for cross-modality sensitivity. While each modality may be well predicted in isolation, it is possible that the two predictions were discordant when considered jointly, or in context of the predicted drug response. For each selected gene–drug pair, we jointly examine the mutation probability inferred from the foundation model embeddings, the corresponding pathway activity score, and the predicted drug sensitivity (AUC). The results show coherent molecular patterns across omics layers and drug response predictions (Fig. 6e). In KRAS-mutated samples, the model assigns higher mutation probabilities and elevated scores for the “KRAS signaling up” pathway, while predicting increased sensitivity to Trametinib–a MEK inhibitor acting downstream of KRAS. Similarly, NPM1-mutated samples show higher predicted mutation probabilities and reduced activity of the apoptosis pathway, with Venetoclax (a BCL-2 inhibitor) predicted as more effective in this subgroup. For TP53, the model correctly associates higher mutation probabilities with reduced sensitivity to Nutlin-3A, an MDM2 inhibitor whose efficacy depends on intact p53 signaling. Altogether, these results indicate that the foundation model embeddings capture biologically consistent associations between mutations, transcriptional programs, and pharmacological responses.

## Discussion

This work has established and analyzed a pan-cancer ex vivo drug screen atlas. In doing so, it addresses outstanding questions in translational cancer research, including (i) is multi-study drug response integration valuable in the presence of batch effects and differing experimental conditions, (ii) how do metastatic and primary samples differ in drug response, and (iii) in which ways do cell line drug responses differ from their ex vivo counterparts. The comparative analyses and modeling results provide guidance for future study designs with respect to data integration strategies and power calculations. This work also established the pan-preclinical (PPC) project and foundation model, computational resources for the cancer biology and functional precision oncology communities.

Several key limitations underlie the PPC dataset and represent important challenges in the development of next-generation methods for predicting ex vivo drug response. First, the degree of matched omics varied widely between studies, making the overall PPC dataset highly sparse in side information. This sparsity led us to design a flexible tensor factorization method rather than building on existing machine learning methods for drug response prediction, which typically rely on matched genomics and/or transcriptomics^49–52^. More consistently matched omics would enable a deeper dive into mechanistic drivers that could explain, for instance, many of the differences we found between cell lines and ex vivo drug response. Similarly, the PPC samples were taken as-is from studies with different selection criteria. Some studies chose to obtain only treatment naive samples, whereas others took heavily pre-treated patients. As we were unable to retrieve complete treatment history for each sample in the dataset, it is difficult in principle to disentangle the inter-related effects of highly progressed cancer, metastatic cancer, and prior anti-neoplastic treatment on drug responses observed across samples. There are also many careful considerations one must make when interpreting drug screen data based on cell viability;^123^ for example, bulk metabolic assays cannot readily distinguish between proliferative arrest and subpopulation-level cell death, among other concerns. Gathering additional time points to estimate growth rate inhibition^124^ or directly tracking single cells or organoids in real time^125^ would alleviate these confounding issues.

Ex vivo screening platforms are advancing at a rapid rate, with newer assays enabling more faithful modeling of the tumor microenvironment (TME). We anticipate that emerging ex vivo assays that incorporate TME characteristics (e.g. through co-culture^126^, microfluidics^127,128^, or explant culture^129,130^) represent the next generation of ex vivo platforms. Tumor cell interactions with other cells in the TME, such as cancer-associated fibroblasts^131,132^, cytotoxic T-cells^133^, regulatory T-cells^134^, and M1/M2-like macrophages^135^, are known to mediate tumor proliferation^136^ and response to some therapies^137^. These platforms could also enable in vitro testing of drugs targeting the TME and its constituent cell types such as PD-1/PD-L1/CTLA-4 blockade for exhausted T-cells and emerging therapies like anti-CSF1/CSF1R therapy for tumor associated macrophages^138^. The PPC project is built flexibly and with an eye towards incorporating a wide array of studies and measurements. As these technologies mature, we anticipate incorporating both TME assays and therapies into the PPC dataset to improve its translational value and reveal more insights into tumor drug resistance and response. The PPC project is therefore poised to continue to push the boundaries of translational cancer biology and predictive oncology.

## Acknowledgments

We thank Maurice Markus for his assistance in adjudicating sample metadata curation decisions. We also thank all of the authors of the PPC component studies that worked with us to share the raw version of their published data and to retrieve clinical annotations, including Ameen Salahudeen, Alexandria Bobe, Florin Selaru, Ling Li, David Tuveson, Herve Tiriac, Hans Clevers, Else Driehuis, Gerald Schwank, Suet Yi Leung, Michael Shen, Sato Toshiro, Kohta Toshimitsu, Alana Welm, Seung-Won Choi, Raul Rabadan, Sven Nelander, Alejandra Bruna, Helen Piwnica-Worms, Reid Powell, Olli Kallioniemi, Astrid Murumägi, Sean McAllister, James Brenton, Filipe Correia Martins, Glenn Marshall, Loretta Lau, Emmy Dolman, Ina Oehme, Heike Peterziel, Gustave Ronteix, Jean Bouteiller, Fanny Jaulin, and Alice Soragni. We are also grateful to the companies and organizations that further assisted in data gathering, including Tempus, Orakl Oncology (for assistance with organoid data funded originally by grant ANR-21-RHUS-0003), and the TARGET/ZERO team at the Children’s Cancer Institute in Australia.

## Author contributions

Manuscript drafting: K.P., J.B.W., J.Q., A.M., C.T., A.D.M., F.H., W.T. Manuscript editing: W.T., K.P., J.B.W., J.Q. K.P. performed general data analyses and wrote much of the software to run and evaluate the code. J.B.W. analyzed expression and genomic data J.Q. developed the data processing, harmonization methodology and software, and the dataset website. A.M. curated and harmonized sample metadata. C.T. formulated and coded the model ultimately used in the manuscript. A.D.M. led development and analysis of the foundation model. E.C. analyzed efficacy of standard-of-care treatments on primary and metastatic samples. F.H. performed power analysis to estimate the model’s effectiveness at identifying unseen drugs. W.T. supervised the research and performed data analyses.

## Data and materials availability

Unless excepted below, the data that support the findings of this study will be made freely available. The data from Mayoh et al. ^29^ is available upon request from the authors of that study. Restrictions may apply to the availability of these data, which were used with agreement for this study.

## Supplementary materials

## Materials and Methods

### Dose-response harmonization

We defined a dose-response observation as a single experiment where a given cancer model was exposed to one or more drugs at defined concentrations. A single dose-response observation must be labeled with a harmonized sample identifier, and one or more harmonized drug identifiers. Drug concentration across all studies was converted to micromolar units (*μM*). All dose-response observations must have a negative control where the same model was exposed to no drug, and optionally can have a positive control where the model was exposed to a compound at a known lethal concentration. We calculate viability as

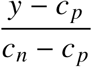

where *y* is the raw readout of the experiment (e.g. fluorescence value, culture area, etc) and *c*_*n*_ and *c*_*p*_are the raw readouts of the negative and positive controls, respectively. In the case where the positive control is not included in the study it takes the value 0 in this formula. In cases where *y* was observed multiple times within the same technical replicate (e.g. if the same model is exposed to the same concentration of drug in multiple wells on the same plate) the mean of those observations is used. The raw readout units vary between studies, but because the viability metric is always normalized by control values measured in the same units it is comparable across studies. Viability is a positive real number, with a value greater than 1.0 indicating the model grew more under drug treatment than under the negative control.

In some source studies raw experiment readouts were not reported and viability metrics were reported directly; in these cases we confirmed that the reported viability metric will be comparable to metrics we calculated directly using the above formula.

### Cancer primary annotation

Cancer diagnoses were harmonized to the OncoTree nomenclature^42^ by matching sample diagnoses to NCIT records’ synonym/abbreviation fields^139^ in addition to Cellosaurus records’ disease field^102^. The combined list of NCIT synonyms in addition to Cellosaurus diseases were then iteratively queried on the OncoTree API^42^, until the most specific OncoTree code that was a direct match was identified. For example, a sample from Lee et al. ^9^ diagnosed as “Myxoid liposarcoma, FNCLCC grade 1/3” simplified to “Myxoid liposarcoma” returns no direct OncoTree code matches^42^. However, the combined results from querying NCI ThesaurusSioutos et al. ^139^ and CellosaurusBairoch ^102^ yield [“MLPS”, “Myxoid Liposarcoma”, “Myxoid/Round Cell Liposarcoma”, “Myxoliposarcoma”], of which the third entry, “Myxoid/Round Cell Liposarcoma”, maps directly to the OncoTree code of “MRLS”. Difficulties were posed by the free text nature of most cancer diagnoses data collection and the lack of standardized acronyms. As a result, when no direct matches could be found by this method, manual adjudication was required to select the most accurate OncoTree code. In a few cases of extremely rare disease with no appropriate match, the OncoTree code was marked as null or the primary tissue type.

### Cancer metastasis annotation

Cancer metastasis sites were harmonized under the MSK-MET classification system^43^. Only distant metastases, that is, neoplasms sufficient to categorize as “M” within the TNM guidelines set by the AJCC^140^ for each cancer type, were considered. For rare cancers and central nervous system cancers (e.g. meningiomas and medulloblastomas) for which TNM staging was nonexistent, the original study’s judgment as to metastasis status was used. Due to the non-standardized nature of metastasis description, manual adjudication was utilized when there was not a direct match to MSK-MET classes in order to select the most accurate metastasis site label.

### Media additive annotation

For all but three studies, detailed media additives were described or referenced in the source publications ^15,28,98^. We noted the presence or absence of specific media additives, including growth factors, small molecules, enzyme degraders, amino acids, antibiotics, antimycotics, antioxidants, proteins, vitamins, serums, and fatty acids. For studies that referenced protocols from previous work rather than detailing their own methods, additives were assumed to be identical to the cited publications. We did not include basal media, which were generally Dulbecco’s Modified Eagle’s Medium/Ham’s F-12 (DMEM/F12) or Roswell Park Memorial Institute (RPMI) media. Additives used in each study were assumed to be applied to all samples, though sample-specific differences were noted in some studies. A total of 50 media additives were documented, 42 of which were used in three or more studies and included in our analyses. Full names for additive abbreviations and acronyms are available in table S6.

### Genomics variant harmonization and annotation

Genomic variants were reported several ways in the source publications; standards such as HGVS Variant Nomenclature and VCF files as well as a variety of *ad hoc* nomenclatures and tabular formats were observed ^141^.

We developed a data model to encapsulate the heterogeneous variant information reported by our source publications and created a software framework to extract and transform all variants into this model. Once extracted, variants were harmonized and enriched using GenomeNexus^142^. GenomeNexus is able to harmonize variants from different reference sequence modalities; for example a variant reported using genomic coordinates like 7:g.55249071C>T can be matched to variants reported using amino acid changes like EGFRp.T790M. We further used GenomeNexus to enrich our annotated variants with information such as transcript consequences, SIFT and Polyphen scores, and clinical information from OncoKB ^143–145^. Frameshift and stop-gain mutations along with missense mutations with deleterious SIFT predictions were annotated as deleterious. Once annotated, all variants were lifted over to the GRCh38 reference genome to facilitate cross-publication analysis ^146^.

### Expression harmonization

Upon review of the relevant publications, we noted that 8 studies performed RNA-seq experiments as part of their protocols and made the resulting raw or normalized count matrices readily publicly accessible^1,4,9,11,30,31,37,41^. Upon request, two authors provided raw FASTQ files from their RNA-seq experiments for analysis^7,34^. One study performed concurrent microarray profiling and made the corresponding quantile-normalized results readily publicly accessible^38^.

For the datasets that required FASTQ file processing, we created a Nextflow pipeline available in a standalone repository1 to enable reproducible quality control, adapter trimming, sequence alignment, read quantification, and initial gene identifier harmonization^147^. The FASTQ files were first processed for quality control using FastQC (v0.12.1), and any identified adapter or poly-G or poly-A tail contamination was resolved using fastp (v0.23.4)^148,149^. The resulting sequences were aligned using STAR (v2.7.11b) to the original 2013 hg38 “soft-masked” reference genome indexed with the GENCODE knownGene transcript model (last updated 2023-06-28), both of which were downloaded from the UCSC Genome Browser^150–152^. The resulting BAM files were indexed using SAMtools^153^.

Once the internally processed or author-provided count matrices were ingested, the resulting gene identifiers were either HUGO Gene Nomenclature Committee (HGNC) gene names or ENSEMBL gene IDs. We used pyensembl^154^ and Ensemble release 108 to map all identifiers to HGNC gene names and combined any resulting multi-mapping transcripts. We also converted sample identifiers to harmonized PPC identifiers to allow for integration with viability and clinical data. Finally, we normalized the resulting count matrices to counts per million to account for differences in library size.

### Drug harmonization

All drugs were matched to the PubChem database utilizing the PubChem Power User Gateway (PUG-REST)^155^, a restful API. We queried the PubChem database based on the primary name of the drug used in the study. PubChem has two separate databases: the Substance database stores depositor-contributed information, while the Compound database contains on entry per unique standardized chemical structure^156^. Because of this we preferentially annotate our drugs with entries from the Compound database, but will use the Substance database if no match to the Compound database is found We used two methods to match study drug names to PubChem; exact name match and search index match. Exact name match only returns an entry if the search term exactly matches that entry’s name. The search index match is equivalent to typing a keyword into the PubChem web application search bar and selecting the best result.

When available in the database entry, we retrieved the SMILES string^157^ for each drug. We also retrieved the parent compound for the entry when available, which is the primary organic component of the compound^158^. If two drugs had the same parent compound, or one drug is the parent of another, we merged them as one drug in our data model. For example, tetracycline hydrochloride (CID 54704426) and tetracycline metaphosphate (CID 54729668) were merged, as they share the same parent compound, tetracycline. Our algorithm for merging two drug entities is given in algorithm S1.

### Drug target annotation

We assessed the feasibility of obtaining drug target annotations from several open-source resources, including DrugBank, OpenTargets, PubChem, and chemical supplier Selleck Chem^158–160^. To select a single resource to use, we manually compared annotated targets of kinase inhibitors, which often inhibit a broad spectrum of such proteins due to their structural and sequence conservation^161^.

table S3 compares the drug target annotations for these four resources to published kinase dissociation constants for multi-kinase inhibitor agerafenib. Only the targets identified by Selleck Chem are either the stated target the compound was designed to inhibit, RAF, or kinases on which agerafenib demonstrated single-digit nanomolar potency using a standard, pan-kinome in vitro screening assay^162^. This finding is consistent with our broader review and led us to use target annotations curated from Selleck Chem.

We attempted to match all drugs in our dataset to the World Health Organization’s International Non-Proprietary Name (INN) list by using a partial match of the INN to the PubChem primary name and any synonymous names available under the PubChem substance or compound entry followed by a manual review ^163,164^. We obtained drug mechanisms of action from the chemical supplier Selleck Chemicals, which provides drug target annotations for many of the compounds in its libraries. We included Selleck’s L1100 inhibitor, L1300 FDA approved drug, and L1700 bioactive libraries, which contain 4,945, 3,067, and 9,125 compounds, respectively.

We then the queried the INN, PubChem name, and PubChem synonyms against the Selleck database on partial matches of the Selleck product name to the primary name or INN, or an exact match to any synonyms. During each query step, matches were reviewed manually to remove duplicates, disambiguate multi-mapping matches, and harmonize drug targets that differed slightly between the various libraries. For example, the non-inhibitor libraries included non-molecular targets that capture secondary effects of the drugs, such as “apoptosis related” or “autophagy,” and were removed from all but drugs that directly target these pathways as their primary mechanism of action (e.g., elesclomol/apoptosis, chloroquine/autophagy). Once one associated name was annotated with a mechanism, no further identifiers were queried for that entry. For matches adjudicated as accurate, we retained all Selleck-provided targets.

### Harmonizing immortalized cancer cell lines

In order to ensure the same immortalized cancer cell lines used in different studies were consistently identified, we queried all immortalized cell line identifiers against the Cellosaurus database using their REST API^102^. Cell lines which were matched against the database were annotated with their Research Resource Identifier (RRID)^165^. Two cell lines in different studies annotated with the same RRID in this way were considered to be same for the purposes of our downstream analysis. We used version 8.4 of GDSC1 and GDSC2.

### Annotating immortalized cancer cell lines

We leveraged metadata in the Cellosaurus database to assign cancer diagnoses and metastatic status and sites to immortalized cell lines. Cellosaurus entries for cancer cell lines are often annotated with an National Cancer Institute Thesaurus (NCIt) ontology code^139^. OncoTree entries are annotated with synonymous NCIt codes, allowing us to assign OncoTree codes to these cell lines. Cellosaurus entries are also annotated with an indicator as to whether the cell line was established from metastatic tissue, and an UBERON ontology code indicating the anatomical site where the tissue was taken from^166^. We constructed a mapping of UBERON ontology terms to the MSK-MET classifications enabling us to annotate the cell lines with MSK-MET categories.

### Non-parametric Bayesian tensor factorization model

Our model utilizes discrete grids over log-concentrations *c*_1_ < *c*_2_ < · · · < *c*_*n*_ as well as viabilities 0 = *y*_1_ < *y*_2_ < · · · < *y*_*n*_ = 1, where the assumption is that all observed concentrations fall within [*c*_1_, *c*_*n*_] and all viabilities fall within [0, 1]. An observed viability *y* of a sample *i* treated with drug *j* at concentration *c* is modeled as

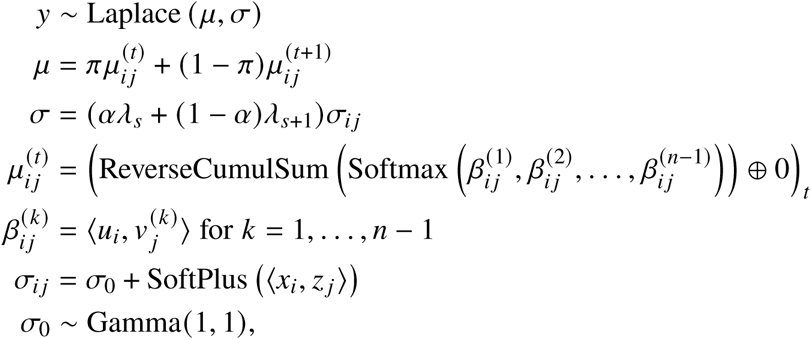

where *π* ∈ [0, 1] and *t* ∈ {1,…, *n*} are the unique values satisfying *c* = *πc*_*t*_ + (1 − *π*)*c*_*t*+1_, and similarly, *α* ∈ [0, 1] and *t* ∈ {1,…, *n*} are the unique values satisfying *μ* = *αy*_*t*_ + (1 − *α*) *y*_*t*+1_. ⟨*u*, *v*⟩ denotes the inner product between vectors *u* and *v*. The softmax function is the vector-valued function

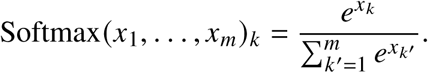

The reverse cumulative sum function is the vector-valued function

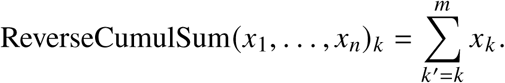

The softplus function is the real-valued function satisfying

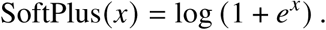

We have further used the notation *v* ⊕ *x* to denote concatenation of vectors and scalars. *u*_*i*_ and 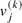 are *p*_1_-dimensional embeddings generated as

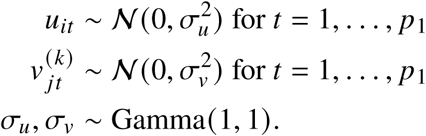

Similarly, *x*_*i*_ and *z* _*j*_ are *p*_2_-dimensional embeddings generated according to

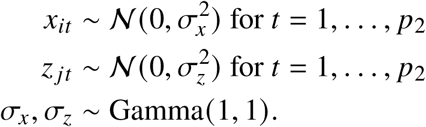

The vector *λ* ∈ R^*n*^, which non-parametrically shrinks the predicted variance as a function of the predicted mean satisfies

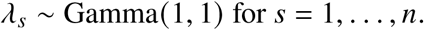

In addition to optimizing the ELBO loss on the observed viabilities, we optimize two contrastive losses over the mean sample embeddings, *u*_*i*_, and mean drug-grid embeddings, 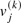. Specifically, for the sample contrastive loss, we sample an index *i*, an index *i*^pos^ corresponding to the same primary site as *i*, and a set of indices 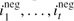 corresponding to primary sites that differ from *i*. The contrastive loss is given by

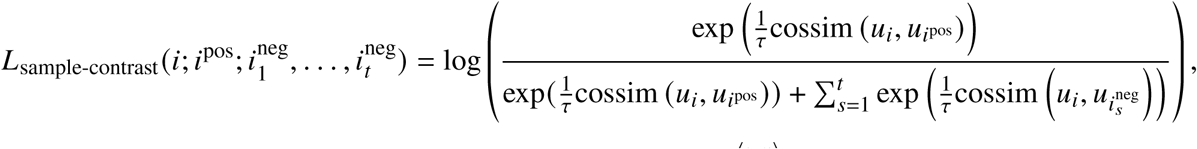

where *τ* > 0 is a temperature parameter, and cossim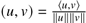 is cosine similarity. Similarly, the drug contrastive loss at grid-point *k* is computed by sampling an index *j*, an index *j*^pos^ corresponding to a drug with a mechanism that is shared with *j*’s mechanisms, and a set of indices 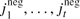 corresponding to drugs whose mechanisms do not overlap with those of *j*.

Then the contrastive loss is given by

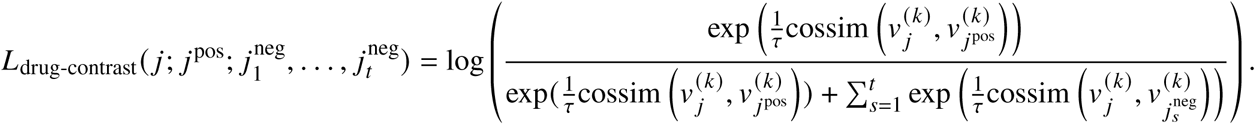

### Preprocessing and training

We restricted the corpus to experiments using drugs appearing in three or more studies, resulting in a vocabulary of 504 drugs. We clipped all viabilities between 0 and 1.

We applied the following preprocessing steps to filter out noisy experimental data. First, we filtered out any measurements for which there exist multiple experimental replicate points at the same dose which are ≥ 0.5 apart in measured viability space (clipped between 0 and 1). Second, given the downward monotonicity of the model, we restricted to curves that have a Spearman *ρ* correlation of ≤ 0.25; that is, we discarded dose-response measurements whose measurements trend strongly upwards as drug dose increases. Finally, we discarded experiments with contiguous measurements (in concentration space) which increase in magnitude at least 0.5 in measured viability space, using sudden upward jumps in viability as a proxy for high experimental noise. Due to low concordance with other datasets, we omitted Johansson et al. ^10^ from our model training and analysis. Dataset sizes throughout filtering steps is given in fig. S5.

All training runs were conducted with the following hyperparameters: *p*_1_ = 150, *p*_2_ = 5, *n* = 25, and *τ* = 0.1. We set *t*, the number of negative sampled contrastive examples, to 10. We trained using stochastic variational inference (SVI)^167^ for 500 epochs with a batch size of 50,000 using Adam^168^ with a learning rate of 5 × 10^−3^.

### Calculating z-scores

For each (sample, drug) pair, our model produces a monotone piecewise linear curve estimating the mean dose-response at each concentration value. We first calculated the Area Under the Curve (AUC) of each dose-response curve by numerically integrating the dose-response curve via the trapezoidal rule from −9.2 log_10_ *μM* to 4.0 log_10_ *μM* in log-concentration space. Since each sample has its own intrinsic baseline sensitivity level, we normalize AUCs by calculating a robust z-score per sample. To do this, we fit a second order polynomial to the AUCs for a given sample across drugs. We then matched this polynomial to a Gaussian distribution using a second order Taylor expansion. This provides a mean and variance estimate for a Gaussian null distribution for drug effects on the sample. Individual drugs were then converted to z-scores using the inverse cumulative distribution function of the Gaussian. This approach was adapted from the literature on empirical Bayes hypothesis testing^56^. Unlike simple standardization (i.e. subtracting the mean and dividing by the standard deviation), this empirical null approach has the advantage of not allowing large effect sizes to skew the mean. In cases of samples’ second-order polynomial being convex (comprising 2.1% of samples), preventing the match from polynomial to second-order Taylor approximation of a Gaussian, we fell back to simple standardization

### Model evaluation

All data-holdout evaluation results are given for a whole-drug holdout five-fold cross-validation setup: each study’s drugs were partitioned into five sets of equal cardinality (modulo differences from rounding), with a single fold’s validation set comprising one such set for each constituent dataset in the corpus. Folds were generated such that no drug occurs in a validation set without being in the corresponding training set for at least some other study; that is, within a cross-validation fold, every drug in a validation set must have some training data from at least one study. This data-holdout setup probes the empirical performance of predicting full dose-response curves for (drug, sample) pairs on which the drug has not been observed on any samples from a given study, but both the sample and drug have some data in the training set.

The per-dataset-pair marginal utility calculations given in Fig. 2b were derived by comparing root mean-squared error (RMSE) on a scored evaluation dataset with and without a held-out dataset. That is, we first ran five-fold cross-validation with the full dataset, using the whole-drug holdout discipline as described above, evaluating the extent to which the model and dataset allow generalization to drugs not observed in a study. Then, for a held-out dataset ℎ, we ran an additional five-fold cross-validation run omitting ℎ from each of the five folds, but otherwise using the same folds so results are comparable. We then compared empirical performance on each probe dataset *p* both with and without held-out dataset ℎ. In particular, we evaluated predictions on *p* using RMSE and calculated *p*-values with a one-sided binomial test, using the Benjamini–Hochberg procedure for false discovery rate adjustment.

The per-disease-type stratified Pearson *r* results in Fig. 2c and Fig. 2d were calculated in the same whole-drug-holdout five-fold cross-validation setup described above. In general a single drug may contribute to multiple groupings in Fig. 2d, as drugs in general have multiple mechanisms labeled. All Pearson *r* correlations were calculated at the level of individual viability point measurements in the whole-drug-holdout cross-validation setting.

### Power analysis

Most drugs in the PPC dataset were tested on a small subset of samples; conversely, most samples only test a small subset of drugs. By accurately modeling drug response, we can prioritize top *in silico* drugs for downstream validation, potentially enabling drug re-purposing and personalized treatment. Therefore, we investigated the model’s power through the lens of nominating the most efficacious drug for a given sample via imputation.

We subsetted the PPC atlas to create a dataset with a high evidence level, which we call (PPC-strict). We select studies in which 1) at least 10 samples were tested, 2) at least 60 drugs were tested, and 3) ≥ 90% of (sample, drug) pairs were tested on at least 3 drug concentrations. PPC-strict contains 1331 unique samples and 495 drugs, which are sourced from 11 studies: Driehuis et al. ^4^, Lee et al. ^9^, Sa et al. ^14^, Murumägi et al. ^16^, Peterziel et al. ^20^, Malani et al. ^26^, Pemovska et al. ^27^, Bottomly et al. ^28^, Powell et al. ^37^, Bruna et al. ^38^, Lau et al. ^41^ There is no sample overlap between any of the studies.

We simulated a 2-stage drug selection campaign on a set of unseen samples, where the goal was to discover the most potent drug for each sample as defined by the lowest half-maximal inhibitory concentration (IC50) as given by the model. In the first stage, *k* drugs were selected and experimentally validated on the new samples to establish a baseline. In the second stage, there was only budget to experimentally validate *n* additional drugs. Instead of handpicking the *n* drugs, we used data from the first stage to rank all drugs in the PPC atlas *in silico* and experimentally validated the top-ranked drugs.

To simulate such a campaign, we pre-trained separate PPC models on 10 studies in a hold-one-out fashion and performed drug selection on the held out study. In the first stage, we randomly selected *k* drugs from the held out study for observation and retrained the model. We then evaluated whether the PPC model could identify the most efficacious drug (lowest IC50) within the *n* drugs it nominated. Results are given in Fig. 2e.

### Baseline models and comparisons

Baseline model comparisons are given in Fig. 2fgh. The *Bucketed mean* baseline estimator is a global dose-level mean curve estimate: that is, we bucketed into 20 equally-spaced buckets (in log-concentration space), from the lowest tested concentration in the dataset to the highest; within each bucket, we calculated the global mean across all curves and inferred that mean for all points in the bucket. This baseline provides a comparison to a global notion of the dataset’s average dose-response curve.

The random forest baseline represents an ensemble of 100 random decision trees trained on bootstrap-resampled instances of the dataset; within a decision tree, intermediate nodes are required to have at least 10, 000 measurements to be split (approximately 1% of a fold’s training set). The random forest is trained to regress to viability by minimizing ℓ_2_ loss.

The neural network baseline model represents a simple multilayer feed-forward network neural network. Each drug and sample are assigned embeddings *e*_*d*_ and *e*_*t*_, respectively; concentrations are bucketed into 20 equally-spaced buckets (in log-concentration space), from the lowest tested concentration in the dataset to the highest, and a measurement at a concentration is given an embedding *e*_*c*_ for its concentration bucket. The model is then FFN(*e*_*d*_ ⊕ *e*_*t*_ ⊕ *e*_*c*_), with ⊕ representing concatenation, and FFN being a sequence of alternating fully-connected affine transformations (with intermediate hidden layers of dimension ℎ) and rectified-linear units, followed by a final fully-connected affine transform into a single scalar viability clipped to [0, 1]. The neural net was optimized via Adam^168^ minimizing ℓ_2_ loss. Within a fold, hyperparameters were selected via a random search (evaluated on 20% of that fold’s training set) and the setting with optimal RMSE out of 10 settings was chosen. Batch size was selected from {8, 16, 32, 64, 128, 256, 512, 1024}; the embedding dimension of *e*_*d*_, *e*_*t*_, and *e*_*c*_ (with all taking the same dimension in a given network) was selected from {16, 32, 64, 128}; the hidden width ℎ was selected from {32, 64, 128, 256, 512, 1024}; the number of hidden layers was selected from {1, 2, 3, 4}; the number of training epochs was chosen from {1, 3, 5, 10, 15, 20}; the Adam learning rate was selected from {1 × 10^−3^, 5 × 10^−4^, 1 × 10^−4^, 5 × 10^−5^, 1 × 10^−5^ }. After a fold’s hyperparameters are selected, an MLP is trained on the fold’s full training set using that setting.

All comparisons are done in a curve-holdout five-fold cross-validation setting. That is, we partition the dataset into five roughly-equal-sized subsets such that each (drug,sample) pair’s measurements is entirely contained within a single fold.

### Low-dimensional projections of sample and drug embeddings

The UMAP plots in Fig. 3b and Fig. 3c were generated from the sample embeddings *u*_*i*_ and concatenated drug embeddings *v*^(*k*)^. Drug embeddings in Fig. 3c were annotated with the Selleck-provided mechanism for the drug; drugs with multiple mechanisms are labeled with the most common mechanism across the dataset.

### Quantifying drug-site effects

To generate Fig. 3d, primary (non-metastatic) samples were grouped by site as annotated in the OncoTree hierarchy. Drugs most heavily comprising pure imputations were filtered out by restricting to drugs with at least 1,000 individual measurements across the dataset. Primary sites with fewer than 5 distinct samples were filtered out. We applied the empirical Bayes methodology described above (Methods) to each resulting group of z-scores, grouped per drug and per site and used Benjamini-Hochberg correction to derive q-values. Relative z-scores are given per-target; that is, each quantity represents the difference between the experimental group’s mean z-score compared to the full mean z-score for that target. We restrict to solid tumors and filter out pancreas samples due to the sparsity of measurements for the site in the dataset.

To compare skin vs. non-skin samples in Fig. 3e, we restricted to drugs annotated with any targets in groupings in table S5. Healthy and non-primary (metastatic) samples were filtered out. Comparisons were taken between all remaining skin and non-skin samples for each drug annotated as targeting MEK or ERK.

The Beat Acute Myelogenous Leukemia (AML) cohort provided the most robust genomic annotations with 576 of the 629 samples (91.6%) for which we predicted drug z-scores possessing documented alterations. To assess the extent to which our model, which did not explicitly include genomic information, identified samples with clinically actionable alterations among Beat AML samples, we standardized fully-imputed z-score ranks by drug for agents with biomarker-defined mechanisms of action (Fig. 3f, top).

### Comparing metastatic and primary samples

Mean AUC differences between metastatic and primary samples given in Fig. 4b were calculated per-drug per-disease; that is, the distribution of effect sizes was calculated by iterating over the full OncoTree hierarchy and, at each node, if that disease has both metastatic and primary samples represented, we calculated the difference in mean dose-response AUC for each drug. We averaged across all diseases for each drug and calculated per-drug mean AUC difference between primary and metastatic samples across diseases. The population-level *p*-value was calculated via a one-sided binomial test.

To compare metastatic vs primary sample resistance to per disease standard of care drugs in Fig. 4c, the 2024 NCCN Treatment by Cancer Type was referenced for each OncoTree code. Only diseases with at least three metastatic samples were included in this analysis. Rhabdomyosarcoma (RMS) and Ewing Sarcoma (ES) were excluded from comparisons as the disease subtypes with the lowest predictive performance in the subset of the dataset used for comparison (Fig. 2c). Treatments ranked as “Preferred” or “Other Recommended” for a disease were split into their individual drugs, and corresponding drugs (if present) in the PPC database were labeled as as standard of care. For each sample, the AUCs for each standard of care drug were extracted. Many NCCN treatment guidelines for a disease consist of multiple drugs, for example “Weekly Paclitaxel and Carboplatin.” In such cases, The AUCs corresponding to each sample/drug/dose triplet were summed. The processed AUCs for all standard of care treatment regimens for a given OncoTree code were concatenated and rank normalized. A two-sided permutation test was applied comparing the PIC50s corresponding to metastatic samples treated with standard of care regimens and all primary samples treated with standard of care regimens.

To calculate per-drug correlations in Fig. 4d, we restricted analysis to non-brain metastases, diseases with both primary and metastatic samples in the dataset, and drug target annotations with at least three drugs represented in the dataset. We aggregated at the level of disease-metastatic-site pair; that is, for each drug, for each primary disease type (as given by OncoTree annotation), and for each metastatic site (as described above), we calculated the difference in mean AUC-based z-score between the metastatic samples and primary samples of the same disease. Each mechanism was thus represented as a *k*-element vector, with *k* the number of distinct (disease, metastatic-site) pairs present in the filtered dataset. Finally, we scaled this collection of *k*-element vectors into z-scores following the same method the AUC-based z-score calculation described above, transforming into z-scores at the (disease, metastatic-site) pair level. Fig. 4d gives the Pearson product-moment correlation coefficients between all pairs of *k*-element z-score vectors representing target annotations. Fig. 4e gives per-mechanism primary/metastatic z-score differences at the disease level. For each disease type (most-granular OncoTree annotation) with at least two metastatic samples in the dataset, the mean difference in AUC-based z-scores between primary and metastatic samples was calculated. Fig. 4e depicts 90% uncertainty bars as given by a bootstrap resampling estimate. Mechanism-level *p*-values were calculated via two-sided Mann-Whitney U test with Benjamini-Hochberg correction. Fig. 4f gives detailed information on drug response on metastatic samples compared to primary samples. Drug target annotations were grouped into coarse classes following the covariance structure given in Fig. 4d and labeled based on shared mechanisms; full mechanism groupings are given in table S5. The leftmost panel of Fig. 4f gives differences in drug response between metastatic and primary samples grouped by primary OncoTree type. Differences in z-scores were calculated for diseases with at least 3 metastatic and at least 3 primary samples present; *p*-values were given by a two-sided permutation test at the sample level and a Benjamini-Hochberg correction was applied. For the middle and right panel of Fig. 4f, each metastatic sample was compared to the mean z-score of the corresponding primary samples of that disease type; that is, for each drug, for each metastatic sample of disease type *t*, we compared the AUC-based z-score of that drug on that sample to the mean z-score of that drug on all primary samples of type *t*. Fig. 4f gives the mean of all such differences, with *p*-values calculated via two-sided permutation test at the sample level. A Benjamini-Hochberg correction was applied to derive *q*-values.

### Comparing ex vivo and cell line clustering

Fig. 5a, which depicts the UMAP projection of cell line and ex vivo samples for the dose-response model trained jointly on both types of data, shows qualitatively that ex vivo sample embeddings cluster more cleanly by disease type than than cell line sample embeddings To quantify this observation (Fig. 5b), we calculated the percentage of sample embeddings’ neighborhoods with the same disease type as the sample. All samples with an OncoTree label with 5 or fewer samples were relabeled to the disease type one level up in the ontology’s hierarchy; this process was iterated until each disease type has at least six samples. We then discarded any samples from diseases not represented in both ex vivo and cell line samples. We calculated each sample’s 5 nearest neighbors by Euclidean distance in the sample embedding space. Samples were aggregated by disease type and, for each disease, we calculated the percentage of the union of its samples’ neighborhoods that have the same disease label. To calculate per-disease significance levels (represented as colors in Fig. 5b), we applied Benjamini-Hochberg correction to *p*-values calculated via two-sided binomial tests with null probability equal to the dataset-level disease prevalence.

### Quantifying cell line sample-neighborhood genomic patterns

The populations used to calculate the fraction of nearest neighbor samples with different mutations in Fig. 5c were restricted to samples from studies with genomics. BRCA1 and BRCA2 were combined into a single category “BRCA.” Tumor mutation burden (TMB) in Fig. 5d was calculated as the raw number of mutations called for a sample.

### Comparing ex vivo and cell line cytotoxicity

Comparative curve-level efficacy across disease types and drug mechanisms is given in Fig. 5e. We calculated response difference for all drug target annotations that are assigned to at least 3 drugs across the model’s inventory of drugs. Samples were partitioned by most granular OncoTree level, with metastatic samples grouped separately from primary samples. Each (disease, mechanism) pair comprises a set of measurements for ex vivo samples and one for cell-line samples. The comparative efficacy of a mechanism’s drugs on a disease (comparing between ex vivo and cell line model types) is given by the mean AUC-based z-score across all drugs annotated for that mechanism on all relevant samples. To calculate *p*-values for each group, we applied a two-sided permutation test, permuting sample labels 10, 000 times and measuring the percentage of permutations with absolute effect size at least as extreme as the observed. To combine per-disease *p*-values into aggregate mechanism-level *p*-values, we used Fisher’s method; aggregate mechanism-level effect sizes are given by macro-averaged disease-specific effect sizes.

### Measuring the effect of cell line data on ex vivo model performance

We quantified the effect of adding cell line data to empirical predictions on held-out ex vivo data Fig. 5f. As above, five-fold cross-validation of ex vivo performance was performed at the drug level; that is, a given constituent study’s drugs is partitioned into five roughly-equal-sized sets with one set held out for each of the five folds’ train runs (that is, a drug in a fold’s evaluation set was not observed on the set’s samples during train). Folds were constructed so that no evaluation drug is fully absent from the train set. We ran ten train/evaluation folds in total—on the one hand, the five-fold validation run only on the ex vivo data; on the other hand, the same five splits but with the full corpus of cell line measurements over the model’s drugs added to the training set. In all cases, the evaluation sets comprised only ex vivo measurements.

Density-estimate curves in Fig. 5f are Gaussian kernel-smoothed estimations on a per-disease basis. For each disease (i.e. the most granular OncoTree annotation), we calculated ex vivo samples’ held-out root-mean squared error (RMSE) across folds. Letting *e*_*d*_ denote disease *d*’s RMSE across all measurements on all relevant samples in the ex vivo-only cross-validation setup, and *e*^′^ denote the RMSE on the cross-validation run which includes cell line data in train, Fig. 5f gives the per-mechanism distributions of relative error differences (*e*_*d*_ − *e*^′^*_d_*)/*e*_*d*_ across diseases *d*. Per-group *q*-values were calculated via applying Benjamini-Hochberg correction to *p*-values calculated via two-sided binomial tests on per-disease binarized improvement indicators (i.e. binary variables taking value 1 if and only if empirical RMSE is lower with cell line data added to train), with null binomial *p* = 0.5.

### Comparing cycling and signaling targeting drugs across cell lines and ex vivo samples

To compare comparative efficacy of drugs targeting cell cycling and cell signaling pathways broadly construed in Fig. 5g, we restrict to primary samples and we use the grouping of targets given in table S4. For each drug, samples are ranked by their z-score sensitivity and ranks are normalized to [0, 1], with higher values indicating higher efficaciousness/sensitivity. Samples are grouped by OncoTree, and only OncoTrees with at least 3 ex vivo and 3 cell line samples are included. The individual lines connect median normalized rank in the corresponding sample set, with the per-model-construct aggregate given by the median of such values.

### Comparing ex vivo RMSE improvement and construct-type sensitivity

To compare relative RMSE improvement to mean z-score difference between ex vivo and cell line models in Fig. 5h, we calculate RMSE improvement for every (disease-indication, mechanism-of-action) pair with observed data, treating primary and metastatic cancers as different subclasses. We then compare to the mean z-score difference as given in Fig. 5e.

### Comparing primary and metastatic z-score differences between ex vivo and cell line samples

To compare the difference in z-score between primary and samples as given in Fig. 5i, we restrict to OncoTree codes that have at least 2 primary samples a and 2 metastatic samples in both cell line and ex vivo datasets and directly compare average z-score discrepancies.

### Comparing drug-drug similarity across ex vivo and cell lines

To quantify the extent to which drugs behave similarly across cell line and ex vivo samples as given in Fig. 5j, we first calculate drug-drug covariance across diseases (fig. S25), where each entry is the Pearson’s product-moment correlation coefficient across per-OncoTree mean z-scores on that drug. We calculate one covariance matrix for cell line samples and one for ex vivo samples (using the same dimensions so they are directly comparable). We then, for each row index *i*, calculate the Pearson *r* of the *i*th row of the two individual covariance matrices, giving us a quantification of how similarly the drug corresponding to that row behaves across the two model construct types.

### Comparing additive effects across ex vivo and cell lines

To assess the potentially confounding effects of media differences, we annotated 50 media additives as previously described. For the 42 additives that occurred in three or more studies, we performed independent two-covariate linear regressions predicting the average drug target z-scores per sample using one-hot encoded variables indicating the presence or absence of a given additive and whether or not the sample was a cell line or ex vivo construct and including an intercept term (fig. S23(a), fig. S24). Hierarchical agglomerative clustering was performed on the media additives to identify groups with similar coefficient profiles across drug targets using Ward’s minimum variance method and Euclidean distance. Given the highly collinear nature of additives, construct type, and study ID, we performed ℓ_1_-regularized multiple regressions predicting the the average drug target z-scores per sample using one-hot encoding for these covariates and an intercept term (fig. S23(b)). We used five-fold cross-validation to determine an optimal regularization parameter and bootstrap resampling to determine a standard error of the mean of the ex vivo coefficient. To assess concordance between the ordering of the negative one-hot encoded ex vivo coefficient by drug target from this analysis and the ordering of deviations between cell line and ex vivo z-scores by drug target as shown in Fig. 5(e), we computed the Spearman correlation coefficient and its corresponding p-value between these two sets.

### Foundation Model Design

#### Architecture

We use a BERT^118^–like architecture with 8 transformer blocks, hidden layer size of 384, intermediate hidden size of 512 and 8 attention heads in each transformer. The foundation model is designed for few-shot learning on structured data: rather than requiring gradient-based optimization at inference time, it directly takes as input a small labeled input set and an unlabeled query set, and outputs predictions for the query samples. This formulation enables efficient adaptation to new tasks with very few examples, without any model parameter updates or tuning of optimization hyperparameters.

#### Model Inputs

In our application, each model input corresponds to a specific biological sample (PDO, PDC, or PDCL) for which viability measurements are available under various drug and dose conditions. The input is structured as paired input and query sets. The input set consists of a small number of labeled triplets (drug, dose, viability), while the query set contains unlabeled pairs (drug^′^, dose^′^) for which the model predicts viability. This setup allows the model to infer viability responses for unseen drug–dose combinations based solely on a few measured examples from the same biological sample.

Formally, denoting the model by *f*, the labeled input set by S_input_ of size *n* ∈ N, and the unlabeled query set by S_query_ of size *m* ∈ N, the predicted viabilities for the query set satisfy:

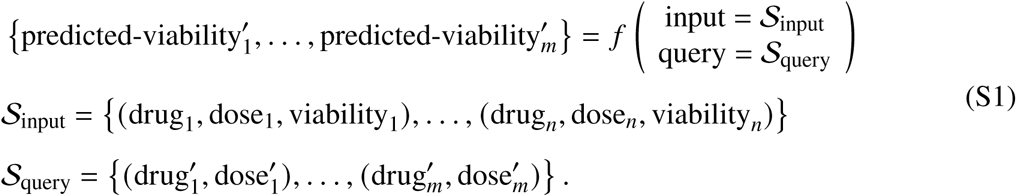

Both input and query sets are encoded using sequence of tokens for the drug, dose and viability. The drug tokens are given by an embedding layer, which is a dictionary retuning a vector of size 128 for each drug. For the dose and viability embedding, we also use vector of size 128. To encode the continuity of these values, the embedding is given by a linear combination of Fourier features with learnable parameters matrix *A* ∈ R^128×128^ and vector *w* ∈ R^128^ and input the scalar “dose” normalized in [0, 1] such that: dose-embedding

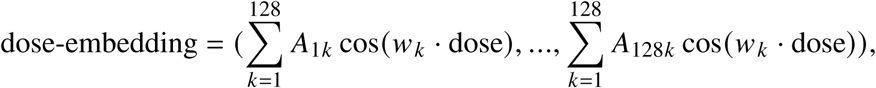

The same type of embedding is used for viability. Note that dose refers to the min-max scaled log-concentration. Finally the three embeddings are concatenated to form the embedded token of size 384 for each (drug, dose, viability) triplet in the input and query sequence.

#### Training on PPC

During training, cross-attention is performed between the tokens of the input and query sequences. No self-attention is performed between tokens of the same sequence. A linear head is used on top of the final embedding of each query token to predict the viability of the unlabeled (drug, dose) query pairs. To provide a sample embedding we average the final token embedding over the query sequence. ℓ_1_ loss is used between predicted and observed viabilities.

The training dataset consists of approximately *n*_sample_ ≃ 5000 splits of the PPC dataset, each corresponding to the (drug, dose, viability) measurements of a single biological sample. One training epoch comprises 100,000 gradient update steps, each iterating over randomly selected samples. At each step, the measurements of the selected sample are randomly divided into two parts of random size: the labeled input set S_input_ and the unlabeled query set S_query_. The true viability values of the query set are held out and used to compute the loss between them and the predicted viabilities obtained according to Equation S1. The model is trained on 200 epochs with learning rates 1*e*−5 using the Adam optimizer^168^.

#### Inference on New Samples

At inference time, the model operates in the same few-shot fashion as during training. Given a new biological sample with a small set of observed (drug, dose, viability) measurements (input set) and a set of query conditions (query set), the model directly predicts the viabilities of the query conditions without any further gradient updates. This property makes inference both computationally efficient and convenient, as no optimization parameters (e.g., learning rate, batch size) need to be specified.

### Foundation Model Results

#### Experimental Setup (**Fig. 6b**)

We train three foundation models (FMs), each trained with a different study held out. For each held-out study, all samples from that study are removed from the

PPC dataset during training. This setup simulates a realistic use case scenario in which a practitioner aims to infer the dose–response curve of a set of drugs for a new biological sample from only a few experiments.

#### Few-Shot Dose–Response Inference

For each held-out study, we conduct a few-shot inference analysis to evaluate how efficiently the foundation model predict dose–responses from a small number of observations. For each sample in the study, we randomly select *n*_few-shots_ (drug, dose, viability) measurements among those available in the dataset, forming the input set to the foundation model. The query set is defined as a different set of (drug, dose) measurements, not contained in the input set. Spearman correlation is computed on the query set between the predicted viability and the ground truth viability. We conduct the analysis for *n*_few-shots_ ∈ {10, 50, 100, 200}, with five random repetitions for each value of *n*_few-shots_. Fig. 6b reports the evolution of Spearman correlation with respect to *n*_few-shots_, with error bars representing two standard deviations across repetitions.

#### Baselines

We compare the foundation model (FM) and its finetuned version (FM++) against three baselines: TabPFN, XGBoost, and Ridge regression. For FM++, a first inference is performed using the few-shot data as both input and query sets. The linear head is then finetuned to match the predictions with the viability values of the few-shot data. TabPFN operates in the same few-shot fashion as the FM: for each sample, it takes as input the *n*_few-shots_ input set and the corresponding query set, and directly outputs the predicted query viabilities. XGBoost and Ridge regression, in contrast, require explicit training. For each sample, these models are fitted on the input set of *n*_few-shots_ observations and then used to predict the query viabilities. Hyperparameters are selected via grid search with cross-validation. For XGBoost the number of estimators is tuned from the set {100, 200, 300} and the learning rate from the set {0.01, 0.1, 0.2}, for Ridge the regularization trade-off parameter alpha is tuned from the set {0.001, 0.01, 0.1, 1.0, 10.0, 100.0}. To provide a drug embedding to these baselines, we use the Morgan fingerprint^169^ over 512 bits. As we observe poor performances of TabPFN with such a large number of features, we extract the 50 first components of the PCA from the matrix of Morgan fingerprint features for every drugs in the PPC dataset.

#### Embedding-Based Prediction of Molecular Features (**Fig. 6c–d**)

We focus on the Beat AML cohort, which provides paired mutation profiles and gene expression measurements for every sample. To obtain sample-level representations, we extract embeddings from the foundation model (FM) by averaging the final token embeddings over the sequence of queries.

#### Prediction of Gene Mutations

Using these fixed embeddings, we train linear probes to predict binary mutation status. To ensure sufficient prevalence, we retain genes with at least 10 mutated samples in Beat AML. We include only samples with available mutation calls and at least 50 profiled drugs (covering over 80% of the cohort), which yields more stable embeddings. For each sample, embeddings are recomputed for *n*_few-shots_ ∈ {10, 50, 100, 200} as in the few-shot setup. Probes are evaluated with 5-fold cross-validation, and we report the mean AUC across folds. Each configuration is repeated 5 times with independent random selections of doses, and metrics are averaged across repetitions. Fig. 6c plots AUC as a function of *n*_few-shots_ and highlights the 4 genes with the highest performance.

#### Prediction of Gene Expression

For continuous gene expression, we aggregate signals at the pathway level using Hallmark gene sets. Let *M* ∈ R^*n*samples×*n*genes^ denote the expression matrix; we normalize each row to sum to 10^6^ and then apply log(*M* + 1). For each pathway, we subset to its member genes and compute the first principal component, yielding a single pathway score per sample. We then fit Ridge regression models (regularization parameter *α* = 1) to predict pathway scores from the 768-dimensional embeddings. Evaluation mirrors the mutation task: 5-fold cross-validation with 5 random dose selection repetitions, reporting the average performance. Fig. 6d shows performance as a function of *n*_few-shots_, highlighting the 12 most accurately predicted pathways.

#### Multi-Omics Analysis (**Fig. 6e**)

We test whether the model produces consistent signals across gene mutation, pathway activity, and targeted drug response. We define three analyses:

1. KRAS with the Hallmark KRAS signaling up pathway and the MEK inhibitor Trametinib.
2. NPM1 with the Hallmark Apoptosis pathway and the Bcl-2 inhibitor Venetoclax.
3. TP53 with the Epithelial–Mesenchymal Transition (EMT) pathway and the MDM2 inhibitor nutlin-3A.

For each sample, we fix *n*_few-shots_ = 100. We select three dose measurements per tested drug from the dataset. This small set is used both to compute the embedding and to perform few-shot dose–response inference. We run the foundation model and extract the 384-dimensional embedding. We then fit linear probes on these embeddings in a leave-one-sample-out scheme. One probe predicts the mutation status of the gene (KRAS, NPM1 or TP53). The other probe predicts the pathway expression score defined earlier from Hallmark gene sets (KRAS signaling up, Apoptosis or EMT). For the held-out sample, we output a mutation logit and a pathway score.

We also infer drug response under the same few-shot setting. For each sample, we use the model to predict viabilities for the targeted drug over the test dose grid. We integrate these predictions to obtain a predicted AUC for that drug. Lower predicted AUC indicates higher sensitivity.

Fig. 6e is organized by columns and rows. Each column corresponds to one analysis and each row to one Omic type. Each subplot provide a boxplot comparison of the wild-type and mutant groups. We assess differences with a one-sided Mann–Whitney U test and report the p-value above each panel.

### Quantifying study-level batch effects

The per-study comparisons in fig. S15 were restricted to most-granular OncoTree annotations such that the disease has at least three samples and is present in at least two studies. All drugs in the vocab described in Methods and used throughout the above were compared.

### Measuring per-study-pair marginal effects

fig. S6 gives the per-dataset marginal absolute RMSE differences given by adding each dataset to the remaining dataset; that is, for each pair of distinct datasets (*d*_ℎ_, *d*_*t*_), we calculated the absolute RMSE difference on the scored dataset *d*_*t*_ with and without the held-out dataset *d*_ℎ_. These are the effect sizes corresponding to the *q*-values in Fig. 2b. All RMSE values were calculated in the whole-drug-holdout five-fold-cross validation setup used in Fig. 2b.

### Quantifying bioactivity across diseases and drugs

fig. S16 gives mean z-scores grouped by (overlapping) drug targets and (non-overlapping) disease annotations. Note CHOL and HCC are grouped together, as are LUSC and LUAD—a number of similar diseases have similar drug responses by model imputation, when viewed in the aggregate.

---

**Figure S1:**
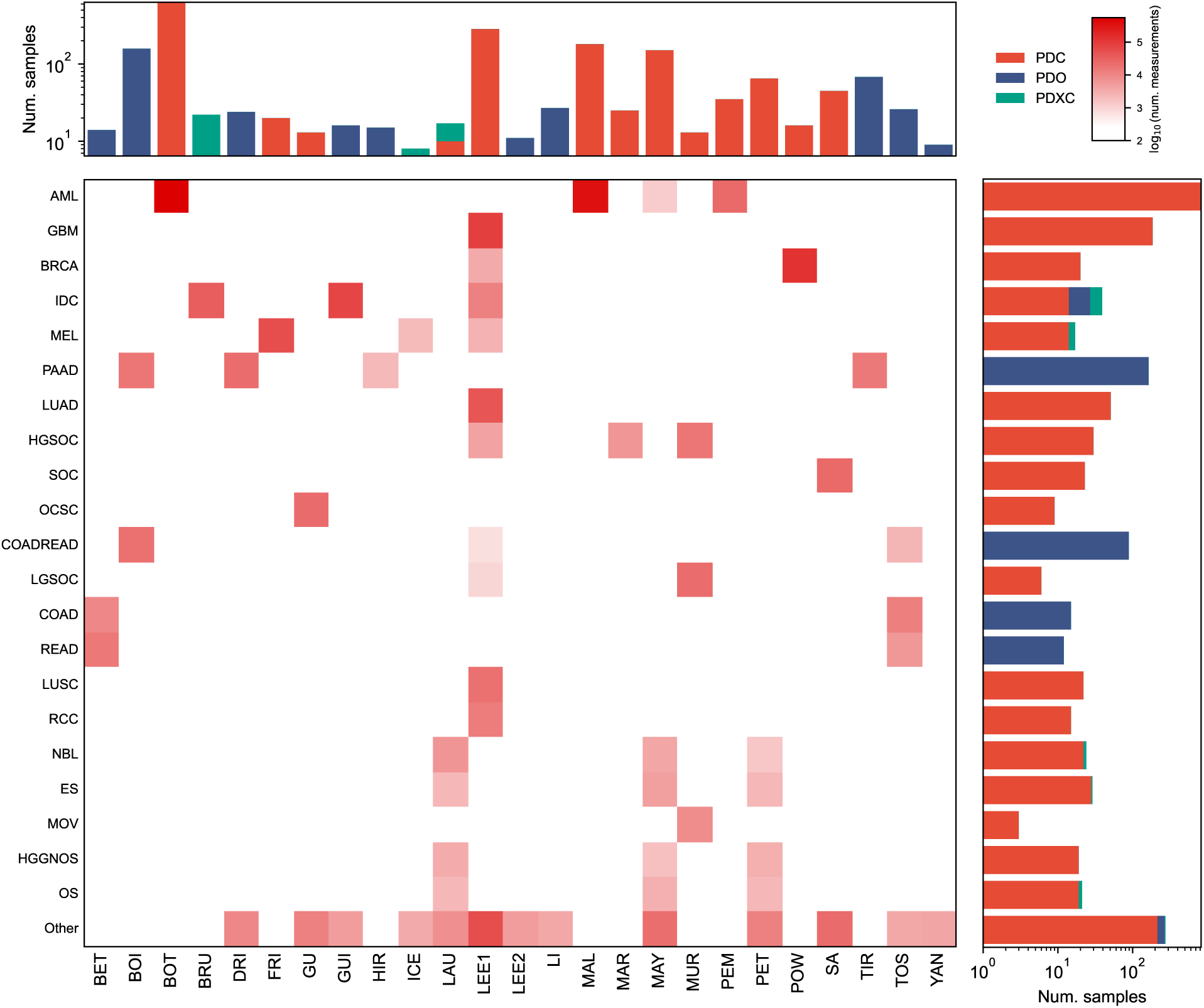
Disease coverage varies across studies. The number of individual viability measurements for the most common disease types, broken down by constituent study. Histograms represent the total number of distinct samples for each study (columns) and disease (rows), colored by model construct (study abbreviations: table S1).

**Figure S2:**
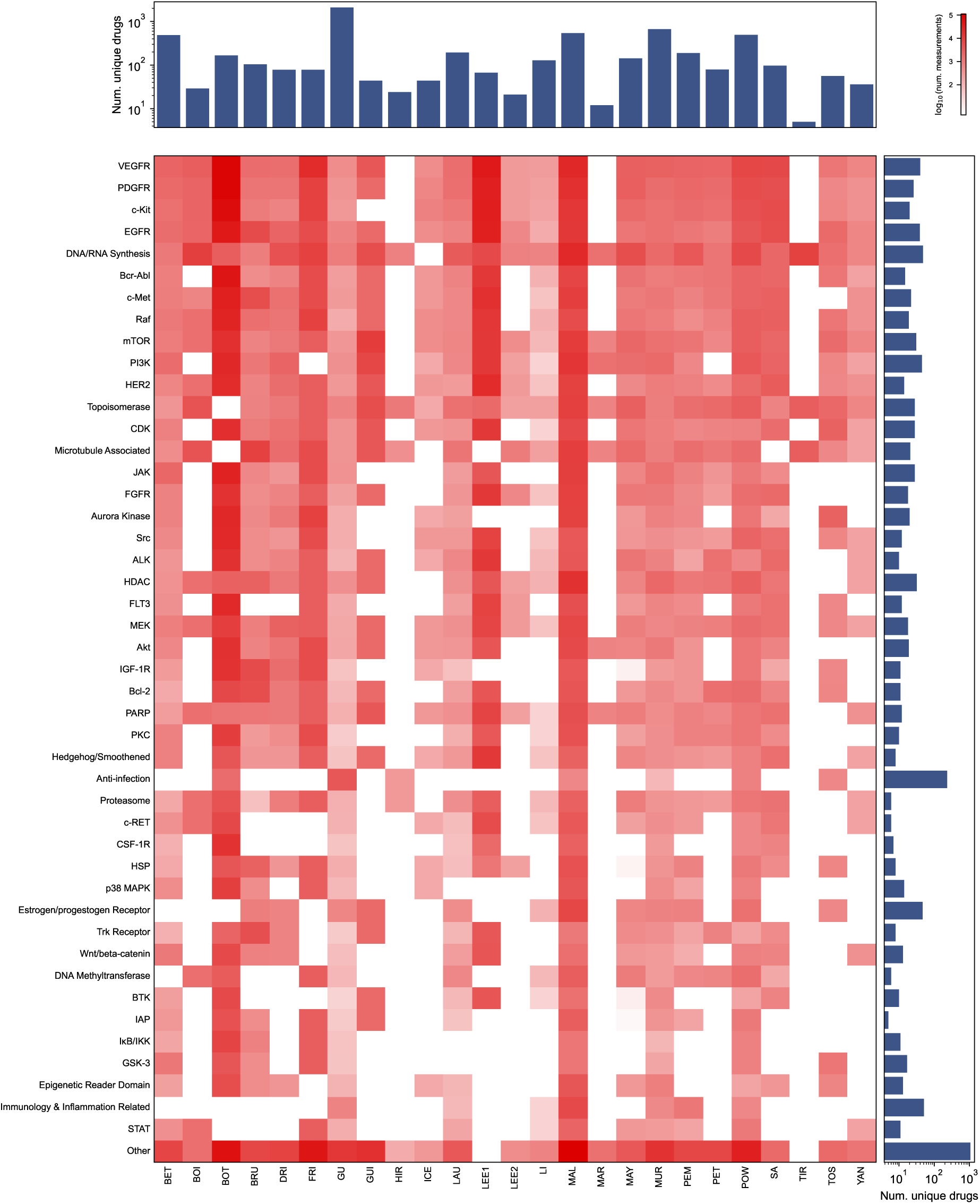
Drug target coverage varies across studies. The number of raw viability measurements stratified by annotated drug target and study. The number of distinct drugs are given as histograms for each study (column) and target (row). As drugs are labeled in general with multiple targets, a given viability measurement may appear in multiple rows (study abbreviations: table S1).

**Figure S3:**
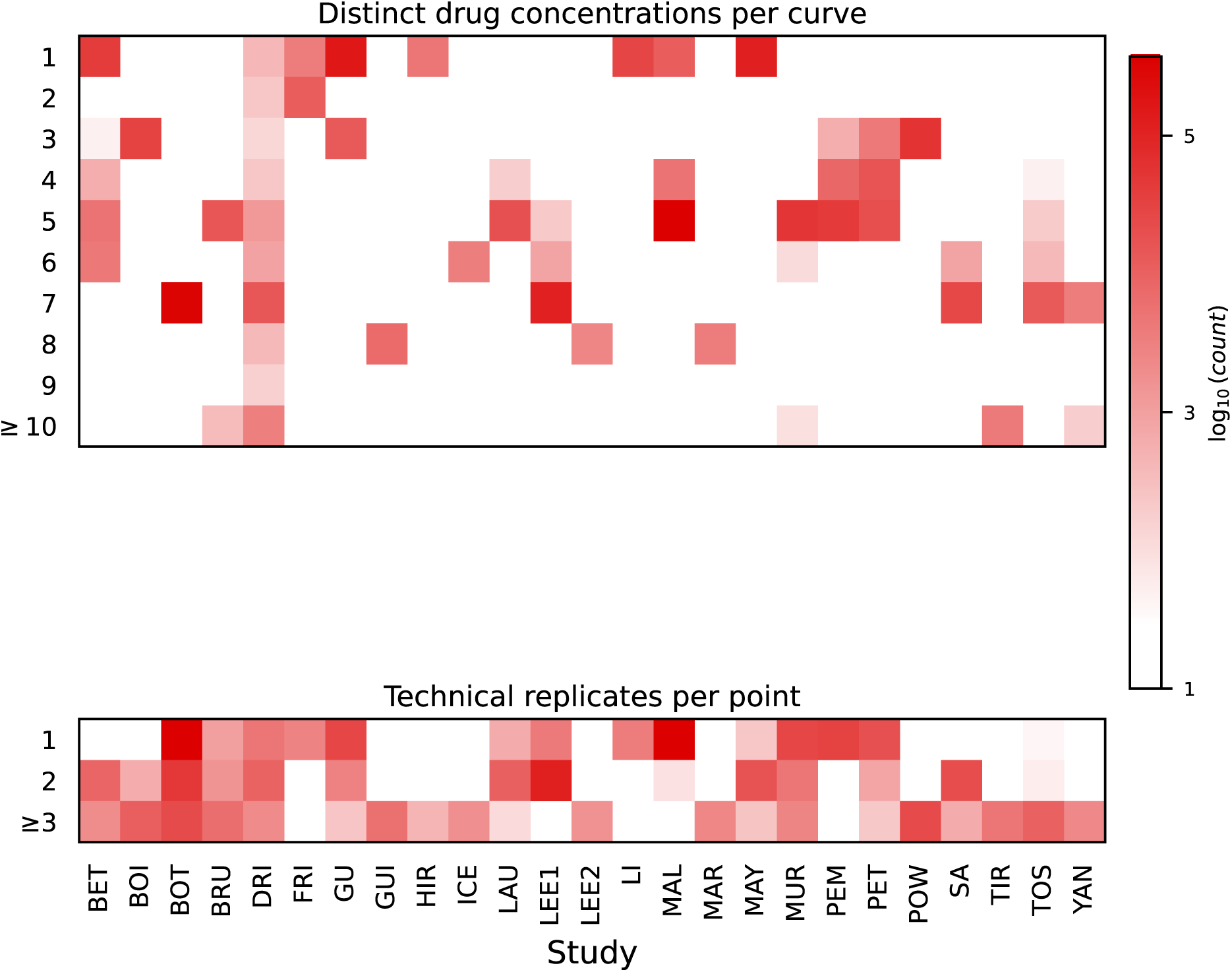
Assay design varies across studies. The numbers of distinct dose concentrations per curve (top) and technical replicates per point (bottom) across different datasets, restricted to single-drug dose-response assays (study abbreviations: table S1).

**Figure S4:**
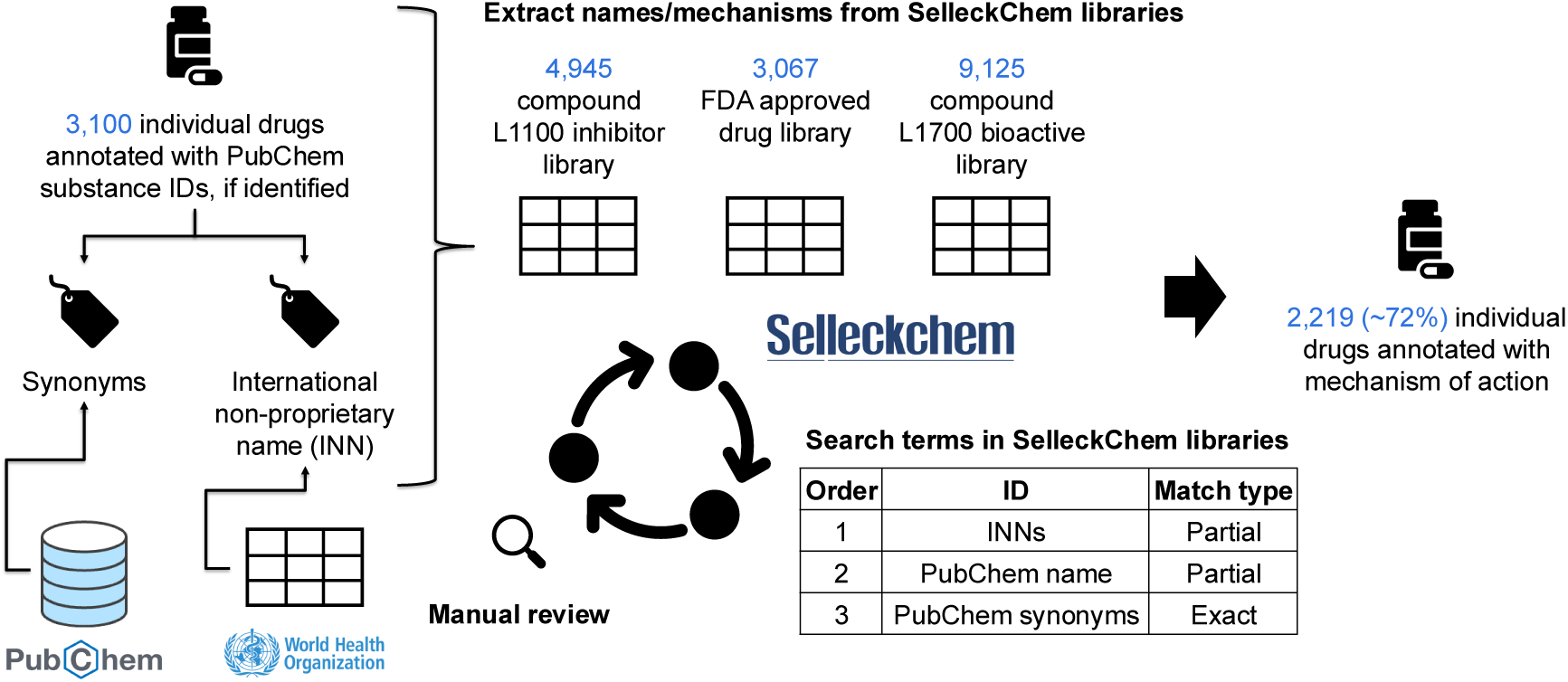
Drug target annotations are harmonized from heterogeneous sources in a semi-automated process.

**Figure S5:**
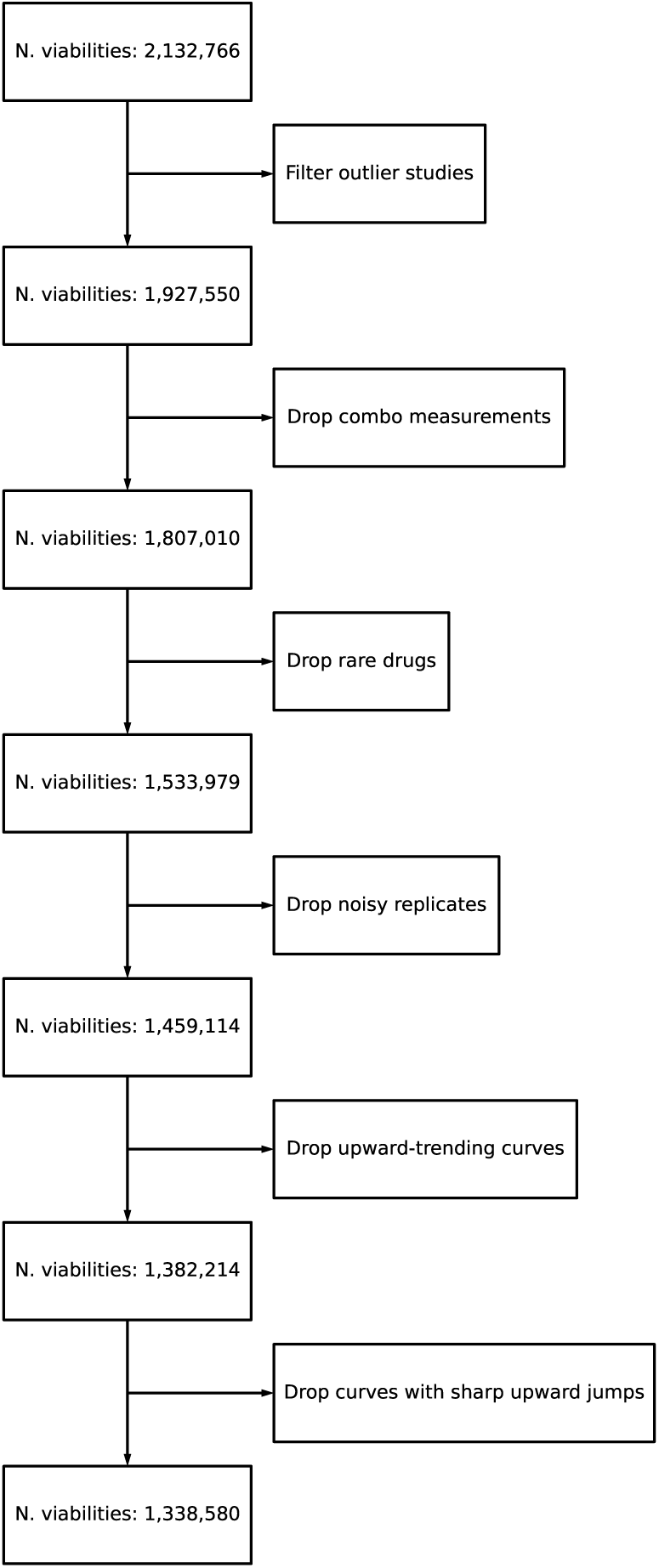
The ex vivo dataset is filtered for quality before model training. The ex vivo dataset size during individual data filtering steps is given.

**Figure S6:**
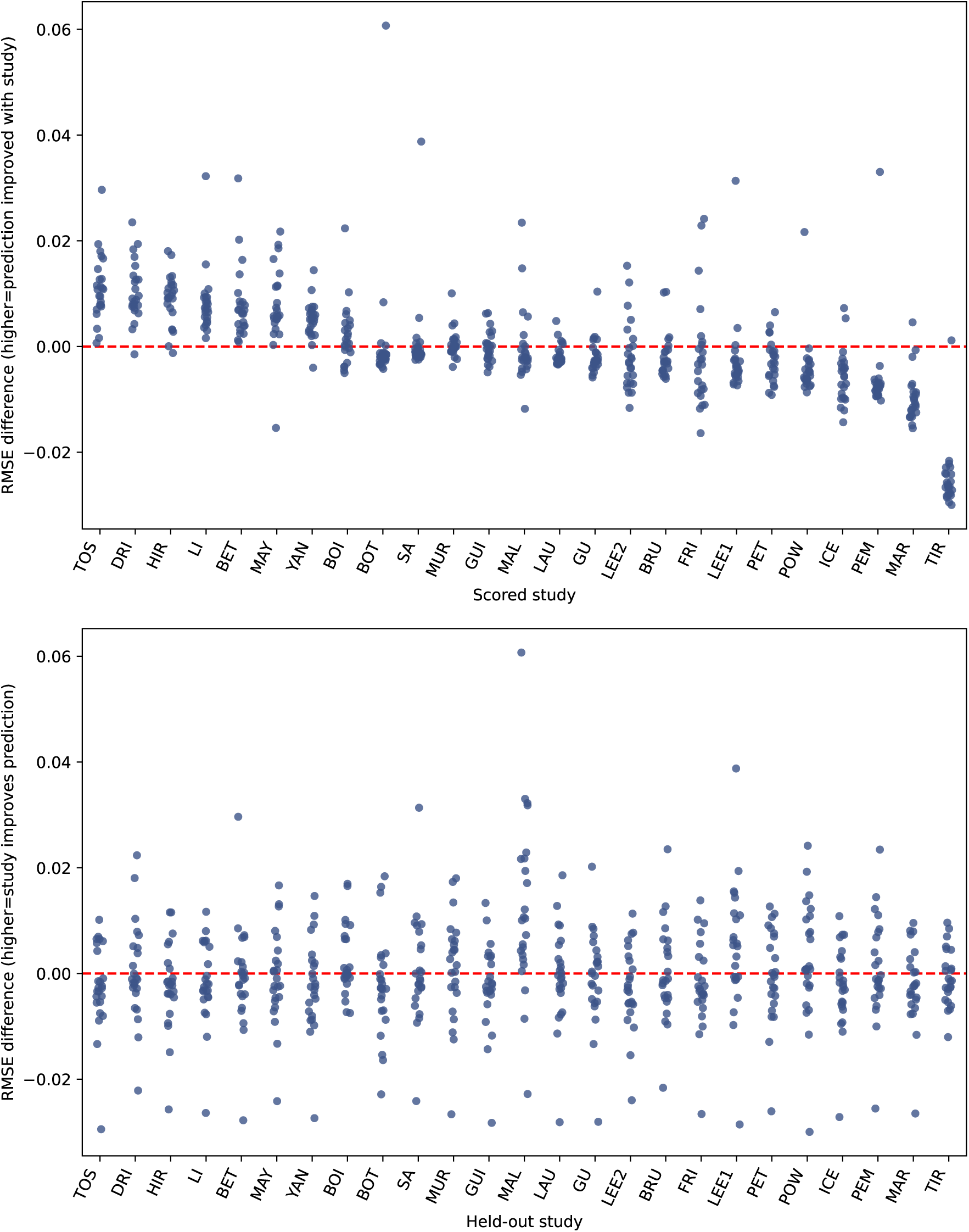
Studies exhibit heterogeneity in pairwise empirical modeling utility. c(*e*_ℎ_ − *e* _*f*_, with *e*_ℎ_ the RMSE ablating the relevant held-out study and *e* _*f*_ the RMSE with the full dataset), grouped by scored study (top) and held-out study (bottom).

**Figure S7:**
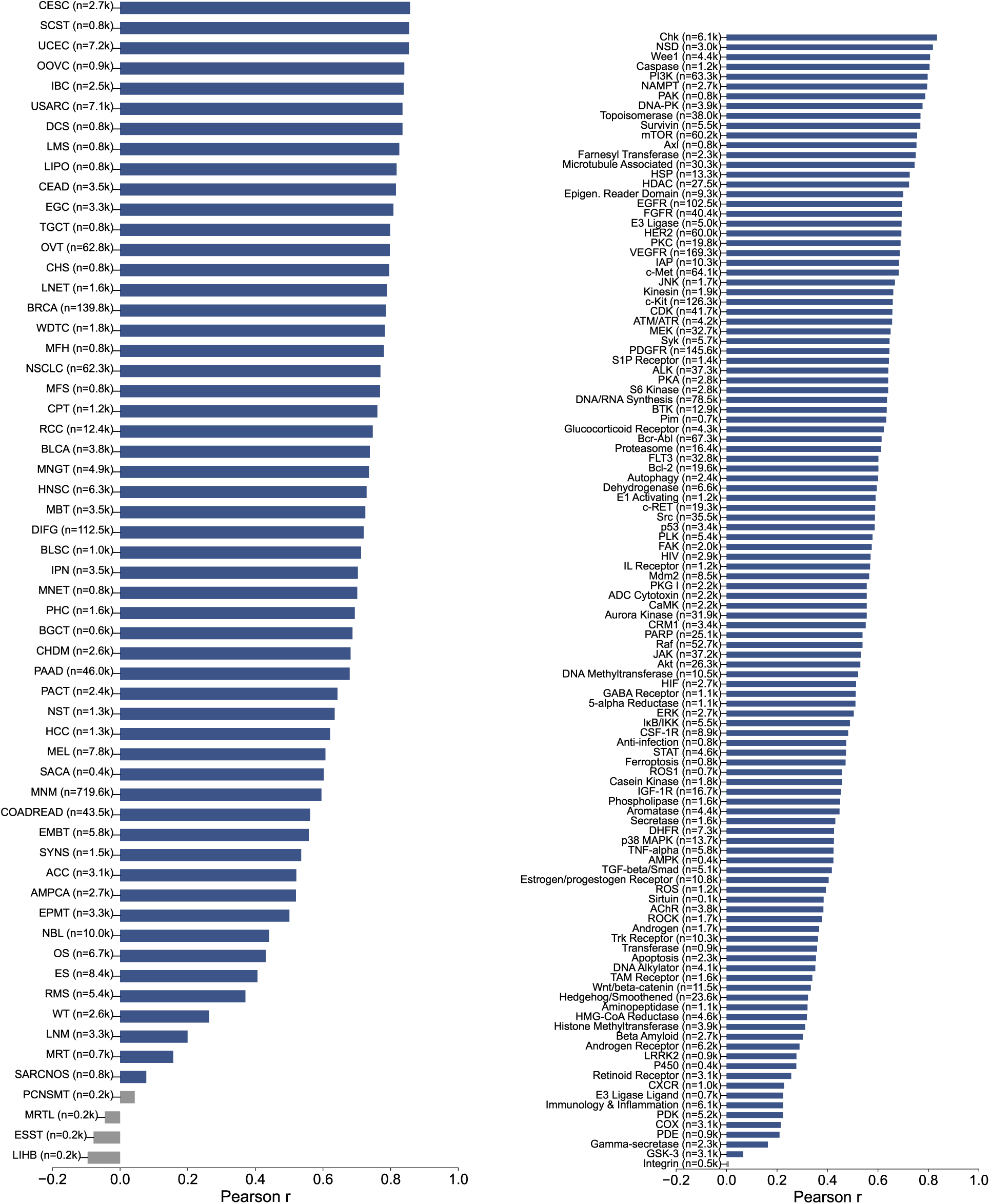
Empirical model performance varies across disease indications and drug targets. Pearson correlation stratified by disease subtype (left) and drug target (right), as depicted in Fig. 2cd, with individual entries labeled.

**Figure S8:**
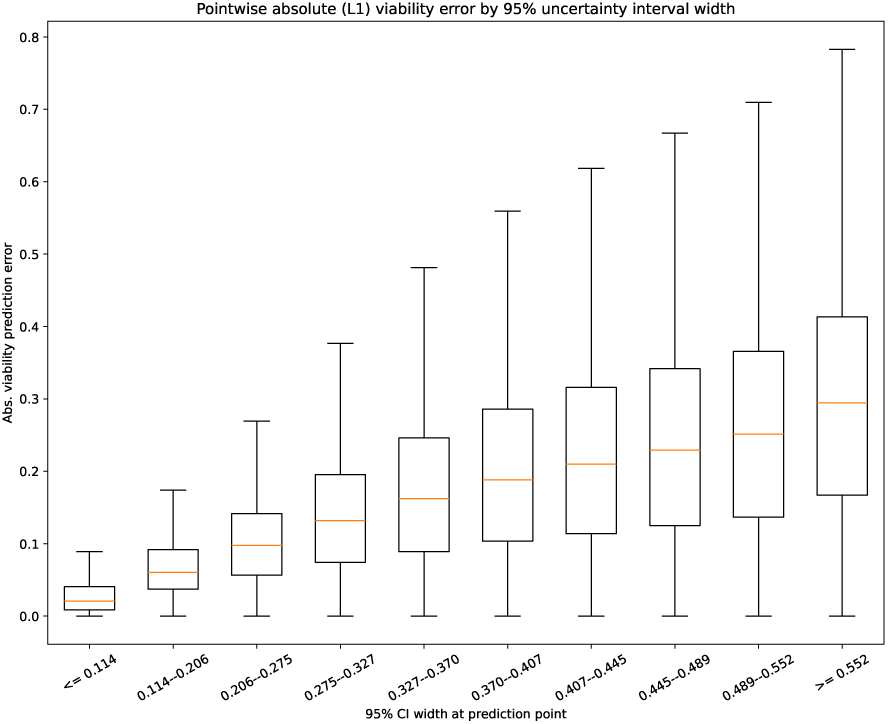
Model prediction error is monotone in model uncertainty. Pointwise mean absolute error for all measurements in the dataset, stratified by the model’s 95% uncertainty interval width at that point. Orange line: median; boxes: first to third quartile; whiskers: 1.5 times inter-quartile range.

**Figure S9:**
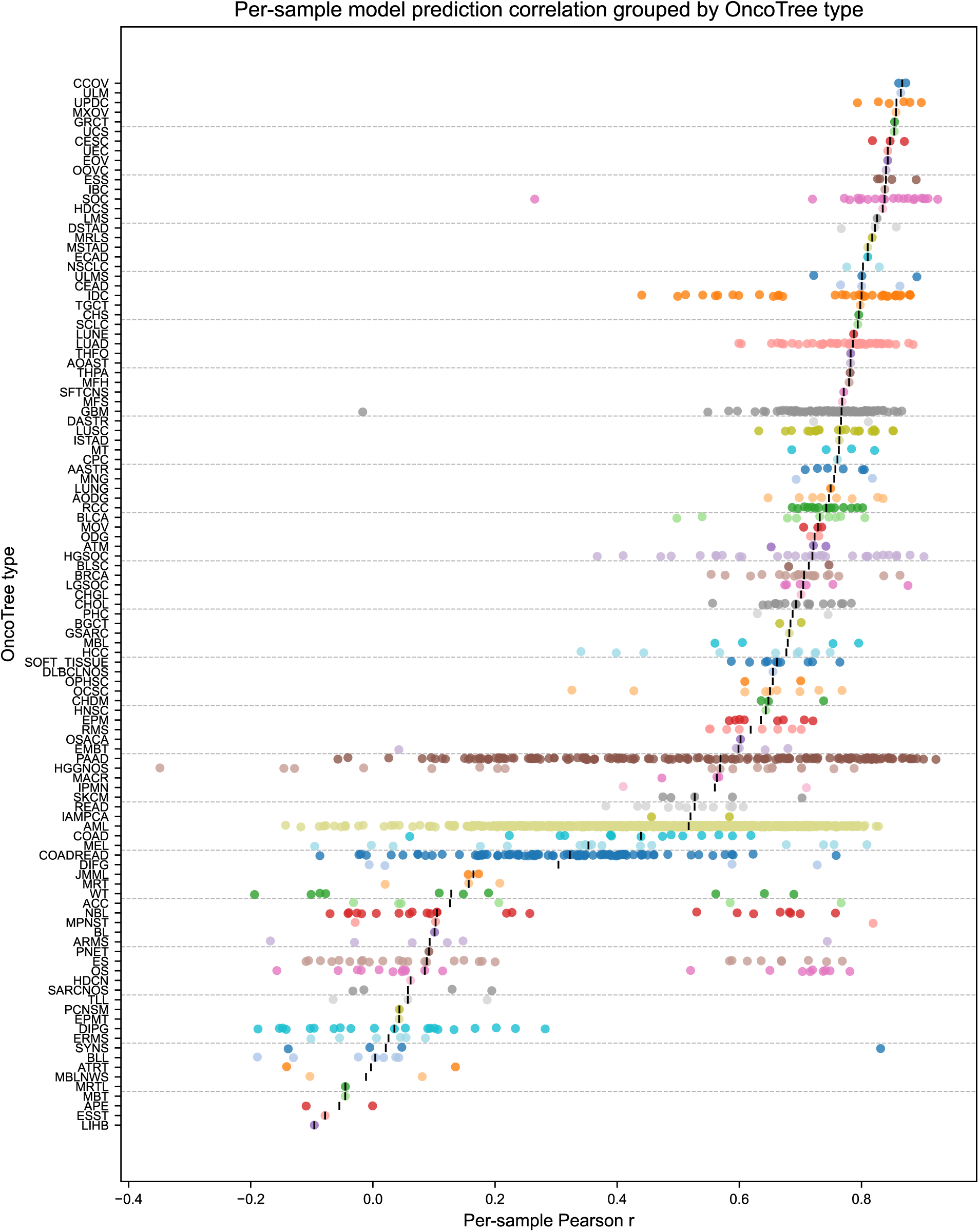
Disease indications exhibit intra- and inter-group heterogeneity in model performance. Per-sample Pearson correlation by disease type, with group medians (black bars).

**Figure S10:**
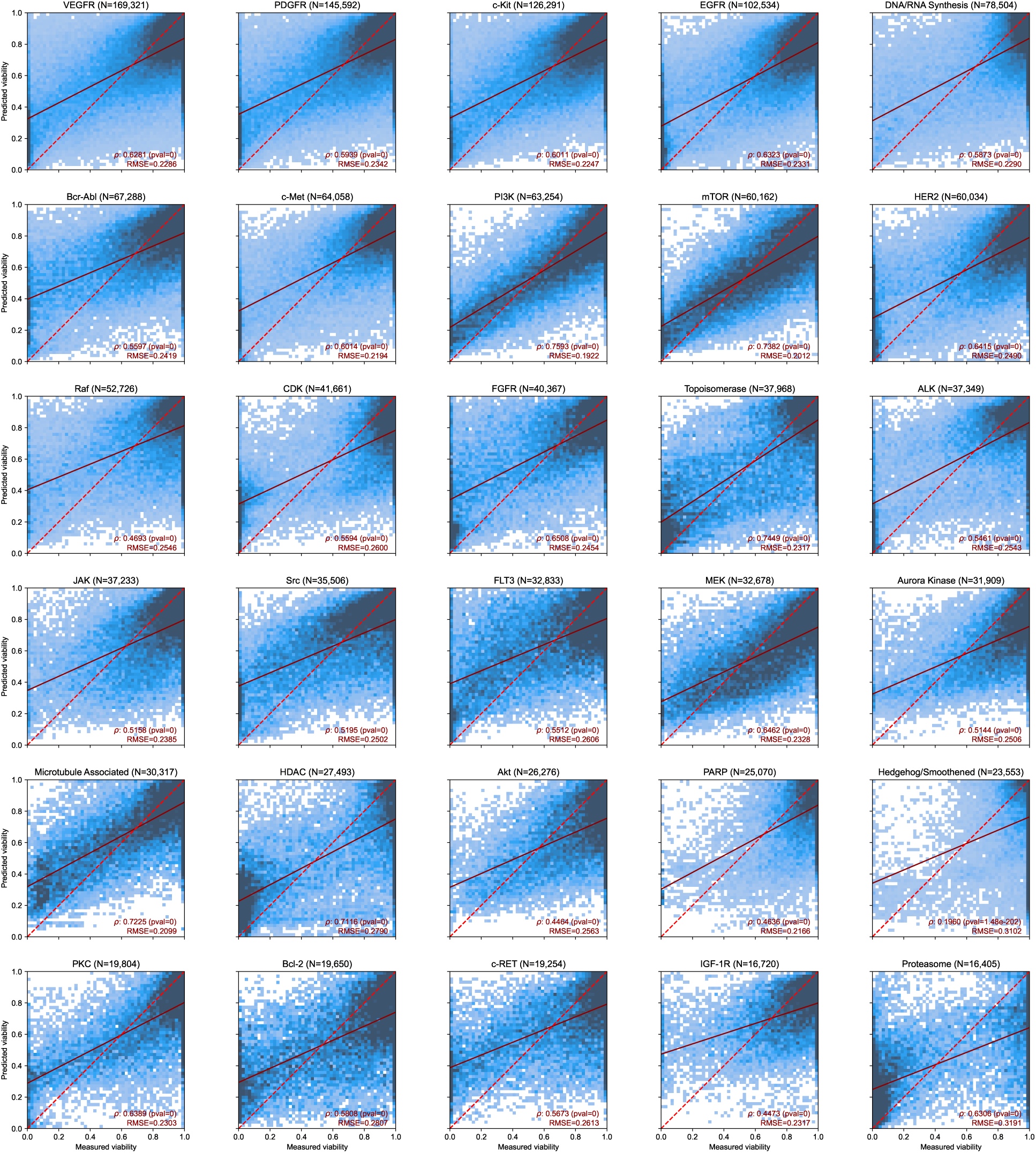
Drug targets exhibit heterogeneity across assay design, measured response, and prediction accuracy. (1 of 4) Predicted vs. measured per-point viability densities, stratified by drug class, with Spearman rank-correlation coefficient, root-mean-squared prediction error, and OLS-regression fit (alongside the *y* = *x* identity line) for each class. Plots are ordered by measurement count in the dataset. All *p*-values in fig. S10– fig. S13 are given by two-sided *t* tests on the Spearman *ρ* coefficients.

**Figure S11:**
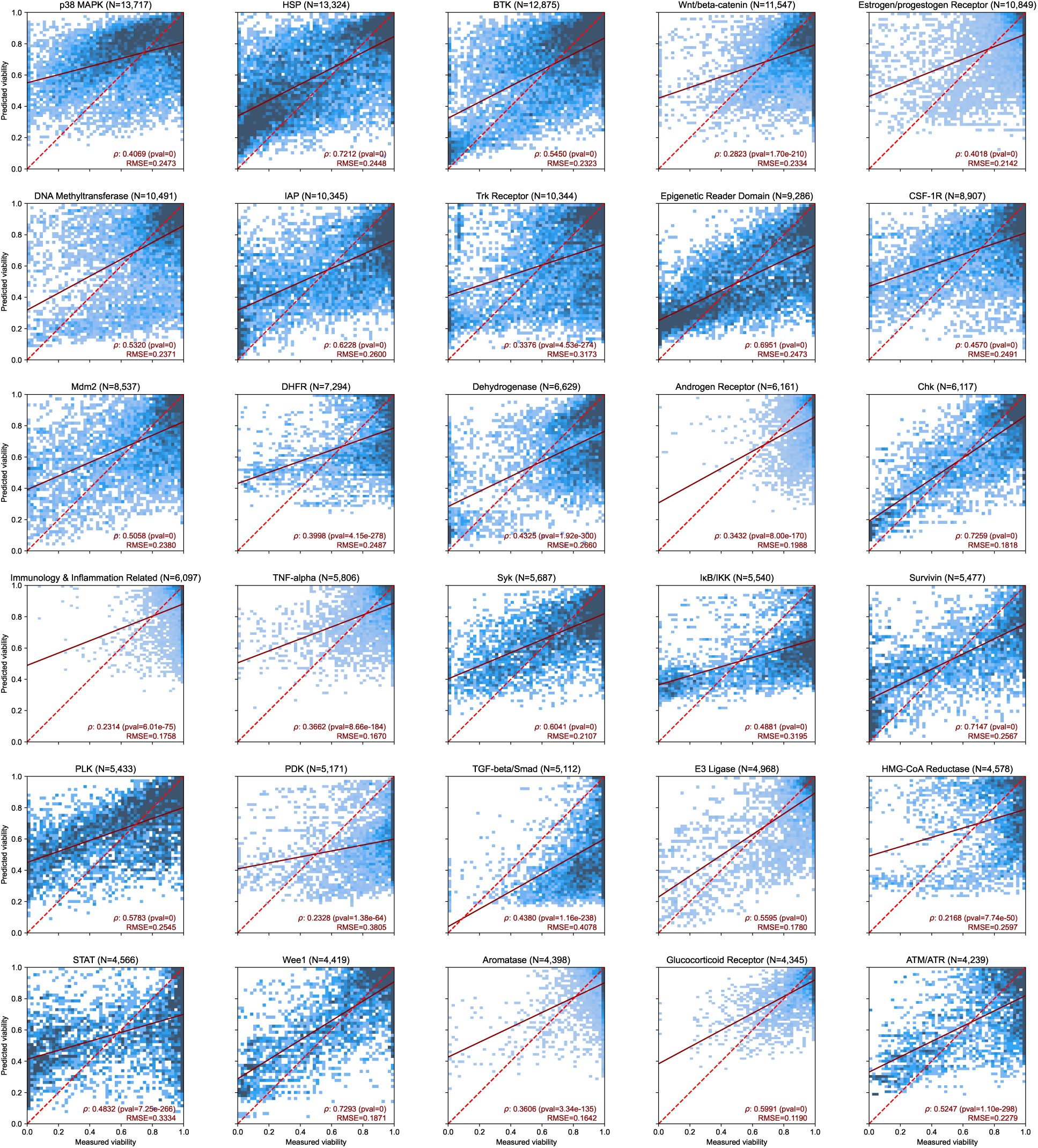
(2 of 4) Continuation of fig. S10.

**Figure S12:**
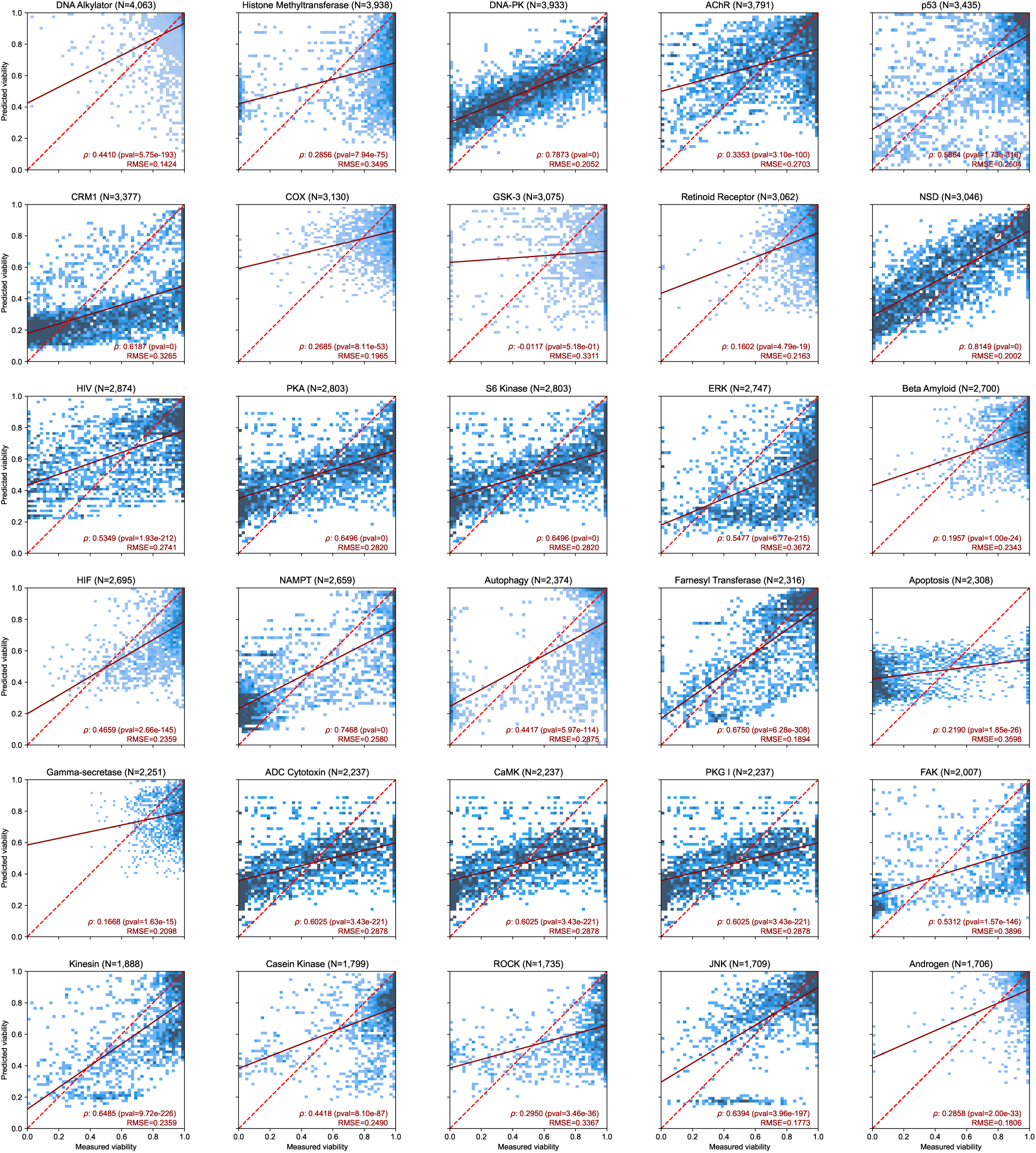
(3 of 4) Continuation of fig. S10.

**Figure S13:**
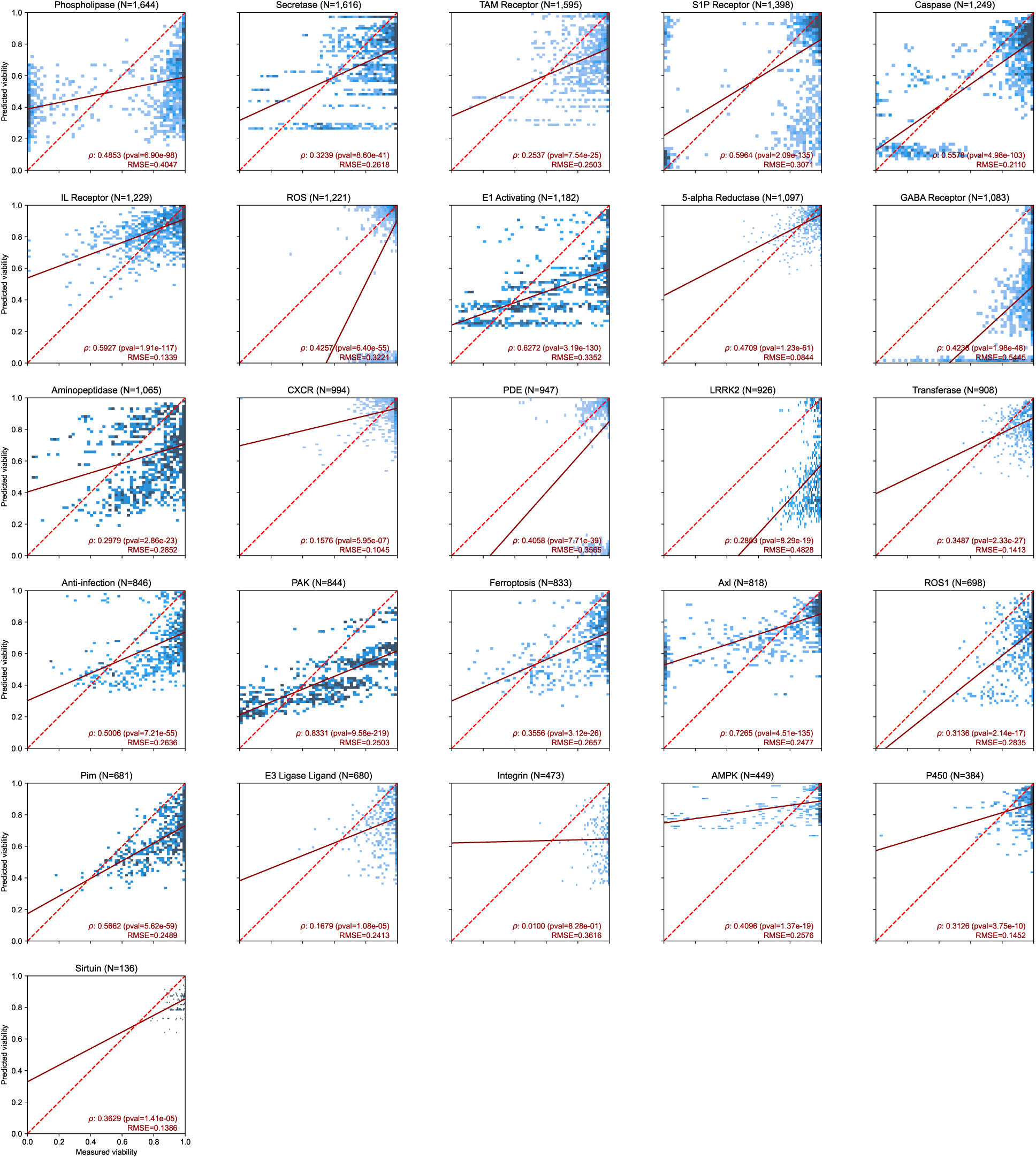
(4 of 4) Continuation of fig. S10.

**Figure S14:**
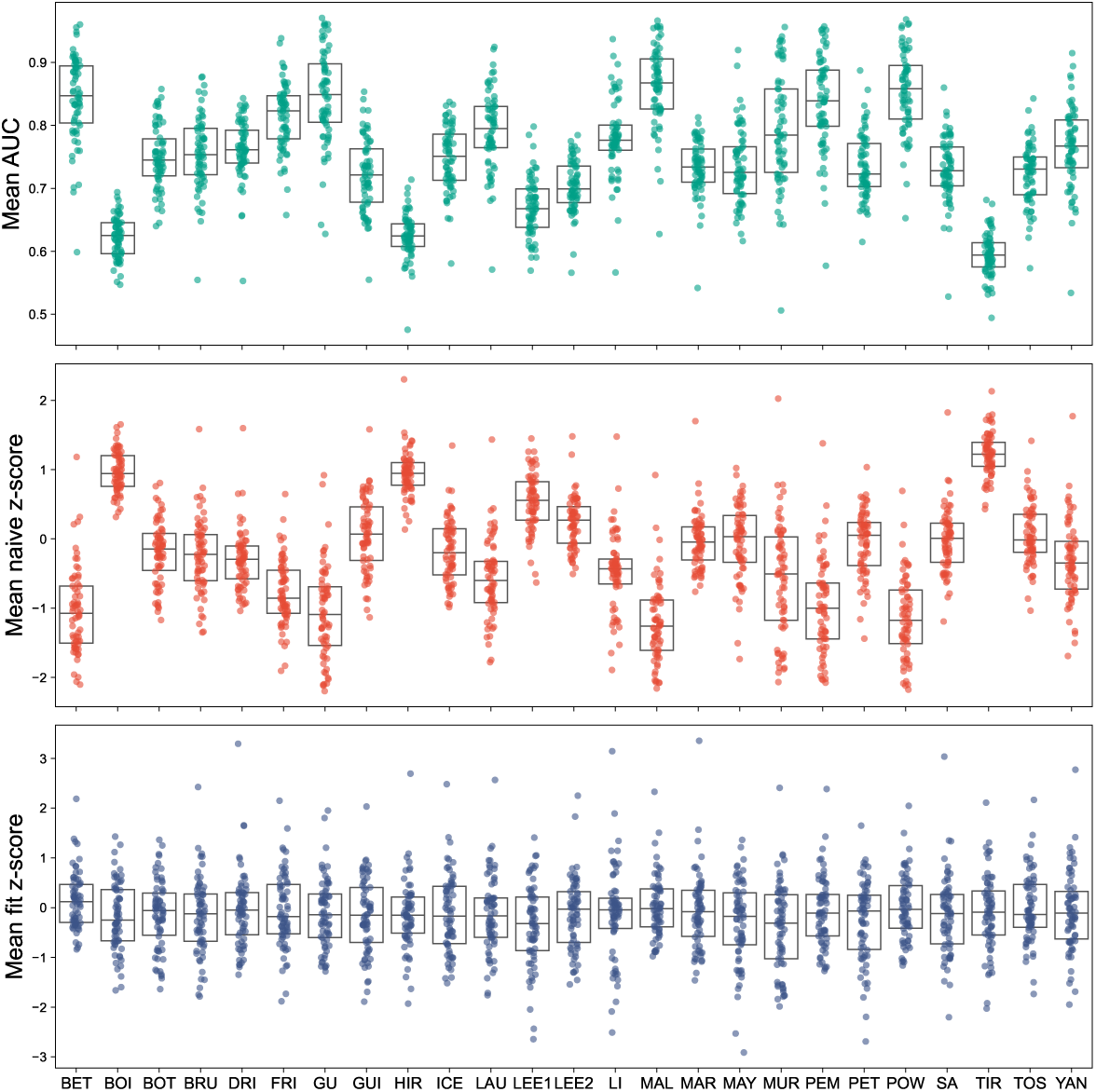
The Empirical Bayes method yields considerably diminished study-level batch effects on z-scores. Per-target AUCs (top), naive z-scores (middle) and empirically fit z-scores (bottom) across studies (study abbreviations: table S1).

**Figure S15:**
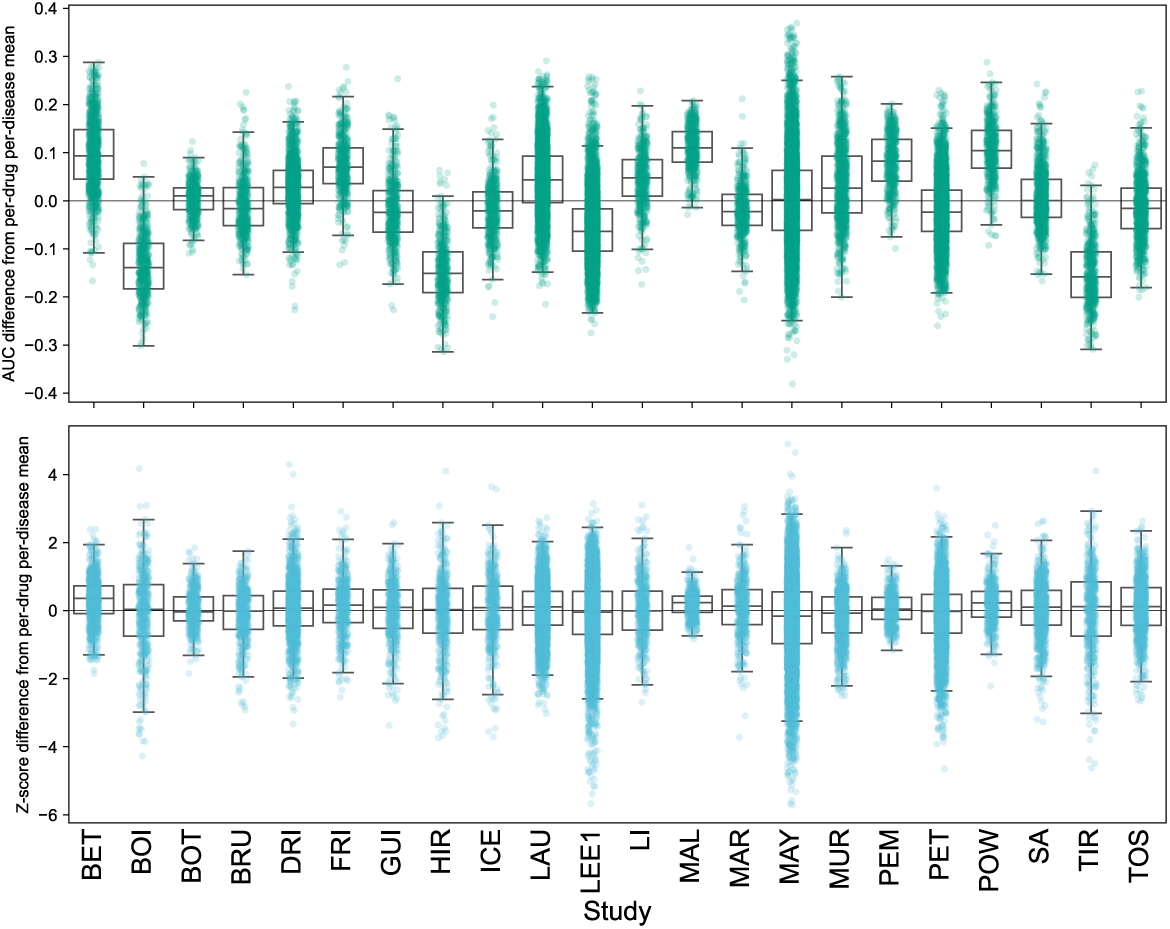
Raw AUC scores exhibit batch effects, which are attenuated by Empirical Bayesian z-scores. Per-drug per-disease differences from study to population means for raw AUC (top) and empirically fit z-scores (bottom). Each point represents a (drug, OncoTree) pair, and quantities are differences in the study’s average score (AUC or z-score, respectively) on that (drug, disease) pair compared to the global average for that pair. The bottom plot shows considerably attenuated study-level bias compared to the top plot. (boxes: quartiles and medians; whiskers: 1.5 IQR; study abbreviations: table S1).

**Figure S16:**
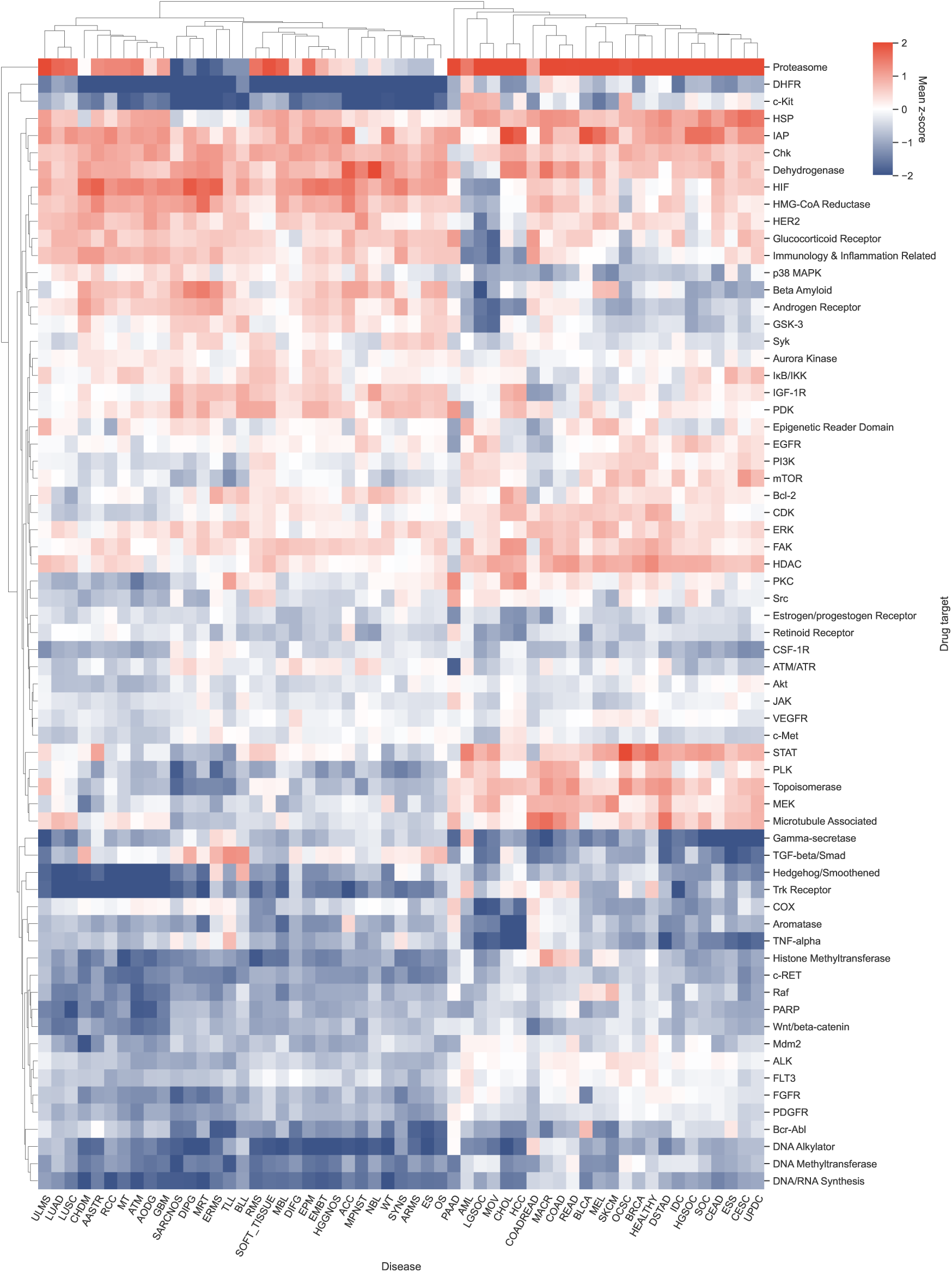
Imputed disease-target z-scores exhibit systematic response heterogeneity. Mean AUC-derived z-scores of model outputs for each target with at least 3 drugs and each disease with at least 3 samples. Rows and columns are clustered by average distance in Euclidean space.

**Figure S17:**
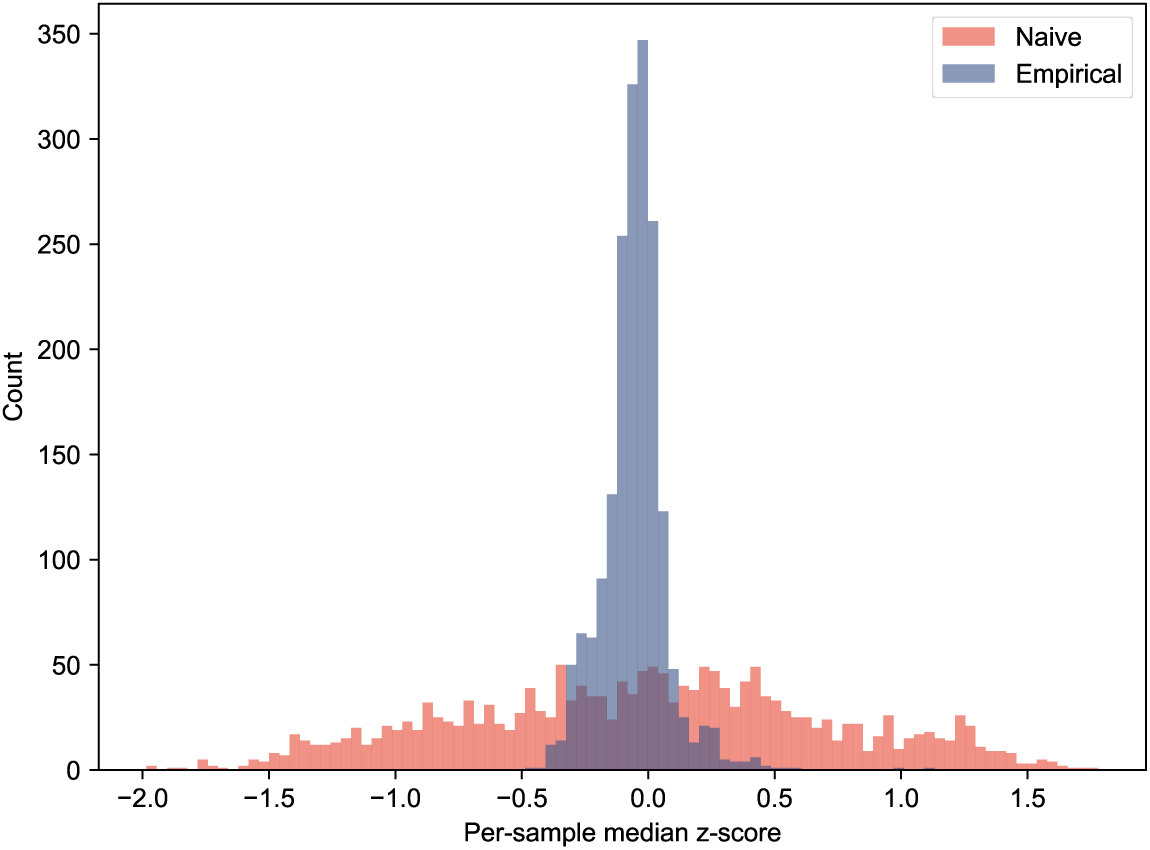
Empirical Bayes z-scores have considerably fewer skewed median per-sample values. Distribution of median z-scores per sample for naive z-score settings (red) and empirically fit z-scores (blue).

**Figure S18:**
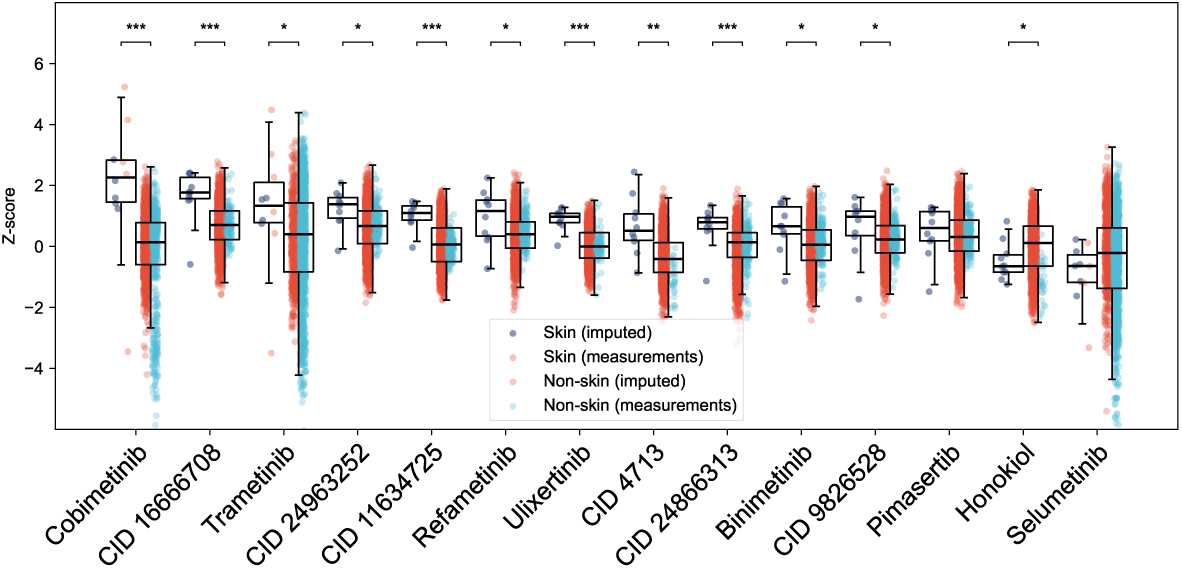
Fully-imputed z-scores mix with measurement-derived z-scores for MEK/ERK-targeting drugs across samples. Distribution of z-scores for all drugs labeled as targeting MEK/ERK, with z-scores from pure imputations (i.e. dose-response curves fully imputed) separated from z-scores derived from curves with measurements in the dataset. (BH-corrected p-values from two-sided Mann-Whitney U tests, *: *p* < 0.1; **: *p* < 0.01; ***: *p* < 0.001. Boxes: first and third quartiles; line: median; whiskers: 1.5 × IQR).

**Figure S19:**
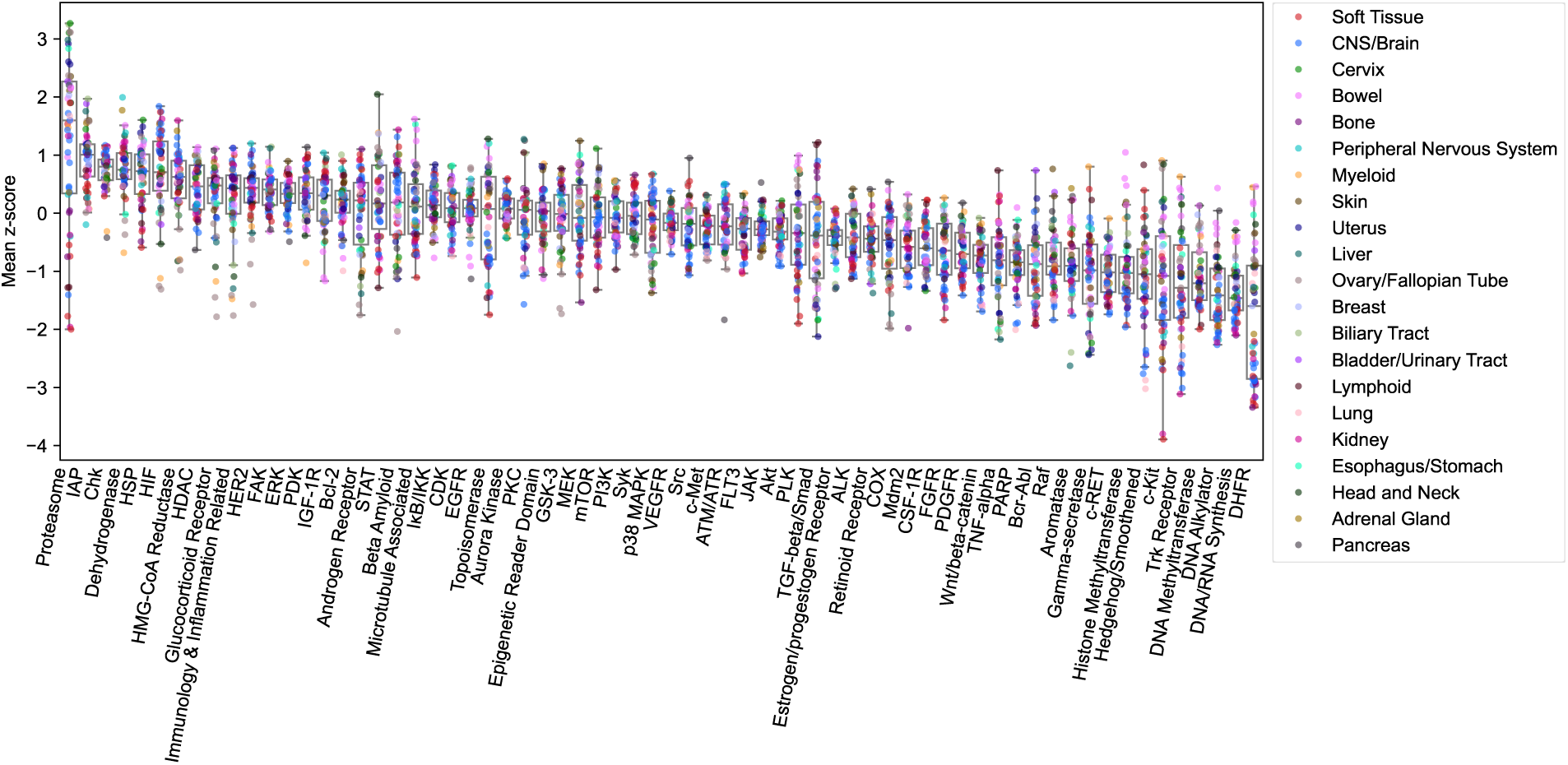
Per-tissue responses exhibit intra- and inter-group heterogeneity across drug targets. Mean z-scores per-target and per-primary-disease-tissue-type, grouped by target (boxes: first and third quartiles; line: median; whiskers: 1.5 × IQR).

**Figure S20:**
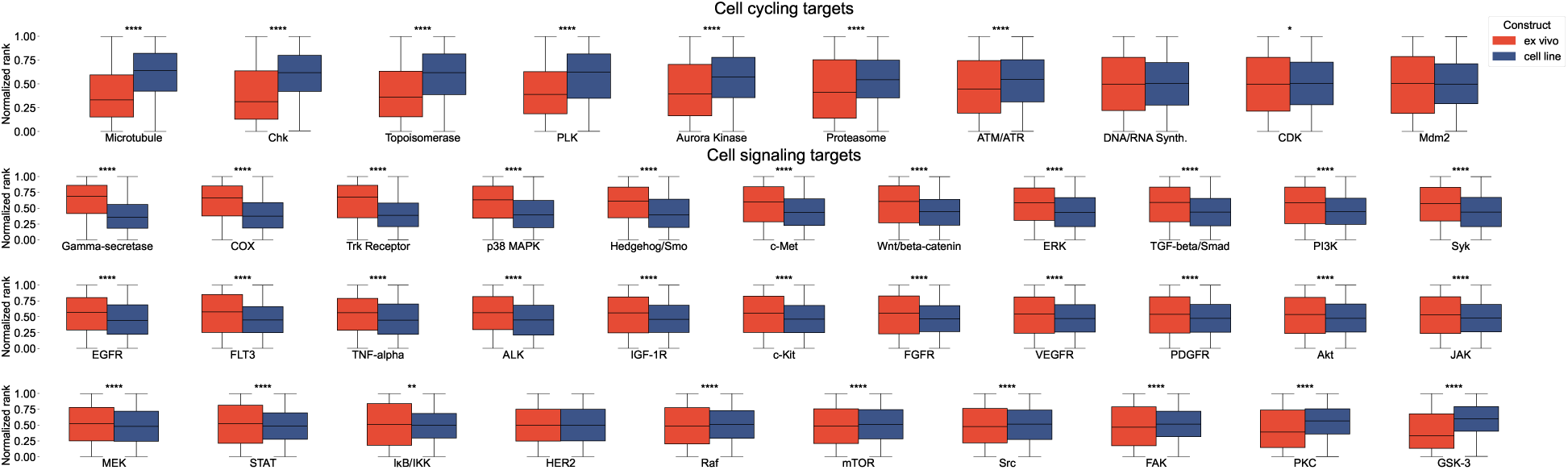
Cell-cycling targeting drugs are more comparatively efficacious on cell line constructs; signaling-targeting drugs are more comparatively efficacious on ex vivo constructs. Distributions of normalized ranks for cell cycling (top) and signaling (bottom) targets, via two curated subsets of the targets in the dataset (boxes: IQR; horizontal lines: medians; whiskers: 1.5 times IQR). Ranks were computed per drug, normalized to between 0 and 1 (higher is more efficacious). Asterisks indicate BH-adjusted significance from two-sided Welch’s t-tests on ex vivo versus cell-line ranks. (*: *q* < 0.1; **: *q* < 0.01; ***: *q* < 0.001; ****: *q* < 0.0001).

**Figure S21:**
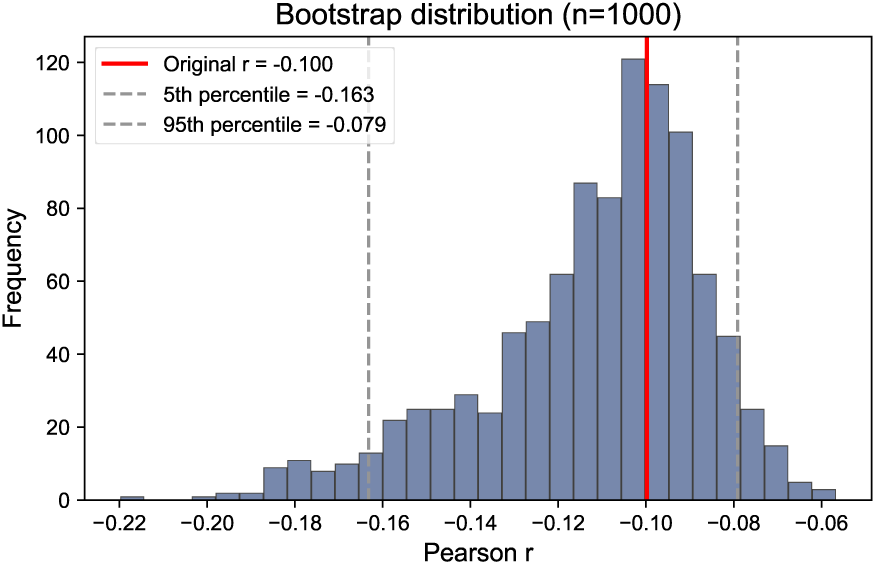
The correlation between RMSE difference and absolute z-score difference when adding cell line data is robust to data resampling. The Pearson correlation *r* value for the linear fit from per-disease/target absolute z-score difference to RMSE improvement, in Fig. 5, is stable under bootstrapped data resampling.

**Figure S22:**
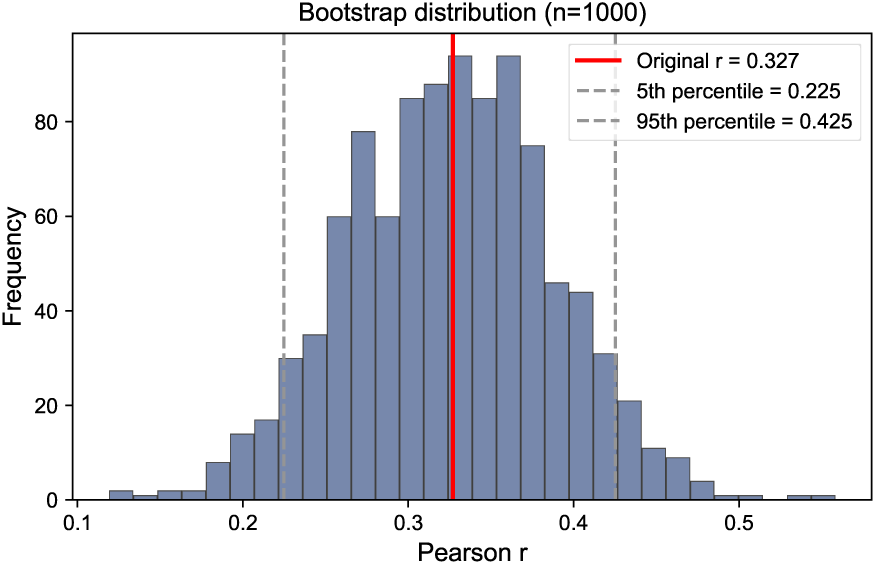
The correlation between TMB and nearest-neighbor TMB on cell line samples is robust to data resampling. The Pearson correlation *r* value for the linear fit from TMB to nearest-neighbor TMB (in log/log space), as given in Fig. 5, improvement, in Fig. 5, is stable under bootstrapped data resampling.

**Figure S23:**
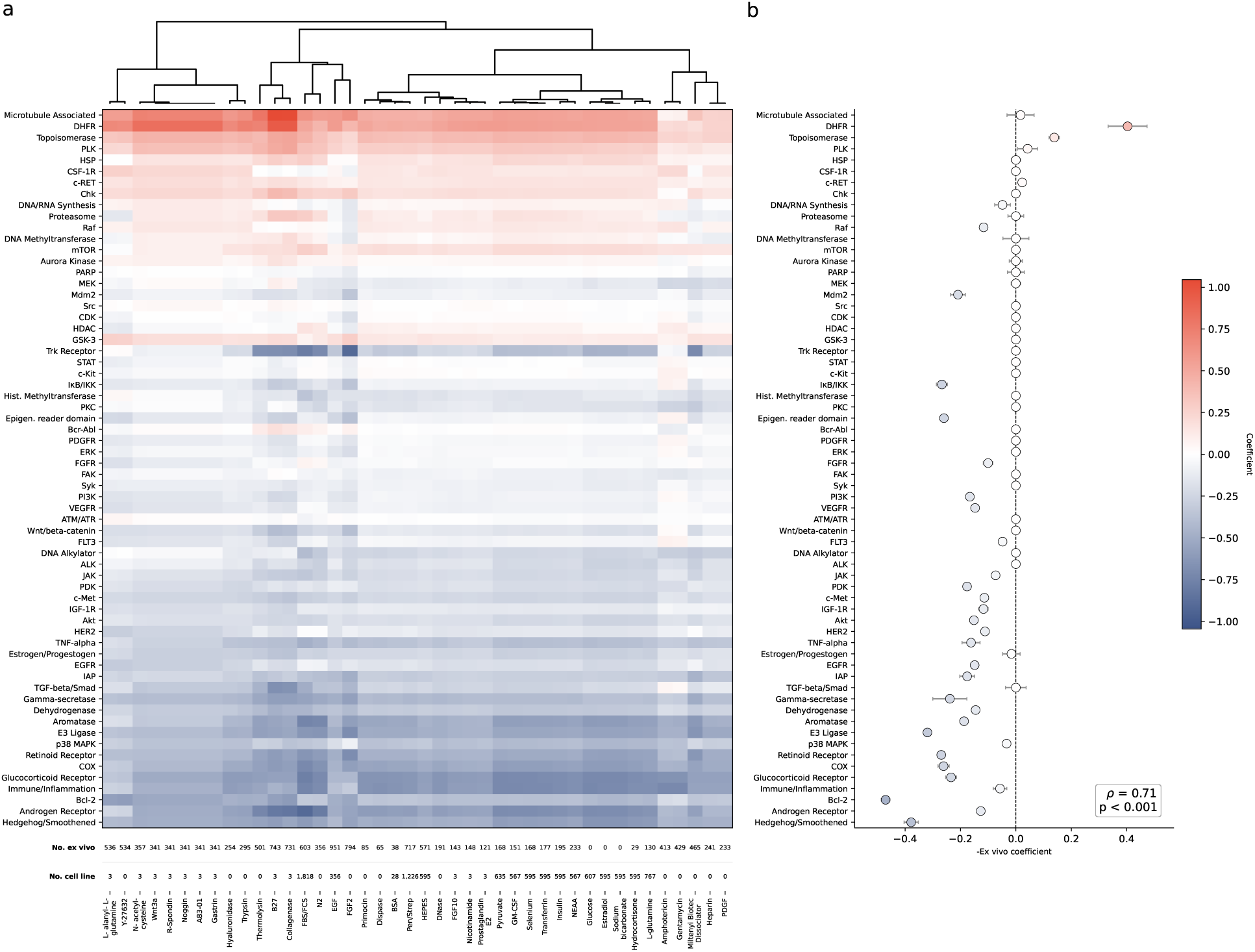
Patterns in ex vivo sensitivity remain consistent when controlling for media additives. (a) Average per sample z-scores by annotated drug target were regressed on the presence or absence of a media additive in a study and whether the sample was a cell line or an ex vivo construct. The negative coefficients on the one-hot encoded ex vivo covariate are shown here per drug target (vertical axis) and per additive (horizontal axis). Rows are ordered according to Fig. 5(e) and columns are grouped using hierarchical agglomerative clustering over Euclidean distance as shown in the dendrogram above. (b) The negative coefficients on the one-hot encoded ex vivo covariate are shown for ℓ_1_-regularized multiple regressions to control for the effects of additives, construct type, and study ID simultaneously (point: coefficient; whiskers: bootstrapped standard errors of the mean). Spearman’s *ρ* of the order of negative ex vivo coefficients to the row order of drug targets in Fig. 5(e) is shown. Full names for additive abbreviations and acronyms are available in table S6.

**Figure S24:**
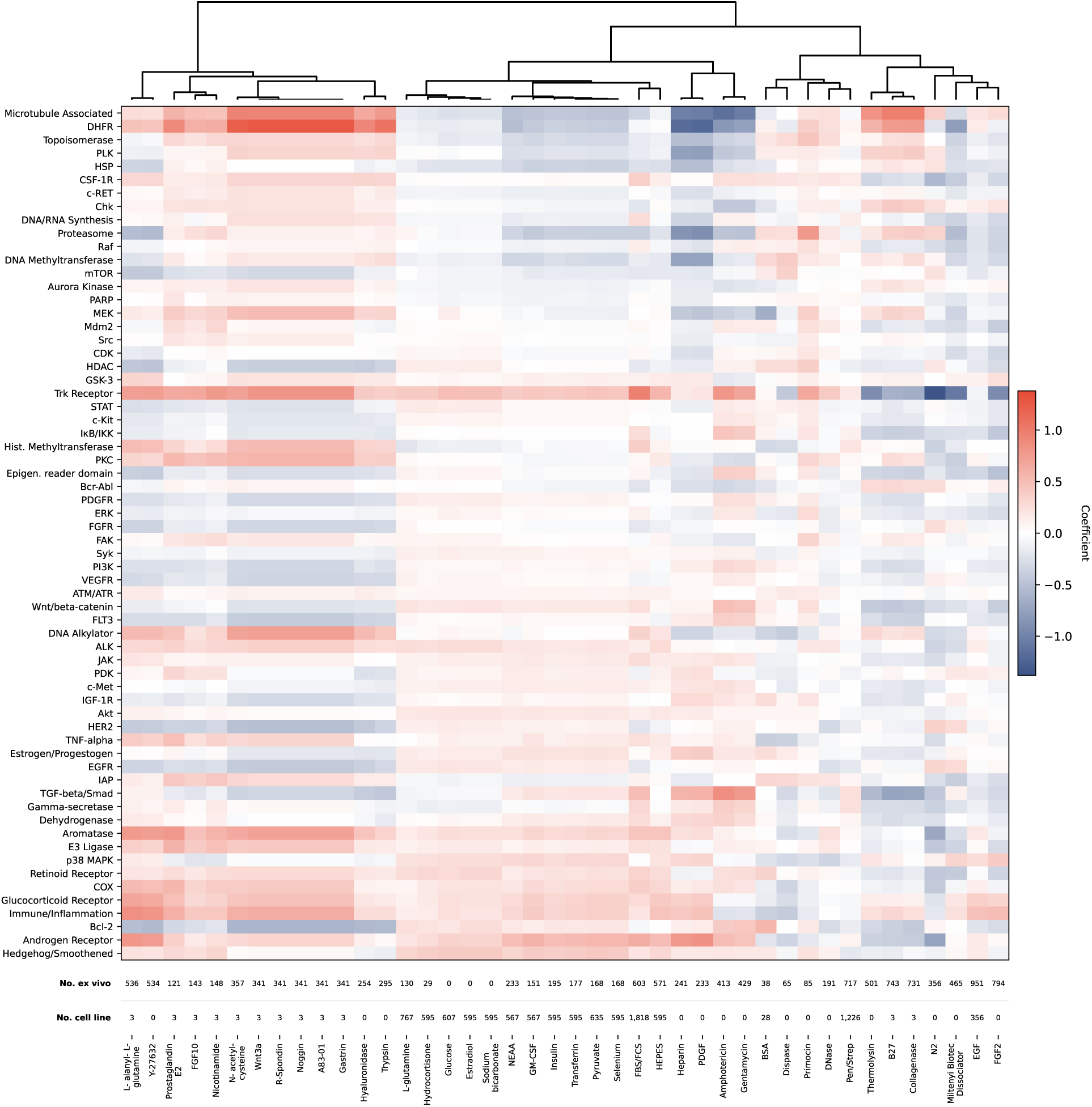
Additives partially explain drug target variability. Average per sample z-scores by annotated drug target were regressed on the presence or absence of a media additive in a study and whether the sample was a cell line or an ex vivo construct. The negative coefficients on the one-hot encoded media additive covariates are shown here per drug target (vertical axis) and per additive (horizontal axis). Rows are ordered according to Fig. 5(e) and columns are grouped using hierarchical agglomerative clustering over Euclidean distance as shown in the dendrogram above. Full names for additive abbreviations and acronyms are available in table S6.

**Figure S25:**
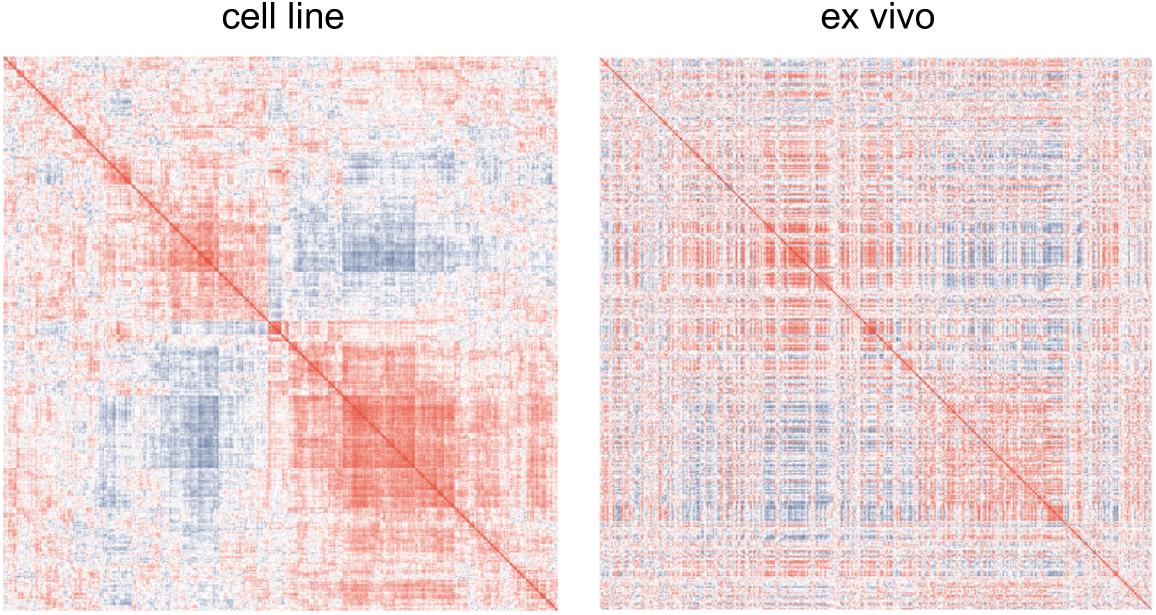
Pairwise drug correlation exhibits similar structure in cell line and ex vivo constructs. Pearson *r* values of mean z-scores calculated across OncoTree groupings exhibit similar blockwise structure, as quantified in Fig. 5.

**Figure S26:**
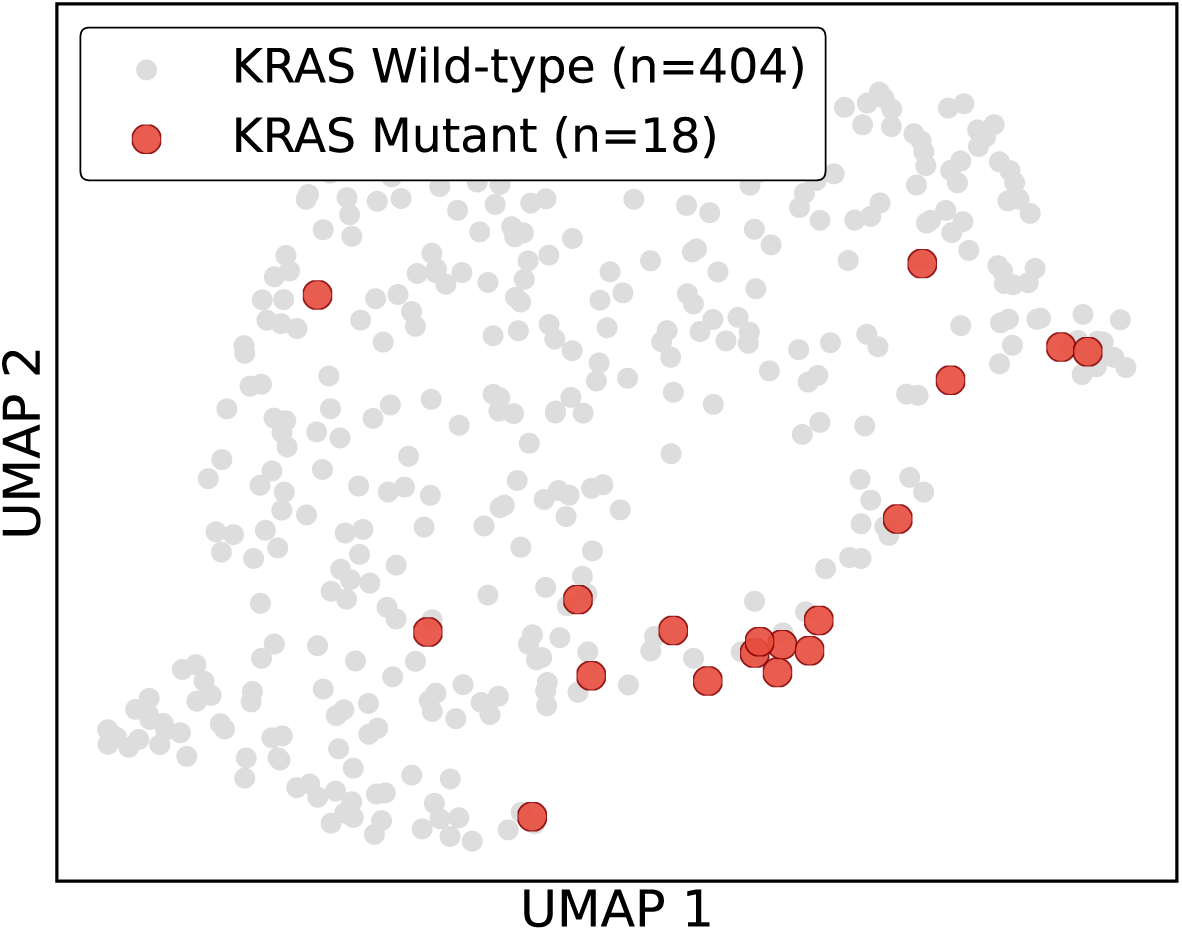
Foundation Model Embedding UMAP Visualization. Two-dimensional UMAP projection of sample embeddings generated by the PPC Foundation Model for the Beat AML cohort. KRAS-mutated samples are highlighted in red.

**Table S1:** Abbreviations used for studies.

| **Abbreviation** | **Study** |
| --- | --- |
| BET | Betge et al.<sup>33</sup> |
| BOI | Boilève et al.<sup>35</sup>, Polit et al.<sup>36</sup> |
| BOT | Bottomly et al.<sup>28</sup> |
| BRU | Bruna et al.<sup>38</sup> |
| DRI | Driehuis et al.<sup>4</sup> |
| FRI | Friedman et al.<sup>24</sup> |
| GU | Gu et al.<sup>25</sup> |
| GUI | Guillen et al.<sup>7</sup> |
| HIR | Hirt et al.<sup>11</sup> |
| ICE | Ice et al.<sup>40</sup> |
| JOH | Johansson et al.<sup>10</sup> |
| LAU | Lau et al.<sup>41</sup> |
| LEE1 | Lee et al.<sup>9</sup> |
| LEE2 | Lee et al.<sup>30</sup> |
| LI | Li et al.<sup>34</sup> |
| MAL | Malani et al.<sup>26</sup> |
| MAR | Martins et al.<sup>15</sup> |
| MAY | Mayoh et al.<sup>29</sup> |
| MUR | Murumägi et al.<sup>16</sup> |
| PEM | Pemovska et al.<sup>27</sup> |
| PET | Peterziel et al.<sup>20</sup> |
| POW | Powell et al.<sup>37</sup> |
| SA | Sa et al.<sup>14</sup> |
| TIR | Tiriac et al.<sup>1</sup> |
| TOS | Toshimitsu et al.<sup>32</sup> |
| YAN | Yan et al.<sup>31</sup> |

**Table S2:**
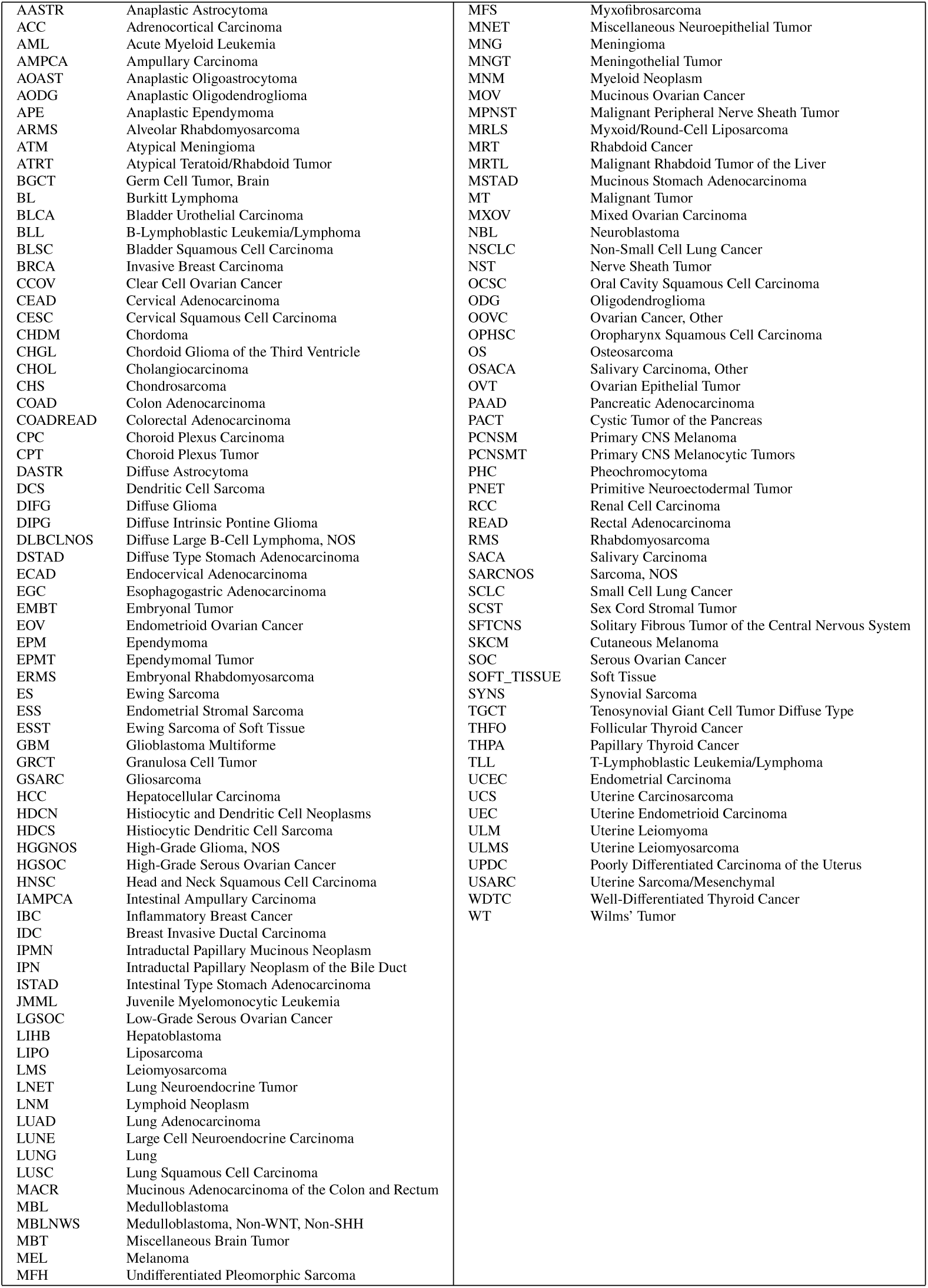
Disease abbreviations. All abbreviations are standard OncoTree ^42^ codes.

**Table S3:** Kinase target annotations for inhibitor agerafenib. Annotations are from various resources compared to published target affinity. Checkmarks denote proteins adjudicated by a given resource to be targets of the drug, while crosses denote proteins that are not indicated to be targets. The “Assay affinity” column denotes published affinity estimates ^162^.

| Target | DrugBank | OpenTargets | PubChem | Selleck Chem | Assay affinity |
| --- | --- | --- | --- | --- | --- |
| CSF-1R | X | X | X | ✓ | ✓ |
| c-Kit | X | X | X | ✓ | ✓ |
| EGFR | X | ✓ | ✓ | X | X |
| EPHA2 | X | X | ✓ | X | X |
| PDGFR | X | X | X | ✓ | ✓ |
| RAF | ✓ | ✓ | ✓ | ✓ | ✓ |
| RET | ✓ | ✓ | ✓ | ✓ | ✓ |

**Table S4:** Broad target annotation groupings for cell signaling and cell cycling comparisons.

| Category | Targets |
| --- | --- |
| Cell Signaling | Akt, PI3K, mTOR, Raf, FGFR, IGF-1R, Trk Receptor, c-Met, MEK, EGFR, HER2, Src, c-Kit, PKC, Wnt/beta-catenin, ALK, PDGFR, VEGFR, JAK, ERK, Gamma-secretase, TGF-beta/Smad, Syk, Hedgehog/Smoothed, STAT, TNF-alpha, c-RET, FLT3, CSF-1R, p38 MAPK, FAK, IκB/IKK, GSK-3, COX |
| Cell Cycling | ATM/ATR, Aurora Kinase, CDK, Chk, DNA/RNA Synthesis, Mdm2, Microtubule Associated, PLK, Proteasome, Topoisomerase |

**Table S5:** Coarse groupings for drug target annotations.

| Group | Targets |
| --- | --- |
| EGFR | EGFR, HER2 |
| FGFR/PDGFR/VEGFR | FGFR, PDGFR, VEGFR, c-Kit, c-RET, CSF-1R |
| IGF1R/MET | c-Met, IGF-1R |
| ALK/TRK | Trk Receptor, ALK |
| BCR-ABL/FLT3/AURK | Aurora Kinase, Bcr-Abl, FLT3 |
| Wnt/Hedgehog/Smo | Wnt/beta-catenin, Hedgehog/Smoothed |
| PI3K/AKT/mTOR | Akt, mTOR, PI3K |
| TNFα/TGFβ | TNF-alpha, TGF-beta/Smad |
| MEK/ERK | ERK, MEK |
| DNA synth/damage/repair | DHFR, DNA/RNA Synthesis, PLK, PARP, DNA Methyl-transferase, Topoisomerase, Microtubule Associated |
| Cell cycle arrest | PKC, Chk, Syk, CDK |
| Protein stability/degradation | E3 Ligase, Proteasome, HSP |
| Anti-androgenic | Estrogen/progestogen Receptor, Androgen Receptor, Aromatase |
| Metabolic | Glucocorticoid Receptor, HMG-CoA Reductase, Dehydrogenase |

**Table S6:** Abbreviations and acronyms for media additives.

| **Abbreviation/acronym** | **Full name** |
| --- | --- |
| BSA | Bovine serum albumin |
| EGF | Epidermal growth factor |
| FBS | Fetal bovine serum |
| FCS | Fetal calf serum |
| FGF2 | Fibroblast growth factor 2 |
| FGF10 | Fibroblast growth factor 10 |
| GM-CSF | Granulocyte-macrophage colony-stimulating factor |
| HEPES | 4-(2-hydroxyethyl)-1-piperazineethanesulfonic acid |
| NEAA | Non-essential amino acids |
| PDGF | Platelet-derived growth factor |
| Pen/Strep | Penicillin-Streptomycin |

**Supplementary Algorithm S1:**
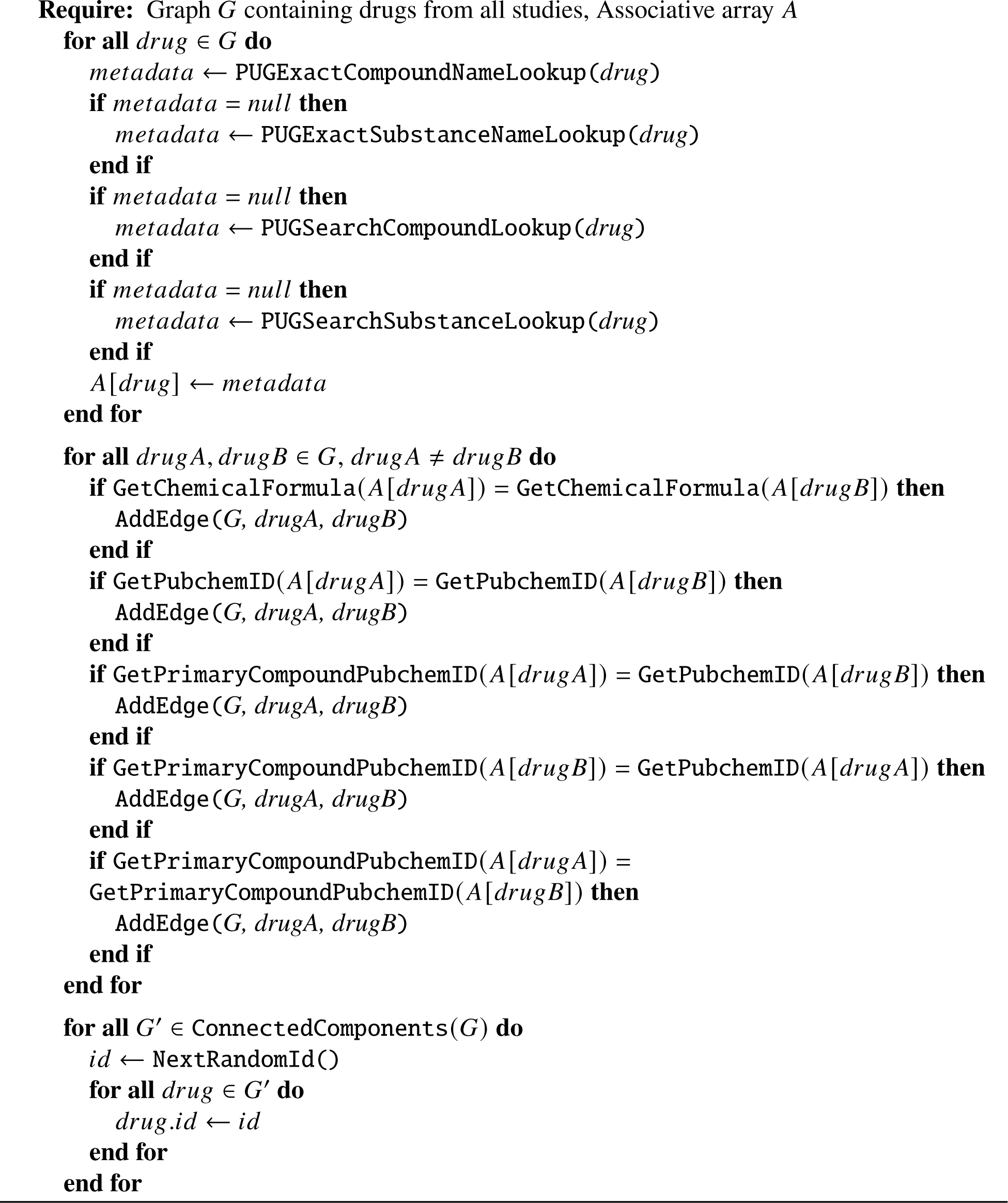
Cross-study drug harmonization.

## Footnotes

l https://github.com/tansey-lab/nf-rnaseq

## References

[1] Hervé Tiriac, Pascal Belleau, Dannielle D. Engle, Dennis Plenker, Astrid Deschênes, Tim D. D. Somerville, Fieke E. M. Froeling, Richard A. Burkhart, Robert E. Denroche, Gun-Ho Jang, Koji Miyabayashi, C. Megan Young, Hardik Patel, Michelle Ma, Joseph F. LaComb, Randze Lerie D. Palmaira, Ammar A. Javed, Jasmine C. Huynh, Molly Johnson, Kanika Arora, Nicolas Robine, Minita Shah, Rashesh Sanghvi, Austin B. Goetz, Cinthya Y. Lowder, Laura Martello, Else Driehuis, Nicolas LeComte, Gokce Askan, Christine A. Iacobuzio-Donahue, Hans Clevers, Laura D. Wood, Ralph H. Hruban, Elizabeth Thompson, Andrew J. Aguirre, Brian M. Wolpin, Aaron Sasson, Joseph Kim, Maoxin Wu, Juan Carlos Bucobo, Peter Allen, Divyesh V. Sejpal, William Nealon, James D. Sullivan, Jordan M. Winter, Phyllis A. Gimotty, Jean L. Grem, Dominick J. DiMaio, Jonathan M. Buscaglia, Paul M. Grandgenett, Jonathan R. Brody, Michael A. Hollingsworth, Grainne M. O’Kane, Faiyaz Notta, Edward Kim, James M. Crawford, Craig Devoe, Allyson Ocean, Christopher L. Wolfgang, Kenneth H. Yu, Ellen Li, Christopher R. Vakoc, Benjamin Hubert, Sandra E. Fischer, Julie M. Wilson, Richard Moffitt, Jennifer Knox, Alexander Krasnitz, Steven Gallinger, and David A. Tuveson. Organoid profiling identifies common responders to chemotherapy in pancreatic cancer. Cancer Discovery, 8(9):1112–1129, 2018.

[2] Ahmad Al Shihabi, Peyton J. Tebon, Huyen Thi Lam Nguyen, Jomjit Chantharasamee, Sara Sartini, Ardalan Davarifar, Alexandra Y Jensen, Miranda Diaz-Infante, Hannah Cox, Alfredo Gonzalez, Summer Swearingen, Nasrin Tavanaie, Sarah M Dry, Arun S. Singh, Bartosz Chmielowski, Joseph G. Crompton, Anusha Kalbasi, Fritz C. Eilber, Francis Hornicek, Nicholas M. Bernthal, Scott D. Nelson, Paul C. Boutros, Noah C Federman, Jane Yanagawa, and Alice Soragni. The landscape of drug sensitivity and resistance in sarcoma. Cell stem cell, 31(10):1524–1542, 2024.

[3] Norman Sachs and Hans Clevers. Organoid cultures for the analysis of cancer phenotypes. Current opinion in genetics & development, 24:68–73, 2014.

[4] Else Driehuis, Arne van Hoeck, Kat Moore, Sigrid Kolders, Hayley E. Francies, M. Can Gulersonmez, Edwin C. A. Stigter, Boudewĳn Burgering, Veerle Geurts, Ana Gracanin, Gergana Bounova, Folkert H. Morsink, Robert Vries, Sylvia Boj, Johan van Es, G. Johan A. Offerhaus, Onno Kranenburg, Mathew J. Garnett, Lodewyk Wessels, Edwin Cuppen, Lodewĳk A. A. Brosens, and Hans Clevers. Pancreatic cancer organoids recapitulate disease and allow personalized drug screening. Proceedings of the National Academy of Sciences, 116(52): 26580–26590, 2019.

[5] Ku-Geng Huo, Elisa D’Arcangelo, and Ming-Sound Tsao. Patient-derived cell line, xenograft and organoid models in lung cancer therapy. Translational Lung Cancer Research, 9(5):2214, 2020.

[6] Christine Yee, Kristie-Ann Dickson, Mohammed N Muntasir, Yue Ma, and Deborah J Marsh. Three-dimensional modelling of ovarian cancer: From cell lines to organoids for discovery and personalized medicine. Frontiers in Bioengineering and Biotechnology, 10:116, 2022.

[7] Katrin P. Guillen, Maihi Fujita, Andrew J. Butterfield, Sandra D. Scherer, Matthew H. Bailey, Zhengtao Chu, Yoko S. DeRose, Ling Zhao, Emilio Cortes-Sanchez, Chieh-Hsiang Yang, Jennifer Toner, Guoying Wang, Yi Qiao, Xiaomeng Huang, Jeffery A. Greenland, Jeffery M. Vahrenkamp, David H. Lum, Rachel E. Factor, Edward W. Nelson, Cindy B. Matsen, Jane M. Poretta, Regina Rosenthal, Anna C. Beck, Saundra S. Buys, Christos Vaklavas, John H. Ward, Randy L. Jensen, Kevin B. Jones, Zheqi Li, Steffi Oesterreich, Lacey E. Dobrolecki, Satya S. Pathi, Xing Yi Woo, Kristofer C. Berrett, Mark E. Wadsworth, Jeffrey H. Chuang, Michael T. Lewis, Gabor T. Marth, Jason Gertz, Katherine E. Varley, Bryan E. Welm, and Alana L. Welm. A human breast cancer-derived xenograft and organoid platform for drug discovery and precision oncology. Nature Cancer, 3(2):232–250, 2022.

[8] Gadi Lalazar, David Requena, Lavoisier Ramos-Espiritu, Denise Ng, Patrick D Bhola, Ype P De Jong, Ruisi Wang, Nicole JC Narayan, Bassem Shebl, Solomon Levin, et al. Identification of novel therapeutic targets for fibrolamellar carcinoma using patient-derived xenografts and direct-from-patient screening. Cancer Discovery, 11(10):2544–2563, 2021.

[9] Jin-Ku Lee, Zhaoqi Liu, Jason K. Sa, Sang Shin, Jiguang Wang, Mykola Bordyuh, Hee Jin Cho, Oliver Elliott, Timothy Chu, Seung Won Choi, Daniel I. S. Rosenbloom, In-Hee Lee, Yong Jae Shin, Hyun Ju Kang, Donggeon Kim, Sun Young Kim, Moon-Hee Sim, Jusun Kim, Taehyang Lee, Yun Jee Seo, Hyemi Shin, Mĳeong Lee, Sung Heon Kim, Yong-Jun Kwon, Jeong-Woo Oh, Minsuk Song, Misuk Kim, Doo-Sik Kong, Jung Won Choi, Ho Jun Seol, Jung-Il Lee, Seung Tae Kim, Joon Oh Park, Kyoung-Mee Kim, Sang-Yong Song, Jeong-Won Lee, Hee-Cheol Kim, Jeong Eon Lee, Min Gew Choi, Sung Wook Seo, Young Mog Shim, Jae Ill Zo, Byong Chang Jeong, Yeup Yoon, Gyu Ha Ryu, Nayoung K. D. Kim, Joon Seol Bae, Woong-Yang Park, Jeongwu Lee, Roel G. W. Verhaak, Antonio Iavarone, Jeeyun Lee, Raul Rabadan, and Do-Hyun Nam. Pharmacogenomic landscape of patient-derived tumor cells informs precision oncology therapy. Nature Genetics, 50(10):1399–1411, 2018.

[10] Patrik Johansson, Cecilia Krona, Soumi Kundu, Milena Doroszko, Sathishkumar Baskaran, Linnéa Schmidt, Claire Vinel, Elin Almstedt, Ramy Elgendy, Ludmila Elfineh, Caroline Gallant, Sara Lundsten, Fernando J. Ferrer Gago, Aleksi Hakkarainen, Petra Sipilä, Maria Häggblad, Ulf Martens, Bo Lundgren, Melanie M. Frigault, David P. Lane, Fredrik J. Swartling, Lene Uhrbom, Marika Nestor, Silvia Marino, and Sven Nelander. A patient-derived cell atlas informs precision targeting of glioblastoma. Cell Reports, 32(2), 2020.

[11] Christian K. Hirt, Tĳmen H. Booĳ, Linda Grob, Patrik Simmler, Nora C. Toussaint, David Keller, Doreen Taube, Vanessa Ludwig, Alexander Goryachkin, Chantal Pauli, Daniela Lenggenhager, Daniel J. Stekhoven, Christian U. Stirnimann, Katharina Endhardt, Femke Ringnalda, Lukas Villiger, Alexander Siebenhüner, Sofia Karkampouna, Marta De Menna, Janette Beshay, Hagen Klett, Marianna Kruithof-de Julio, Julia Schüler, and Gerald Schwank. Drug screening and genome editing in human pancreatic cancer organoids identifies drug-gene interactions and candidates for off-label therapy. Cell Genomics, 2(2), 2022.

[12] Shawn HR Lee, Wenjian Yang, Yoshihiro Gocho, August John, Lauren Rowland, Brandon Smart, Hannah Williams, Dylan Maxwell, Jeremy Hunt, Wentao Yang, et al. Pharmacotypes across the genomic landscape of pediatric acute lymphoblastic leukemia and impact on treatment response. Nature Medicine, 29(1):170–179, 2023.

[13] Amy Burd, Ross L Levine, Amy S Ruppert, Alice S Mims, Uma Borate, Eytan M Stein, Prapti Patel, Maria R Baer, Wendy Stock, Michael Deininger, et al. Precision medicine treatment in acute myeloid leukemia using prospective genomic profiling: feasibility and preliminary efficacy of the beat aml master trial. Nature Medicine, 26(12):1852–1858, 2020.

[14] Jason K. Sa, Jae Ryoung Hwang, Young-Jae Cho, Ji-Yoon Ryu, Jung-Joo Choi, Soo Young Jeong, Jihye Kim, Myeong Seon Kim, E. Sun Paik, Yoo-Young Lee, Chel Hun Choi, Tae-Joong Kim, Byoung-Gie Kim, Duk-Soo Bae, Yeri Lee, Nam-Gu Her, Yong Jae Shin, Hee Jin Cho, Ja Yeon Kim, Yun Jee Seo, Harim Koo, Jeong-Woo Oh, Taebum Lee, Hyun-Soo Kim, Sang Yong Song, Joon Seol Bae, Woong-Yang Park, Hee Dong Han, Hyung Jun Ahn, Anil K. Sood, Raul Rabadan, Jin-Ku Lee, Do-Hyun Nam, and Jeong-Won Lee. Pharmacogenomic analysis of patient-derived tumor cells in gynecologic cancers. Genome Biology, 20(1), 2019. ISSN 1474-760X. doi: 10.1186/s13059-019-1848-3.

[15] Filipe Correia Martins, Dominique-Laurent Couturier, Ines de Santiago, Carolin Margarethe Sauer, Maria Vias, Mihaela Angelova, Deborah Sanders, Anna Piskorz, James Hall, Karen Hosking, Anumithra Amirthanayagam, Sabina Cosulich, Larissa Carnevalli, Barry Davies, Thomas B. K. Watkins, Ionut G. Funingana, Helen Bolton, Krishnayan Haldar, John Latimer, Peter Baldwin, Robin Crawford, Matthew Eldridge, Bristi Basu, Mercedes Jimenez-Linan, Andrew W. Mcpherson, Nicholas McGranahan, Kevin Litchfield, Sohrab P. Shah, Iain McNeish, Carlos Caldas, Gerard Evan, Charles Swanton, and James D. Brenton. Clonal somatic copy number altered driver events inform drug sensitivity in high-grade serous ovarian cancer. Nature Communications, 13(1), 2022.

[16] Astrid Murumägi, Daniela Ungureanu, Suleiman Khan, Mariliina Arjama, Katja Välimäki, Aleksandr Ianevski, Philipp Ianevski, Rebecka Bergström, Alice Dini, Anna Kanerva, et al. Drug response profiles in patient-derived cancer cells across histological subtypes of ovarian cancer: real-time therapy tailoring for a patient with low-grade serous carcinoma. British Journal of Cancer, 128(4):678–690, 2022.

[17] Xuexue Xie, Xinyu Li, and Wei Song. Tumor organoid biobank-new platform for medical research. Scientific Reports, 13(1):1819, 2023.

[18] Saul E Acevedo, Tim M Stearns, Phillip Webster, Vivek Philip, Michael W Lloyd, Anuj Srivastava, Steven Neuhauser, Dale Begley, Debbie Krupke, Emily L Jocoy, et al. Pediatric preclinical in vivo testing (pivot) data portal enables access to 15 years of retrospective treatment study data in support of prospective study design. Cancer Research, 84(6_Supplement): 5468–5468, 2024.

[19] Aviad Tsherniak, Francisca Vazquez, Phil G. Montgomery, Barbara A. Weir, Gregory Kryukov, Glenn S. Cowley, Stanley Gill, William F. Harrington, Sasha Pantel, John M. Krill-Burger, Robin M. Meyers, Levi Ali, Amy Goodale, Yenarae Lee, Guozhi Jiang, Jessica Hsiao, William F.J. Gerath, Sara Howell, Erin Merkel, Mahmoud Ghandi, Levi A. Garraway, David E. Root, Todd R. Golub, Jesse S. Boehm, and William C. Hahn. Defining a cancer dependency map. Cell, 170(3):564–576, 2017. ISSN 0092-8674.

[20] Heike Peterziel, Nora Jamaladdin, Dina ElHarouni, Xenia F. Gerloff, Sonja Herter, Petra Fiesel, Yannick Berker, Mirjam Blattner-Johnson, Kathrin Schramm, Barbara C. Jones, David Reuss, Laura Turunen, Aileen Friedenauer, Tim Holland-Letz, Martin Sill, Lena Weiser, Christopher Previti, Gnanaprakash Balasubramanian, Nicolas U. Gerber, Johannes Gojo, Caroline Hutter, Ingrid Øra, Olli Lohi, Antonis Kattamis, Bram de Wilde, Frank Westermann, Stephan Tippelt, Norbert Graf, Michaela Nathrath, Monika Sparber-Sauer, Astrid Sehested, Christof M. Kramm, Uta Dirksen, Olli Kallioniemi, Stefan M. Pfister, Cornelis M. van Tilburg, David T. W. Jones, Jani Saarela, Vilja Pietiäinen, Natalie Jäger, Matthias Schlesner, Annette Kopp-Schneider, Sina Oppermann, Till Milde, Olaf Witt, and Ina Oehme. Drug sensitivity profiling of 3D tumor tissue cultures in the pediatric precision oncology program INFORM. NPJ Precision Oncology, 6(1), 2022.

[21] Loretta MS Lau, Chelsea Mayoh, Jinhan Xie, Paulette Barahona, Karen L MacKenzie, Marie Wong, Alvin Kamili, Maria Tsoli, Tim W Failes, Amit Kumar, et al. In vitro and in vivo drug screens of tumor cells identify novel therapies for high-risk child cancer. EMBO Molecular Medicine, 14(4):e14608, 2022.

[22] Arlet M Acanda De La Rocha, Noah E Berlow, Maggie Fader, Ebony R Coats, Cima Saghira, Paula S Espinal, Jeanette Galano, Ziad Khatib, Haneen Abdella, Ossama M Maher, et al. Feasibility of functional precision medicine for guiding treatment of relapsed or refractory pediatric cancers. Nature Medicine, 30(4):990–1000, 2024.

[23] Anthony Letai, Patrick Bhola, and Alana L Welm. Functional precision oncology: testing tumors with drugs to identify vulnerabilities and novel combinations. Cancer cell, 40(1): 26–35, 2022.

[24] Adam A. Friedman, Arnaud Amzallag, Iulian Pruteanu-Malinici, Subash Baniya, Zachary A. Cooper, Adriano Piris, Leeza Hargreaves, Vivien Igras, Dennie T. Frederick, Donald P. Lawrence, Daniel A. Haber, Keith T. Flaherty, Jennifer A. Wargo, Sridhar Ramaswamy, Cyril H. Benes, and David E. Fisher. Landscape of targeted anti-cancer drug synergies in melanoma identifies a novel BRAF-VEGFR/PDGFR combination treatment. PLOS ONE, 10 (10):e0140310, 2015.

[25] Ziyue Gu, Yanli Yao, Guizhu Yang, Guopei Zhu, Zhen Tian, Rui Wang, Qi Wu, Yujue Wang, Yaping Wu, Lan Chen, Chong Wang, Jiamin Gao, Xindan Kang, Jie Zhang, Lizhen Wang, Shengzhong Duan, Zhongming Zhao, Zhiyuan Zhang, and Shuyang Sun. Pharmacogenomic landscape of head and neck squamous cell carcinoma informs precision oncology therapy. Science Translational Medicine, 14(661):eabo5987, 2022.

[26] Disha Malani, Ashwini Kumar, Oscar Brück, Mika Kontro, Bhagwan Yadav, Monica Hellesøy, Heikki Kuusanmäki, Olli Dufva, Matti Kankainen, Samuli Eldfors, Swapnil Potdar, Jani Saarela, Laura Turunen, Alun Parsons, Imre Västrik, Katja Kivinen, Janna Saarela, Riikka Räty, Minna Lehto, Maĳa Wolf, Bjorn Tore Gjertsen, Satu Mustjoki, Tero Aittokallio, Krister Wennerberg, Caroline A. Heckman, Olli Kallioniemi, and Kimmo Porkka. Implementing a functional precision medicine tumor board for acute myeloid leukemia. Cancer Discovery, 12(2):388–401, 2021.

[27] Tea Pemovska, Mika Kontro, Bhagwan Yadav, Henrik Edgren, Samuli Eldfors, Agnieszka Szwajda, Henrikki Almusa, Maxim M. Bespalov, Pekka Ellonen, Erkki Elonen, Bjørn T. Gjertsen, Riikka Karjalainen, Evgeny Kulesskiy, Sonja Lagström, Anna Lehto, Maĳa Lepistö, Tuĳa Lundán, Muntasir Mamun Majumder, Jesus M. Lopez Marti, Pirkko Mattila, Astrid Murumägi, Satu Mustjoki, Aino Palva, Alun Parsons, Tero Pirttinen, Maria E. Rämet, Minna Suvela, Laura Turunen, Imre Västrik, Maĳa Wolf, Jonathan Knowles, Tero Aittokallio, Caroline A. Heckman, Kimmo Porkka, Olli Kallioniemi, and Krister Wennerberg. Individualized systems medicine strategy to tailor treatments for patients with chemorefractory acute myeloid leukemia. Cancer Discovery, 3(12):1416–1429, 2013.

[28] Daniel Bottomly, Nicola Long, Anna Reister Schultz, Stephen E. Kurtz, Cristina E. Tognon, Kara Johnson, Melissa Abel, Anupriya Agarwal, Sammantha Avaylon, Erik Benton, Aurora Blucher, Uma Borate, Theodore P. Braun, Jordana Brown, Jade Bryant, Russell Burke, Amy Carlos, Bill H. Chang, Hyun Jun Cho, Stephen Christy, Cody Coblentz, Aaron M. Cohen, Amanda d’Almeida, Rachel Cook, Alexey Danilov, Kim-Hien T. Dao, Michie Degnin, James Dibb, Christopher A. Eide, Isabel English, Stuart Hagler, Heath Harrelson, Rachel Henson, Hibery Ho, Sunil K. Joshi, Brian Junio, Andy Kaempf, Yoko Kosaka, Ted Laderas, Matt Lawhead, Hyunjung Lee, Jessica T. Leonard, Chenwei Lin, Evan F. Lind, Selina Qiuying Liu, Pierrette Lo, Marc M. Loriaux, Samuel Luty, Julia E. Maxson, Tara Macey, Jacqueline Martinez, Jessica Minnier, Andrea Monteblanco, Motomi Mori, Quinlan Morrow, Dylan Nelson, Justin Ramsdill, Angela Rofelty, Alexandra Rogers, Kyle A. Romine, Peter Ryabinin, Jennifer N. Saultz, David A. Sampson, Samantha L. Savage, Robert Schuff, Robert Searles, Rebecca L. Smith, Stephen E. Spurgeon, Tyler Sweeney, Ronan T. Swords, Aashis Thapa, Karina Thiel-Klare, Elie Traer, Jake Wagner, Beth Wilmot, Joelle Wolf, Guanming Wu, Amy Yates, Haĳiao Zhang, Christopher R. Cogle, Robert H. Collins, Michael W. Deininger, Christopher S. Hourigan, Craig T. Jordan, Tara L. Lin, Micaela E. Martinez, Rachel R. Pallapati, Daniel A. Pollyea, Anthony D. Pomicter, Justin M. Watts, Scott J. Weir, Brian J. Druker, Shannon K. McWeeney, and Jeffrey W. Tyner. Integrative analysis of drug response and clinical outcome in acute myeloid leukemia. Cancer Cell, 40(8):850–864, 2022.

[29] Chelsea Mayoh, Jie Mao, Jinhan Xie, Gábor Tax, Shu-Oi Chow, Roxanne Cadiz, Karina Pazaky, Paulette Barahona, Pamela Ajuyah, Peter Trebilcock, Angela Malquori, Kate Gunther, Anica Avila, Doo-Young Yun, Stephanie Alfred, Anjana Gopalakrishnan, Alvin Kamili, Marie Wong, Mark J. Cowley, Sophie Jessop, Loretta M S Lau, Toby N. Trahair, David S. Ziegler, Jamie I. Fletcher, Andrew J. Gifford, Maria Tsoli, Glenn M. Marshall, Michelle Haber, Vanessa Tyrrell, Tim W. Failes, Greg M Arndt, Richard B. Lock, Paul G. Ekert, and M Emmy M Dolman. High-throughput drug screening of primary tumor cells identifies therapeutic strategies for treating children with high-risk cancer. Cancer Research, 83: 2716–2732, 2023.

[30] Suk Hyung Lee, Wenhuo Hu, Justin T. Matulay, Mark V. Silva, Tomasz B. Owczarek, Kwanghee Kim, Chee Wai Chua, LaMont J. Barlow, Cyriac Kandoth, Alanna B. Williams, Sarah K. Bergren, Eugene J. Pietzak, Christopher B. Anderson, Mitchell C. Benson, Jonathan A. Coleman, Barry S. Taylor, Cory Abate-Shen, James M. McKiernan, Hikmat Al-Ahmadie, David B. Solit, and Michael M. Shen. Tumor evolution and drug response in patient-derived organoid models of bladder cancer. Cell, 173(2):515–528, 2018.

[31] Helen H.N. Yan, Hoi Cheong Siu, Simon Law, Siu Lun Ho, Sarah S.K. Yue, Wai Yin Tsui, Dessy Chan, April S. Chan, Stephanie Ma, Ka On Lam, Sina Bartfeld, Alice H.Y. Man, Bernard C.H. Lee, Annie S.Y. Chan, Jason W.H. Wong, Priscilla S.W. Cheng, Anthony K.W. Chan, Jiangwen Zhang, Jue Shi, Xiaodan Fan, Dora L.W. Kwong, Tak W. Mak, Siu Tsan Yuen, Hans Clevers, and Suet Yi Leung. A comprehensive human gastric cancer organoid biobank captures tumor subtype heterogeneity and enables therapeutic screening. Cell Stem Cell, 23(6):882–897, 2018.

[32] Kohta Toshimitsu, Ai Takano, Masayuki Fujii, Kazuhiro Togasaki, Mami Matano, Sirirat Takahashi, Takanori Kanai, and Toshiro Sato. Organoid screening reveals epigenetic vulnerabilities in human colorectal cancer. Nature Chemical Biology, 18(6):605–614, 2022.

[33] Johannes Betge, Niklas Rindtorff, Jan Sauer, Benedikt Rauscher, Clara Dingert, Haristi Gaitantzi, Frank Herweck, Kauthar Srour-Mhanna, Thilo Miersch, Erica Valentini, Kim E. Boonekamp, Veronika Hauber, Tobias Gutting, Larissa Frank, Sebastian Belle, Timo Gaiser, Inga Buchholz, Ralf Jesenofsky, Nicolai Härtel, Tianzuo Zhan, Bernd Fischer, Katja Breitkopf-Heinlein, Elke Burgermeister, Matthias P. Ebert, and Michael Boutros. The drug-induced phenotypic landscape of colorectal cancer organoids. Nature Communications, 13(1), 2022.

[34] Ling Li, Hildur Knutsdottir, Ken Hui, Matthew J. Weiss, Jin He, Benjamin Philosophe, Andrew M. Cameron, Christopher L. Wolfgang, Timothy M. Pawlik, Gabriel Ghiaur, Andrew J. Ewald, Esteban Mezey, Joel S. Bader, and Florin M. Selaru. Human primary liver cancer organoids reveal intratumor and interpatient drug response heterogeneity. JCI Insight, 4(2), 2019.

[35] Alice Boilève, Jérôme Cartry, Negaar Goudarzi, Sabrina Bedja, Jacques R.R. Mathieu, Mohamed Amine Bani, Remy Nicolle, Ali Mouawia, Ryme Bouyakoub, Claudio Nicotra, Maud Ngo-Camus, Bastien Job, Karélia Lipson, Valérie Boige, Marine Valéry, Anthony Tarabay, P. Dartigues, Lambros Tselikas, Thierry de Baère, A. Italiano, Simona Cosconea, Maximiliano Gelli, Elena Fernandez de Sevilla, Maxime Annereau, David Malka, Cristina Smolenschi, Michel Ducreux, Antoine Hollebecque, and Fanny Jaulin. Organoids for functional precision medicine in advanced pancreatic cancer. Gastroenterology, 167(5): 961–976, 2024.

[36] Lélia Polit, Jacques R.R. Mathieu, Cartry Jerome, Sabrina Bedja, Alice Boilève, Michel Ducreux, Fanny Jaulin, and Gustave Ronteix. Leveraging large-scale PDO-based assays to optimize antibody-drug conjugate efficacy in CRC. Journal of Clinical Oncology, 2025.

[37] Reid T. Powell, Abena Redwood, Xuan Liu, Lei Guo, Shirong Cai, Xinhui Zhou, Yizheng Tu, Xiaomei Zhang, Yuan Qi, Yan Jiang, Gloria Echeverria, Ningping Feng, XiaoYan Ma, Virginia Giuliani, Joseph R. Marszalek, Timothy P. Heffernan, Christopher P. Vellano, Jason B. White, Clifford Stephan, Peter J. Davies, Stacy Moulder, W. Fraser Symmans, Jeffrey T. Chang, and Helen Piwnica-Worms. Pharmacologic profiling of patient-derived xenograft models of primary treatment-naïve triple-negative breast cancer. Scientific Reports, 10(1), 2020.

[38] Alejandra Bruna, Oscar M. Rueda, Wendy Greenwood, Ankita Sati Batra, Maurizio Callari, Rajbir Nath Batra, Katherine Pogrebniak, Jose Sandoval, John W. Cassidy, Ana Tufegdzic-Vidakovic, Stephen-John Sammut, Linda Jones, Elena Provenzano, Richard Baird, Peter Eirew, James Hadfield, Matthew Eldridge, Anne McLaren-Douglas, Andrew Barthorpe, Howard Lightfoot, Mark J. O’Connor, Joe Gray, Javier Cortes, Jose Baselga, Elisabetta Marangoni, Alana L. Welm, Samuel Aparicio, Violeta Serra, Mathew J. Garnett, and Carlos Caldas. A biobank of breast cancer explants with preserved intra-tumor heterogeneity to screen anticancer compounds. Cell, 167(1):260–274, 2016.

[39] Adam A. Friedman, Yun Xia, Lorenzo Trippa, Long Phi Le, Vivien Igras, Dennie T. Frederick, Jennifer A. Wargo, Kenneth K. Tanabe, Donald P. Lawrence, Donna S. Neuberg, Keith T. Flaherty, and David E. Fisher. Feasibility of ultra-high-throughput functional screening of melanoma biopsies for discovery of novel cancer drug combinations. Clinical Cancer Research, 23(16):4680–4692, 2017.

[40] Ryan J. Ice, Michelle Chen, Max Sidorov, Tam Le Ho, Rinette W. L. Woo, Aida Rodriguez-Brotons, Tri Luu, Damon Jian, Kevin B. Kim, Stanley P. Leong, HanKyul Kim, Angela Kim, Des Stone, Ari Nazarian, Alyssia Oh, Gregory J. Tranah, Mehdi Nosrati, David de Semir, Altaf A. Dar, Stephen Chang, Pierre-Yves Desprez, Mohammed Kashani-Sabet, Liliana Soroceanu, and Sean D. McAllister. Drug responses are conserved across patient-derived xenograft models of melanoma leading to identification of novel drug combination therapies. British Journal of Cancer, 122(5):648–657, 2019.

[41] Loretta M S Lau, Chelsea Mayoh, Jinhan Xie, Paulette Barahona, Karen L MacKenzie, Marie Wong, Alvin Kamili, Maria Tsoli, Tim W Failes, Amit Kumar, Emily V A Mould, Andrew Gifford, Shu-Oi Chow, Mark Pinese, Jamie I Fletcher, Greg M Arndt, Dong-Anh Khuong-Quang, Carol Wadham, Daniel Batey, Georgina Eden, Peter Trebilcock, Swapna Joshi, Stephanie Alfred, Anjana Gopalakrishnan, Aaminah Khan, Dylan Grebert Wade, Patrick A Strong, Elodie Manouvrier, Lisa T Morgan, Miriam Span, Jin Yi Lim, Roxanne Cadiz, Caitlin Ung, David M Thomas, Katherine M Tucker, Meera Warby, Geoffrey B McCowage, Luciano Dalla-Pozza, Jennifer A Byrne, Federica Saletta, Andrew Fellowes, Stephen B Fox, Murray D Norris, Vanessa Tyrrell, Toby N Trahair, Richard B Lock, Mark J Cowley, Paul G Ekert, Michelle Haber, David S Ziegler, and Glenn M Marshall. In vitro and in vivo drug screens of tumor cells identify novel therapies for high-risk child cancer. EMBO Molecular Medicine, 14(4), 2021.

[42] Ritika Kundra, Hongxin Zhang, Robert Sheridan, Sahussapont Joseph Sirintrapun, Avery Wang, Angelica Ochoa, Manda Wilson, Benjamin Gross, Yichao Sun, Ramyasree Madupuri, Baby A Satravada, Dalicia Reales, Efsevia Vakiani, Hikmat A Al-Ahmadie, Ahmet Dogan, Maria Arcila, Ahmet Zehir, Steven Maron, Michael F Berger, Cristina Viaplana, Katherine Janeway, Matthew Ducar, Lynette Sholl, Snjezana Dogan, Philippe Bedard, Lea F Surrey, Iker Huerga Sanchez, Aĳaz Syed, Anoop Balakrishnan Rema, Debyani Chakravarty, Sarah Suehnholz, Moriah Nissan, Gopakumar V Iyer, Rajmohan Murali, Nancy Bouvier, Robert A Soslow, David Hyman, Anas Younes, Andrew Intlekofer, James J Harding, Richard D Carvajal, Paul J Sabbatini, Ghassan K Abou-Alfa, Luc Morris, Yelena Y Janjigian, Meighan M Gallagher, Tara A Soumerai, Ingo K Mellinghoff, Abraham A Hakimi, Matthew Fury, Jason T Huse, Aditya Bagrodia, Meera Hameed, Stacy Thomas, Stuart Gardos, Ethan Cerami, Tali Mazor, Priti Kumari, Pichai Raman, Priyanka Shivdasani, Suzanne MacFarland, Scott Newman, Angela Waanders, Jianjiong Gao, David Solit, and Nikolaus Schultz. OncoTree: A cancer classification system for precision oncology. JCO Clinical Cancer Informatics, 5:221–230, Feb 2021.

[43] Bastien Nguyen, Christopher Fong, Anisha Luthra, Shaleigh A Smith, Renzo G DiNatale, Subhiksha Nandakumar, Henry Walch, Walid K Chatila, Ramyasree Madupuri, Ritika Kundra, Craig M Bielski, Brooke Mastrogiacomo, Mark T A Donoghue, Adrienne Boire, Sarat Chandarlapaty, Karuna Ganesh, James J Harding, Christine A Iacobuzio-Donahue, Pedram Razavi, Ed Reznik, Charles M Rudin, Dmitriy Zamarin, Wassim Abida, Ghassan K Abou-Alfa, Carol Aghajanian, Andrea Cercek, Ping Chi, Darren Feldman, Alan L Ho, Gopakumar Iyer, Yelena Y Janjigian, Michael Morris, Robert J Motzer, Eileen M O’Reilly, Michael A Postow, Nitya P Raj, Gregory J Riely, Mark E Robson, Jonathan E Rosenberg, Anton Safonov, Alexander N Shoushtari, William Tap, Min Yuen Teo, Anna M Varghese, Martin Voss, Rona Yaeger, Marjorie G Zauderer, Nadeem Abu-Rustum, Julio Garcia-Aguilar, Bernard Bochner, Abraham Hakimi, William R Jarnagin, David R Jones, Daniela Molena, Luc Morris, Eric Rios-Doria, Paul Russo, Samuel Singer, Vivian E Strong, Debyani Chakravarty, Lora H Ellenson, Anuradha Gopalan, Jorge S Reis-Filho, Britta Weigelt, Marc Ladanyi, Mithat Gonen, Sohrab P Shah, Joan Massague, Jianjiong Gao, Ahmet Zehir, Michael F Berger, David B Solit, Samuel F Bakhoum, Francisco Sanchez-Vega, and Nikolaus Schultz. Genomic characterization of metastatic patterns from prospective clinical sequencing of 25,000 patients. Cell, 185(3):563–575, Feb 2022.

[44] Selleck Chemicals. Inhibitor Library, 2025. URL https://www.selleckchem.com/screening/inhibitor-library.html.

[45] Selleck Chemicals. FDA-approved drug library, 2025. URL https://www.selleckchem.com/screening/fda-approved-drug-library.html.

[46] Selleck Chemicals. Bioactive Compound Library-I, 2025. URL https://www.selleckchem.com/screening/chemical-library.html.

[47] Timothy J Ley, Li Ding, Matthew J Walter, Michael D McLellan, Tamara Lamprecht, David E Larson, Cyriac Kandoth, Jacqueline E Payton, Jack Baty, John Welch, Christopher C Harris, Cheryl F Lichti, R Reid Townsend, Robert S Fulton, David J Dooling, Daniel C Koboldt, Heather Schmidt, Qunyuan Zhang, John R Osborne, Ling Lin, Michelle O’Laughlin, Joshua F McMichael, Kim D Delehaunty, Sean D McGrath, Lucinda A Fulton, Vincent J Magrini, Tammi L Vickery, Jasreet Hundal, Lisa L Cook, Joshua J Conyers, Gary W Swift, Jerry P Reed, Patricia A Alldredge, Todd Wylie, Jason Walker, Joelle Kalicki, Mark A Watson, Sharon Heath, William D Shannon, Nobish Varghese, Rakesh Nagarajan, Peter Westervelt, Michael H Tomasson, Daniel C Link, Timothy A Graubert, John F DiPersio, Elaine R Mardis, and Richard K Wilson. DNMT3A mutations in acute myeloid leukemia. N. Engl. J. Med., 363(25):2424–2433, December 2010.

[48] Brunangelo Falini, Lorenzo Brunetti, Paolo Sportoletti, and Maria Paola Martelli. NPM1-mutated acute myeloid leukemia: from bench to bedside. Blood, 136(15):1707–1721, October 2020.

[49] Tuan Nguyen, Giang TT Nguyen, Thin Nguyen, and Duc-Hau Le. Graph convolutional networks for drug response prediction. IEEE/ACM transactions on computational biology and bioinformatics, 19(1):146–154, 2021.

[50] Pengfei Liu, Hongjian Li, Shuai Li, and Kwong-Sak Leung. Improving prediction of phenotypic drug response on cancer cell lines using deep convolutional network. BMC bioinformatics, 20(1), 2019.

[51] Fangfang Xia, Jonathan E. Allen, Prasanna Balaprakash, Thomas S. Brettin, Cristina Garcia-Cardona, Austin R. Clyde, Judith D. Cohn, James H. Doroshow, Xiaotian Duan, Veronika B. Dubinkina, Yvonne A. Evrard, Ya Ju Fan, Jason D. Gans, Stewart He, Pinyi Lu, Sergei Maslov, Alexander Partin, Maulik Shukla, Eric A. Stahlberg, Justin M. Wozniak, Hyun Seung Yoo, George Zaki, Yitan Zhu, and Rick L. Stevens. A cross-study analysis of drug response prediction in cancer cell lines. Briefings in Bioinformatics, 23, 2021.

[52] Likun Jiang, Changzhi Jiang, Xinyu Yu, Rao Fu, Shuting Jin, and Xiangrong Liu. DeepTTA: a transformer-based model for predicting cancer drug response. Briefings in bioinformatics, 23(3), 2022.

53. Aaron van den Oord, Yazhe Li, and Oriol Vinyals. Representation learning with contrastive predictive coding. arXiv preprint arXiv:1807.03748, 2018.

[54] Leland McInnes and John Healy. UMAP: Uniform manifold approximation and projection for dimension reduction. arXiv preprint arXiv:1802.03426, 2018.

[55] Dalia Kamel, Christopher Gray, Jagdeep Singh Walia, and Vikaash Kumar. PARP inhibitor drugs in the treatment of breast, ovarian, prostate and pancreatic cancers: An update of clinical trials. Current Drug Targets, 19(1), January 2018. ISSN 1389-4501.

[56] Bradley Efron. Large-scale simultaneous hypothesis testing: The choice of a null hypothesis. Journal of the American Statistical Association, 99(465):96–104, 2004.

[57] Bradley Efron. Large-scale inference: Empirical Bayes methods for estimation, testing, and prediction. 2012.

[58] T Dragovich, D Laheru, F Dayyani, V Bolejack, L Smith, J Seng, H Burris, P Rosen, M Hidalgo, P Ritch, A F Baker, N Raghunand, J Crowley, and D D Von Hoff. Phase II trial of vatalanib in patients with advanced or metastatic pancreatic adenocarcinoma after first-line gemcitabine therapy (PCRT o4-001). Cancer chemotherapy and pharmacology, 74 (2):379–387, 2014.

[59] A Mittelman, S Savona, C Puccio, H Chun, T Ahmed, E Feldman, P Sullivan, P Arnold, and Z Arlin. Phase II trial of fludarabine phosphate (F-ara-AMP) in patients with advanced head and neck cancer. Investigational new drugs, 8(S1):S65–S67, March 1990.

[60] Vincent Grégoire, K Kian Ang, Jean-François Rosier, Marc Beauduin, Adam S Garden, Marc Hamoir, Walter N Hittelman, Yves Humblet, Fadlo R Khuri, Luka Milas, Carine Mitine, and Pierre Scalliet. A phase I study of fludarabine combined with radiotherapy in patients with intermediate to locally advanced head and neck squamous cell carcinoma. Radiotherapy and oncology, 63(2):187–193, May 2002.

[61] Turang E Behbahani, Eben L Rosenthal, William B Parker, and Eric J Sorscher. Intratumoral generation of 2-fluoroadenine to treat solid malignancies of the head and neck. Head Neck, 41(6):1979–1983, June 2019.

[62] Nicolas A Fraunhoffer, Analía Meilerman Abuelafia, Martin Bigonnet, Odile Gayet, Julie Roques, Emmanuel Telle, Patricia Santofimia-Castaño, María Teresa Borrello, Eduardo Chuluyan, Nelson Dusetti, and Juan Iovanna. Evidencing a pancreatic ductal adenocarcinoma subpopulation sensitive to the proteasome inhibitor carfilzomib. Clinical Cancer Research, 26(20):5506–5519, October 2020.

[63] Y Liu, W Wu, W Hong, X Sun, J Wu, and Q Huang. Raltitrexed-based chemotherapy for advanced colorectal cancer. Clin. Res. Hepatol. Gastroenterol., 38(2):219–225, April 2014.

[64] Bo Hwa Sohn, Sung Hwan Lee, Yun Seong Jeong, and Ju-Seog Lee. Clinical implication of EZH2 inhibitors in hepatocellular carcinoma. Cancer Research, 83(7_Supplement): 6139–6139, 2023.

[65] Lola Jade Palmieri, Sophie Cousin, Mariella Spalato, Jean Philippe Guégan, Alban Bessede, Simon Pernot, and Antoine Italiano. Targeting EZH2 to overcome the resistance to immunotherapy in microsatellite stable colorectal cancer: Results from the CAIRE study. Journal of Clinical Oncology, 41(16 supp):3599–3599, June 2023. ISSN 1527-7755.

[66] Apostolia-Maria Tsimberidou, Francis Giles, Madeleine Duvic, Luis Fayad, and Razelle Kurzrock. Phase II study of pentostatin in advanced t-cell lymphoid malignancies: Update of an M.D. Anderson Cancer Center series. Cancer, 100(2):342–349, January 2004.

[67] Peter J O’Dwyer, Barbara Wagner, Brian R. Leyland-Jones, Robert Wittes, Bruce D. Cheson, and Daniel F. Hoth. 2’-deoxycoformycin (pentostatin) for lymphoid malignancies. Annals of internal medicine, 108(5):733, May 1988.

[68] U.S. Food and Drug Administration. FDA approves Cotellic as part of combination treatment for advanced melanoma, 2015. URL https://wayback.archive-it.org/7993/20161022101159/ http://www.fda.gov/NewsEvents/Newsroom/PressAnnouncements/ucm471934.htm.

[69] U.S. Food and Drug Administration. FDA approves Mekinist in combination with Tafinlar for advanced melanoma, 2014. URL https://wayback.archive-it.org/7993/20161023125600/ http://www.fda.gov/NewsEvents/Newsroom/PressAnnouncements/ucm381159.htm.

[70] U.S. Food and Drug Administration. FDA approves encorafenib and binimetinib in combination for unresectable or metastatic melanoma with BRAF mutations, 2018. URL https://www.fda.gov/drugs/resources-information-approved-drugs/fda-approves-encorafenib-and-binimetinib-combination-unresectable-or-metastatic-

[71] U.S. Food and Drug Administration. FDA approves selumetinib for neurofibromatosis type 1 with symptomatic, inoperable plexiform neurofibromas, 2020. URL https://www.fda.gov/drugs/resources-information-approved-drugs/fda-approves-selumetinib-neurofibromatosis-type-1-symptomatic-inoperable-plexifo

[72] Michael J Fisher, Jaishri O Blakeley, Brian D Weiss, Eva Dombi, Shivani Ahlawat, Srivandana Akshintala, Allan J Belzberg, Miriam Bornhorst, Miriam A Bredella, Wenli Cai, Rosalie E Ferner, Andrea M Gross, Gordon J Harris, Robert Listernick, Ina Ly, Staci Martin, Victor F Mautner, Johannes M Salamon, Kilian E Salerno, Robert J Spinner, Verena Staedtke, Nicole J Ullrich, Meena Upadhyaya, Pamela L Wolters, Kaleb Yohay, and Brigitte C Widemann. Management of neurofibromatosis type 1-associated plexiform neurofibromas. Neurooncology, 24(11):1827–1844, November 2022.

[73] Adrienne D Cox and Channing J Der. The RAF inhibitor paradox revisited. Cancer Cell, 21 (2):147–149, February 2012.

[74] Harry P. Erba, Pau Montesinos, Hee-Je Kim, Elżbieta Patkowska, Radovan Vrhovac, Pavel Žák, Po-Nan Wang, Tsvetomir Mitov, James Hanyok, Yasser Mostafa Kamel, Jaime E. Connolly Rohrbach, Li Liu, Aziz Benzohra, Arnaud Lesegretain, Jorge Cortes, Alexander E. Perl, Mikkael A. Sekeres, Hervé Dombret, Sergio Amadori, Jianxiang Wang, Mark J. Levis, and Richard F. Schlenk. Quizartinib plus chemotherapy in newly diagnosed patients with FLT3-internal-tandem-duplication-positive acute myeloid leukaemia (QuANTUM-first): A randomised, double-blind, placebo-controlled, phase 3 trial. The Lancet, 401(10388): 1571–1583, May 2023.

[75] Diarmuid M Moran and Carl G Maki. Nutlin-3a induces cytoskeletal rearrangement and inhibits the migration and invasion capacity of p53 wild-type cancer cells. Mol Cancer Ther, 9(4):895–905, April 2010.

[76] U.S. Food and Drug Administration. FDA approves quizartinib for newly diagnosed acute myeloid leukemia, 2023. URL https://www.fda.gov/drugs/drug-approvals-and-databases/fda-approves-quizartinib-newly-diagnosed-acute-myeloid-leukemia.

[77] U.s. food and drug administration. FDA approves drug for adults with rare form of bone marrow disorder, 2022. URL https://www.fda.gov/drugs/news-events-human-drugs/fda-approves-drug-adults-rare-form-bone-marrow-disorder.

[78] Astex Pharmaceutical. Astex Pharmaceuticals Discontinues Amuvatinib Clinical Development Program, 2012. URL https://astx.com/astex-pharmaceuticals-discontinues-amuvatinib-clinical-development-program/.

[79] Robert Williams. Discontinued drugs in 2012: Oncology drugs. Expert opinion on investigational drugs, 22(12):1627–1644, December 2013.

[80] Adis Insight. Drug Profile: ENMD 2076, 2020. Accessed: May 22, 2025.

[81] Adis Insight. Drug Profile: KW 2449, 2023. Accessed: May 22, 2025.

[82] National Comprehensive Cancer Network, Inc. NCCN Guidelines: Treatment by Cancer Type, 2024. URL https://www.nccn.org/guidelines/category_1. Accessed May 2024.

[83] DB Longley and PG Johnston. Molecular mechanisms of drug resistance. The Journal of Pathology, 205(2):275–292, 2005.

[84] Wei-Hsin Lin, Yi-Wen Chang, Min-Xiang Hong, Te-Cheng Hsu, Ko-Chuan Lee, Che Lin, and Jia-Lin Lee. STAT3 phosphorylation at Ser727 and Tyr705 differentially regulates the EMT–MET switch and cancer metastasis. Oncogene, 40:791–805, 2020.

[85] Mohammad Zahid Kamran, Prachi Patil, and Rajiv P Gude. Role of STAT3 in cancer metastasis and translational advances. BioMed research international, 2013(1), 2013.

[86] Esther P.B. Rodman, Michael J. Emch, Xiaonan Hou, Archit Bajaj, Nicole A. Pearson, August J. John, Yamillie Ortiz, Adam D. Bass, Saloni Singh, Gustavo Baldassarre, Scott H. Kaufmann, S. John Weroha, and John R. Hawse. Lestaurtinib’s antineoplastic activity converges on JAK/STAT signaling to inhibit treatment naïve and therapy resistant forms ovarian cancer. NPJ Precision Oncology, 9, 2025.

[87] Audrey Lamora, Julie Talbot, Mathilde Mullard, Benedicte Brounais-Le Royer, Françoise Redini, and Franck Verrecchia. TGF-*β* signaling in bone remodeling and osteosarcoma progression. J. Clin. Med., 5(11):96, November 2016.

[88] Rongrong Ge and Gavin M. Huang. Targeting transforming growth factor beta signaling in metastatic osteosarcoma. Journal of Bone Oncology, 43:100513, December 2023. ISSN 2212-1374.

[89] Emma D. Wrenn, April A. Apfelbaum, Erin R. Rudzinski, Xuemei Deng, Wei Jiang, Sudha Sud, Raelene A. Van Noord, Erika A. Newman, Nicolas M. Garcia, Aya Miyaki, Virginia J. Hoglund, Shruti S. Bhise, Sami B. Kanaan, Olivia G. Waltner, Scott N. Furlan, and Elizabeth R. Lawlor. Cancer-associated fibroblast-like tumor cells remodel the Ewing sarcoma tumor microenvironment. Clinical Cancer Research, 29(24):5140–5154, July 2023. ISSN 1557-3265.

[90] Kristina Y. Aguilera and David W. Dawson. WNT ligand dependencies in pancreatic cancer. Frontiers in Cell and Developmental Biology, 9, April 2021. ISSN 2296-634X.

[91] Gat Rauner, Piyush B Gupta, and Charlotte Kuperwasser. From 2D to 3D and beyond: The evolution and impact of in vitro tumor models in cancer research. Nature Methods, pages 1–12, 2025.

[92] Steven M. Corsello, Rohith T. Nagari, Ryan D. Spangler, Jordan Rossen, Mustafa Kocak, Jordan G. Bryan, Ranad Humeidi, David Peck, Xiaoyun Wu, Andrew A. Tang, Vickie M. Wang, Samantha A. Bender, Evan Lemire, Rajiv Narayan, Philip Montgomery, Uri Ben-David, Colin W. Garvie, Yejia Chen, Matthew G. Rees, Nicholas J. Lyons, James M. McFarland, Bang T. Wong, Li Wang, Nancy Dumont, Patrick J. O’Hearn, Eric Stefan, John G. Doench, Caitlin N. Harrington, Heidi Greulich, Matthew Meyerson, Francisca Vazquez, Aravind Subramanian, Jennifer A. Roth, Joshua A. Bittker, Jesse S. Boehm, Christopher C. Mader, Aviad Tsherniak, and Todd R. Golub. Discovering the anticancer potential of non-oncology drugs by systematic viability profiling. Nature Cancer, 1(2):235–248, 2020.

[93] Nishanth Ulhas Nair, Patricia Greninger, Xiaohu Zhang, Adam A. Friedman, Arnaud Amzallag, Eliane Cortez, Avinash Das Sahu, Joo Sang Lee, Anahita Dastur, Regina K. Egan, Ellen Murchie, Michele Ceribelli, Giovanna S. Crowther, Erin Beck, Joseph McClanaghan, Carleen Klump-Thomas, Jessica L. Boisvert, Leah J. Damon, Kelli M. Wilson, Jeffrey Ho, Angela Tam, Crystal McKnight, Sam Michael, Zina Itkin, Mathew J. Garnett, Jeffrey A. Engelman, Daniel A. Haber, Craig J. Thomas, Eytan Ruppin, and Cyril H. Benes. A landscape of response to drug combinations in non-small cell lung cancer. Nature Communications, 14(1), 2023.

[94] Susan L. Holbeck, Richard Camalier, James A. Crowell, Jeevan Prasaad Govindharajulu, Melinda Hollingshead, Lawrence W. Anderson, Eric Polley, Larry Rubinstein, Apurva Srivastava, Deborah Wilsker, Jerry M. Collins, and James H. Doroshow. The National Cancer Institute ALMANAC: A comprehensive screening resource for the detection of anticancer drug pairs with enhanced therapeutic activity. Cancer Research, 77(13):3564–3576, 2017.

[95] Wanjuan Yang, Jorge Soares, Patricia Greninger, Elena J. Edelman, Howard Lightfoot, Simon Forbes, Nidhi Bindal, Dave Beare, James A. Smith, I. Richard Thompson, Sridhar Ramaswamy, P. Andrew Futreal, Daniel A. Haber, Michael R. Stratton, Cyril Benes, Ultan McDermott, and Mathew J. Garnett. Genomics of Drug Sensitivity in Cancer (GDSC): A resource for therapeutic biomarker discovery in cancer cells. Nucleic Acids Research, 41(D1): D955–D961, 2013.

[96] Amrita Basu, Nicole E. Bodycombe, Jaime H. Cheah, Edmund V. Price, Ke Liu, Giannina I. Schaefer, Richard Y. Ebright, Michelle L. Stewart, Daisuke Ito, Stephanie Wang, Abigail L. Bracha, Ted Liefeld, Mathias Wawer, Joshua C. Gilbert, Andrew J. Wilson, Nicolas Stransky, Gregory V. Kryukov, Vlado Dancik, Jordi Barretina, Levi A. Garraway, C. Suk-Yee Hon, Benito Munoz, Joshua A. Bittker, Brent R. Stockwell, Dineo Khabele, Andrew M. Stern, Paul A. Clemons, Alykhan F. Shamji, and Stuart L. Schreiber. An interactive resource to identify cancer genetic and lineage dependencies targeted by small molecules. Cell, 154(5): 1151–1161, 2013.

[97] Brinton Seashore-Ludlow, Matthew G. Rees, Jaime H. Cheah, Murat Cokol, Edmund V. Price, Matthew E. Coletti, Victor Jones, Nicole E. Bodycombe, Christian K. Soule, Joshua Gould, Benjamin Alexander, Ava Li, Philip Montgomery, Mathias J. Wawer, Nurdan Kuru, Joanne D. Kotz, C. Suk-Yee Hon, Benito Munoz, Ted Liefeld, Vlado Dančík, Joshua A. Bittker, Michelle Palmer, James E. Bradner, Alykhan F. Shamji, Paul A. Clemons, and Stuart L. Schreiber. Harnessing connectivity in a large-scale small-molecule sensitivity dataset. Cancer Discovery, 5(11):1210–1223, 2015.

[98] Jennifer O’Neil, Yair Benita, Igor Feldman, Melissa Chenard, Brian Roberts, Yaping Liu, Jing Li, Astrid Kral, Serguei Lejnine, Andrey Loboda, William Arthur, Razvan Cristescu, Brian B. Haines, Christopher Winter, Theresa Zhang, Andrew Bloecher, and Stuart D. Shumway. An unbiased oncology compound screen to identify novel combination strategies. Molecular Cancer Therapeutics, 15(6):1155–1162, 2016. ISSN 1538-8514. doi: 10.1158/1535-7163. mct-15-0843.

[99] Patricia Jaaks, Elizabeth A. Coker, Daniel J. Vis, Olivia Edwards, Emma F. Carpenter, Simonetta M. Leto, Lisa Dwane, Francesco Sassi, Howard Lightfoot, Syd Barthorpe, Dieudonne van der Meer, Wanjuan Yang, Alexandra Beck, Tatiana Mironenko, Caitlin Hall, James Hall, Iman Mali, Laura Richardson, Charlotte Tolley, James Morris, Frances Thomas, Ermira Lleshi, Nanne Aben, Cyril H. Benes, Andrea Bertotti, Livio Trusolino, Lodewyk Wessels, and Mathew J. Garnett. Effective drug combinations in breast, colon and pancreatic cancer cells. Nature, 603(7899):166–173, 2022.

[100] Jordi Barretina, Giordano Caponigro, Nicolas Stransky, Kavitha Venkatesan, Adam A. Margolin, Sungjoon Kim, Christopher J. Wilson, Joseph Lehár, Gregory V. Kryukov, Dmitriy Sonkin, Anupama Reddy, Manway Liu, Lauren Murray, Michael F. Berger, John E. Monahan, Paula Morais, Jodi Meltzer, Adam Korejwa, Judit Jané-Valbuena, Felipa A. Mapa, Joseph Thibault, Eva Bric-Furlong, Pichai Raman, Aaron Shipway, Ingo H. Engels, Jill Cheng, Guoying K. Yu, Jianjun Yu, Peter Aspesi, Melanie de Silva, Kalpana Jagtap, Michael D. Jones, Li Wang, Charles Hatton, Emanuele Palescandolo, Supriya Gupta, Scott Mahan, Carrie Sougnez, Robert C. Onofrio, Ted Liefeld, Laura MacConaill, Wendy Winckler, Michael Reich, Nanxin Li, Jill P. Mesirov, Stacey B. Gabriel, Gad Getz, Kristin Ardlie, Vivien Chan, Vic E. Myer, Barbara L. Weber, Jeff Porter, Markus Warmuth, Peter Finan, Jennifer L. Harris, Matthew Meyerson, Todd R. Golub, Michael P. Morrissey, William R. Sellers, Robert Schlegel, and Levi A. Garraway. The Cancer Cell Line Encyclopedia enables predictive modelling of anticancer drug sensitivity. Nature, 483(7391):603–607, 2012.

[101] Eric Polley, Mark Kunkel, David Evans, Thomas Silvers, Rene Delosh, Julie Laudeman, Chad Ogle, Russell Reinhart, Michael Selby, John Connelly, Erik Harris, Nicole Fer, Dmitriy Sonkin, Gurmeet Kaur, Anne Monks, Shakun Malik, Joel Morris, and Beverly A. Teicher. Small cell lung cancer screen of oncology drugs, investigational agents, and gene and microRNA expression. Journal of the National Cancer Institute, 108(10), 2016.

[102] Amos Bairoch. The Cellosaurus, a cell-line knowledge resource. Journal of biomolecular techniques: JBT, 29(2):25, 2018.

[103] Henning Karlsson, Mårten Fryknäs, Rolf Larsson, and Peter Nygren. Loss of cancer drug activity in colon cancer HCT-116 cells during spheroid formation in a new 3-D spheroid cell culture system. Experimental Cell Research, 318(13):1577–1585, 2012.

[104] Alicja Urbaniak, Sergio Piña-Oviedo, Youzhong Yuan, Adam Huczyński, and Timothy C Chambers. Limitations of an ex vivo breast cancer model for studying the mechanism of action of the anticancer drug paclitaxel. European Journal of Pharmacology, 891:173780, 2021.

[105] Zofia M Komar, Nicole S Verkaik, Ahmed Dahmani, Elodie Montaudon, Roland Kanaar, Adriaan B Houtsmuller, Agnes Jager, Elisabetta Marangoni, and Dik C van Gent. Development and validation of a functional ex vivo paclitaxel and eribulin sensitivity assay for breast cancer, the REMIT assay. NPJ Breast Cancer, 11(1):17, 2025.

[106] Yi-He Ling, Yandan Yang, Carmen Tornos, Balraj Singh, and Roman Perez-Soler. Paclitaxel-induced apoptosis is associated with expression and activation of c-Mos gene product in human ovarian carcinoma SKOV3 cells. Cancer research, 58(16):3633–3640, 1998.

[107] Tin Myo Khing, Won Seok Choi, Dong Min Kim, Wah Wah Po, Wynn Thein, Chang Yell Shin, and Uy Dong Sohn. The effect of paclitaxel on apoptosis, autophagy and mitotic catastrophe in AGS cells. Scientific reports, 11(1):23490, 2021.

[108] Emiliano Cocco, Maurizio Scaltriti, and Alexander E. Drilon. NTRK fusion-positive cancers and TRK inhibitor therapy. Nature Reviews Clinical Oncology, 15:731–747, 2018.

[109] Christian Rolfo, Rossana Ruiz, Elisa Giovannetti, Ignacio Gil-Bazo, Antonio Russo, Francesco Passiglia, Marco Giallombardo, Marc Peeters, and Luis Raez. Entrectinib: a potent new TRK, ROS1, and ALK inhibitor. *Expert opinion on investigational drugs*, 24(11):1493–1500, 2015.

[110] Jeffrey J Kooĳman, Wilhelmina E. van Riel, Jelle Dylus, Martine B. W. Prinsen, Yvonne Grobben, Tessa J. J. de Bitter, Antoon M. van Doornmalen, Janneke J.T.M. Melis, Joost C. M. Uitdehaag, Yugo Narumi, Yusuke Kawase, Jeroen A.D.M. de Roos, Nicole Willemsen-Seegers, and Guido J. R. Zaman. Comparative kinase and cancer cell panel profiling of kinase inhibitors approved for clinical use from 2018 to 2020. Frontiers in Oncology, 12, 2022.

[111] Oriol Pich, Chris Bailey, Thomas BK Watkins, Simone Zaccaria, Mariam Jamal-Hanjani, and Charles Swanton. The translational challenges of precision oncology. Cancer Cell, 40 (5):458–478, 2022.

[112] Jianzhu Ma, Samson H Fong, Yunan Luo, Christopher J Bakkenist, John Paul Shen, Soufiane Mourragui, Lodewyk FA Wessels, Marc Hafner, Roded Sharan, Jian Peng, et al. Few-shot learning creates predictive models of drug response that translate from high-throughput screens to individual patients. Nature Cancer, 2(2):233–244, 2021.

[113] Brent M Kuenzi, Jisoo Park, Samson H Fong, Kyle S Sanchez, John Lee, Jason F Kreisberg, Jianzhu Ma, and Trey Ideker. Predicting drug response and synergy using a deep learning model of human cancer cells. Cancer cell, 38(5):672–684, 2020.

[114] Fatemeh Rafiei, Hojjat Zeraati, Karim Abbasi, Jahan B Ghasemi, Mahboubeh Parsaeian, and Ali Masoudi-Nejad. DeepTraSynergy: Drug combinations using multimodal deep learning with transformers. Bioinformatics, 39(8), 2023.

[115] Mohamed Reda El Khili, Safyan Aman Memon, and Amin Emad. MARSY: A multitask deep-learning framework for prediction of drug combination synergy scores. Bioinformatics, 39(4), 2023.

[116] Iljung Jin and Hojung Nam. HiDRA: Hierarchical network for drug response prediction with attention. Journal of Chemical Information and Modeling, 61(8):3858–3867, 2021.

[117] Tianyu Liu, Tinyi Chu, Xiao Luo, and Hongyu Zhao. Building a unified model for drug synergy analysis powered by large language models. Nature Communications, 16(1):4537, 2025.

[118] Jacob Devlin, Ming-Wei Chang, Kenton Lee, and Kristina Toutanova. BERT: Pre-training of deep bidirectional transformers for language understanding. In Proceedings of the 2019 conference of the North American chapter of the association for computational linguistics: human language technologies, volume 1 (long and short papers), pages 4171–4186, 2019.

[119] Noah Hollmann, Samuel Müller, Lennart Purucker, Arjun Krishnakumar, Max Körfer, Shi Bin Hoo, Robin Tibor Schirrmeister, and Frank Hutter. Accurate predictions on small data with a tabular foundation model. Nature, 637(8045):319–326, 2025.

[120] Tianqi Chen and Carlos Guestrin. XGBoost: A scalable tree boosting system. In Proceedings of the 22nd ACM SIGKDD International Conference on Knowledge Discovery and Data Mining, pages 785––794, 2016.

[121] Rishi Bommasani, Drew A. Hudson, Ehsan Adeli, Russ Altman, Simran Arora, Sydney von Arx, Michael S. Bernstein, Jeannette Bohg, Antoine Bosselut, Emma Brunskill, Erik Brynjolfsson, Shyamal Buch, Dallas Card, Rodrigo Castellon, Niladri Chatterji, Annie Chen, Kathleen Creel, Jared Quincy Davis, Dora Demszky, Chris Donahue, Moussa Doumbouya, Esin Durmus, Stefano Ermon, John Etchemendy, Kawin Ethayarajh, Li Fei-Fei, Chelsea Finn, Trevor Gale, Lauren Gillespie, Karan Goel, Noah Goodman, Shelby Grossman, Neel Guha, Tatsunori Hashimoto, Peter Henderson, John Hewitt, Daniel E. Ho, Jenny Hong, Kyle Hsu, Jing Huang, Thomas Icard, Saahil Jain, Dan Jurafsky, Pratyusha Kalluri, Siddharth Karamcheti, Geoff Keeling, Fereshte Khani, Omar Khattab, Pang Wei Koh, Mark Krass, Ranjay Krishna, Rohith Kuditipudi, Ananya Kumar, Faisal Ladhak, Mina Lee, Tony Lee, Jure Leskovec, Isabelle Levent, Xiang Lisa Li, Xuechen Li, Tengyu Ma, Ali Malik, Christopher D. Manning, Suvir Mirchandani, Eric Mitchell, Zanele Munyikwa, Suraj Nair, Avanika Narayan, Deepak Narayanan, Ben Newman, Allen Nie, Juan Carlos Niebles, Hamed Nilforoshan, Julian Nyarko, Giray Ogut, Laurel Orr, Isabel Papadimitriou, Joon Sung Park, Chris Piech, Eva Portelance, Christopher Potts, Aditi Raghunathan, Rob Reich, Hongyu Ren, Frieda Rong, Yusuf Roohani, Camilo Ruiz, Jack Ryan, Christopher Ré, Dorsa Sadigh, Shiori Sagawa, Keshav Santhanam, Andy Shih, Krishnan Srinivasan, Alex Tamkin, Rohan Taori, Armin W. Thomas, Florian Tramèr, Rose E. Wang, William Wang, Bohan Wu, Jiajun Wu, Yuhuai Wu, Sang Michael Xie, Michihiro Yasunaga, Jiaxuan You, Matei Zaharia, Michael Zhang, Tianyi Zhang, Xikun Zhang, Yuhui Zhang, Lucia Zheng, Kaitlyn Zhou, and Percy Liang. On the opportunities and risks of foundation models. arXiv preprint arXiv:2108.07258, 2021.

[122] Arthur Liberzon, Chet Birger, Helga Thorvaldsdóttir, Mahmoud Ghandi, Jill P Mesirov, and Pablo Tamayo. The molecular signatures database hallmark gene set collection. Cell systems, 1(6):417–425, 2015.

[123] Scott J. Dixon and Michael J. Lee. Quick tips for interpreting cell death experiments. Nature Cell Biology, 25(12):1720–1723, 2023.

[124] Marc Hafner, Mario Niepel, Mirra Chung, and Peter K Sorger. Growth rate inhibition metrics correct for confounders in measuring sensitivity to cancer drugs. Nature Methods, 13(6): 521–527, 2016.

[125] Peyton J Tebon, Bowen Wang, Alexander L Markowitz, Ardalan Davarifar, Brandon L Tsai, Patrycja Krawczuk, Alfredo E Gonzalez, Sara Sartini, Graeme F Murray, Huyen Thi Lam Nguyen, et al. Drug screening at single-organoid resolution via bioprinting and interferometry. Nature Communications, 14(1):3168, 2023.

[126] Bassel Alsaed, Johannes Smolander, Hanna Laitinen, Linh Lin, Nina Bobik, Lilja Lahtinen, Mikko Räsänen, Shadi Jansouz, Karita Peltonen, Emmi Jokinen, Jay Klievink, Keerthana Ganesh, Mari Ainola, Eva Sutinen, Mikko Rönty, Elli Narvi, Anil Thotakura, Pipsa Saharinen, Satu Mustjoki, Ilkka K Ilonen, and Heidi M. Haikala. Ex vivo modeling of precision immuno-oncology responses in lung cancer. Science Advances, 10, 2024.

[127] Alexandra Sontheimer-Phelps, Bryan A Hassell, and Donald E Ingber. Modelling cancer in microfluidic human organs-on-chips. Nature reviews cancer, 19(2):65–81, 2019.

[128] Mohammad Jouybar, Charlotte M de Winde, Katarina Wolf, Peter Friedl, Reina E Mebius, and Jaap MJ den Toonder. Cancer-on-chip models for metastasis: Importance of the tumor microenvironment. Trends in biotechnology, 42(4):431–448, 2024.

[129] Ramya Sivakumar, Marina Chan, Jiye Stella Shin, Nao Nishida-Aoki, Heidi L Kenerson, Olivier Elemento, Himisha Beltran, Raymond Yeung, and Taranjit S Gujral. Organotypic tumor slice cultures provide a versatile platform for immuno-oncology and drug discovery. Oncoimmunology, 8(12), 2019.

[130] Paraskevi Dimou, Sumita Trivedi, Maria Liousia, Reena R D’Souza, and Astero Klampatsa. Precision-cut tumor slices (PCTS) as an ex vivo model in immunotherapy research. Antibodies, 11(2):26, 2022.

[131] Shimrit Mayer, Tomer Milo, Achinoam Isaacson, Coral Halperin, Shoval Miyara, Yaniv Stein, Chen Lior, Meirav Pevsner-Fischer, Eldad Tzahor, Avi Mayo, et al. The tumor microenvironment shows a hierarchy of cell-cell interactions dominated by fibroblasts. Nature communications, 14(1), 2023.

[132] Dakai Yang, Jing Liu, Hui Qian, and Qin Zhuang. Cancer-associated fibroblasts: from basic science to anticancer therapy. Experimental & Molecular Medicine, 55(7):1322–1332, 2023.

[133] Hans Raskov, Adile Orhan, Jan Pravsgaard Christensen, and Ismail Gögenur. Cytotoxic CD8+ T cells in cancer and cancer immunotherapy. British journal of cancer, 124(2):359–367, 2021.

[134] Charlotte J Imianowski, Qiang Chen, Creg J Workman, and Dario AA Vignali. Regulatory T cells in the tumour microenvironment. Nature Reviews Cancer, pages 1–20, 2025.

[135] Mathilde Bied, William W Ho, Florent Ginhoux, and Camille Blériot. Roles of macrophages in tumor development: A spatiotemporal perspective. Cellular & molecular immunology, 20 (9):983–992, 2023.

[136] Ming-Zhu Jin and Wei-Lin Jin. The updated landscape of tumor microenvironment and drug repurposing. Signal transduction and targeted therapy, 5(1):166, 2020.

[137] Yidi Qu, Bo Dou, Horyue Tan, Yibin Feng, Ning Wang, and Di Wang. Tumor microenvironment-driven non-cell-autonomous resistance to antineoplastic treatment. Molecular cancer, 18(1):69, 2019.

[138] Alberto Mantovani, Paola Allavena, Federica Marchesi, and Cecilia Garlanda. Macrophages as tools and targets in cancer therapy. Nature reviews drug discovery, 21(11):799–820, 2022.

[139] Nicholas Sioutos, Sherri de Coronado, Margaret W. Haber, Frank W. Hartel, Wen-Ling Shaiu, and Lawrence W. Wright. NCI Thesaurus: A semantic model integrating cancer-related clinical and molecular information. Journal of Biomedical Informatics, 40(1):30–43, 2007. ISSN 1532-0464.

[140] Mahul B Amin, Frederick L Greene, Stephen B Edge, Carolyn C Compton, Jeffrey E Gershenwald, Robert K Brookland, Laura Meyer, Donna M Gress, David R Byrd, and David P Winchester. The eighth edition AJCC cancer staging manual: Continuing to build a bridge from a population-based to a more “personalized”’ approach to cancer staging. CA: A cancer journal for clinicians, 67(2):93–99, 2017.

[141] Johan T den Dunnen, Raymond Dalgleish, Donna R Maglott, Reece K Hart, Marc S Greenblatt, Jean McGowan-Jordan, Anne-Francoise Roux, Timothy Smith, Stylianos E Antonarakis, and Peter E M Taschner. HGVS recommendations for the description of sequence variants: 2016 update. Hum Mutat, 37(6):564–569, Jun 2016.

[142] Ino de Bruĳn, Xiang Li, Selcuk Onur Sumer, Benjamin Gross, Robert Sheridan, Angelica Ochoa, Manda Wilson, Avery Wang, Hongxin Zhang, Aaron Lisman, Adam Abeshouse, Emily Zhang, Alice Thum, Ananthan Sadagopan, Zachary Heins, Cyriac Kandoth, Sander Rodenburg, Sander Tan, Pieter Lukasse, Sjoerd van Hagen, Remond J A Fĳneman, Gerrit A Meĳer, Nikolaus Schultz, and Jianjiong Gao. Genome Nexus: A comprehensive resource for the annotation and interpretation of genomic variants in cancer. JCO Clinical Cancer Informatics, 6, Feb 2022.

[143] Ngak-Leng Sim, Prateek Kumar, Jing Hu, Steven Henikoff, Georg Schneider, and Pauline C. Ng. SIFT web server: Predicting effects of amino acid substitutions on proteins. Nucleic Acids Research, 40(W1):W452–W457, 06 2012.

[144] Ivan Adzhubei, Daniel M Jordan, and Shamil R Sunyaev. Predicting functional effect of human missense mutations using PolyPhen-2. Current protocols in human genetics, 76(1): 7–20, 2013.

[145] Sarah P Suehnholz, Moriah H Nissan, Hongxin Zhang, Ritika Kundra, Subhiksha Nandakumar, Calvin Lu, Stephanie Carrero, Amanda Dhaneshwar, Nicole Fernandez, Benjamin W Xu, Maria E Arcila, Ahmet Zehir, Aĳazuddin Syed, A Rose Brannon, Julia E Rudolph, Eder Paraiso, Paul J Sabbatini, Ross L Levine, Ahmet Dogan, Jianjiong Gao, Marc Ladanyi, Alexander Drilon, Michael F Berger, David B Solit, Nikolaus Schultz, and Debyani Chakravarty. Quantifying the expanding landscape of clinical actionability for patients with cancer. Cancer Discovery, 14(1):49–65, Jan 2024.

[146] Valerie A Schneider, Tina Graves-Lindsay, Kerstin Howe, Nathan Bouk, Hsiu-Chuan Chen, Paul A Kitts, Terence D Murphy, Kim D Pruitt, Françoise Thibaud-Nissen, Derek Albracht, Robert S Fulton, Milinn Kremitzki, Vincent Magrini, Chris Markovic, Sean McGrath, Karyn Meltz Steinberg, Kate Auger, William Chow, Joanna Collins, Glenn Harden, Timothy Hubbard, Sarah Pelan, Jared T Simpson, Glen Threadgold, James Torrance, Jonathan M Wood, Laura Clarke, Sergey Koren, Matthew Boitano, Paul Peluso, Heng Li, Chen-Shan Chin, Adam M Phillippy, Richard Durbin, Richard K Wilson, Paul Flicek, Evan E Eichler, and Deanna M Church. Evaluation of GRCh38 and de novo haploid genome assemblies demonstrates the enduring quality of the reference assembly. Genome research, 27(5): 849–864, May 2017.

[147] Paolo Di Tommaso, Maria Chatzou, Evan W Floden, Pablo Prieto Barja, Emilio Palumbo, and Cedric Notredame. Nextflow enables reproducible computational workflows. Nature Biotechnology, 35(4):316–319, 2017.

[148] Simon Andrews, Felix Krueger, Anne Segonds-Pichon, Laura Biggins, Christel Krueger, and Steven Wingett. FastQC. Babraham Institute, January 2012.

[149] Shifu Chen, Yanqing Zhou, Yaru Chen, and Jia Gu. fastp: An ultra-fast all-in-one FASTQ preprocessor. Bioinformatics, 34(17):i884–i890, 09 2018. ISSN 1367-4803.

[150] Alexander Dobin, Carrie A. Davis, Felix Schlesinger, Jorg Drenkow, Chris Zaleski, Sonali Jha, Philippe Batut, Mark Chaisson, and Thomas R. Gingeras. STAR: Ultrafast universal RNA-seq aligner. Bioinformatics, 29(1):15–21, 10 2012.

[151] Luis R. Nassar, Galt P. Barber, Anna Benet-Pagès, Jonathan Casper, Hiram Clawson, Mark E. Diekhans, Clay Fischer, Jairo Navarro Gonzalez, A. Hinrichs, Brian T. Lee, Christopher M. Lee, Pranav Muthuraman, Beagan Nguy, Tiana Pereira, Parisa Nejad, Gerardo Perez, Brian J. Raney, Daniel Schmelter, Matthew L. Speir, Brittney D. Wick, Ann S. Zweig, David Haussler, Robert M. Kuhn, Maximilian Haeussler, and W. James Kent. The UCSC Genome Browser database: 2023 update. Nucleic Acids Research, 51(D1):D1188–D1195, 11 2022.

[152] Adam Frankish, Silvia Carbonell Sala, Mark E. Diekhans, Irwin Jungreis, Jane E. Loveland, Jonathan M. Mudge, Cristina Sisu, James C. Wright, Carme Arnan, If H. A. Barnes, Abhimanyu Banerjee, Ruth Bennett, Andrew E. Berry, Alexandra Bignell, Carles Boix, Ferriol Calvet Riera, Daniel Cerdán-Vélez, Fiona Cunningham, Claire Davidson, Sarah M. Donaldson, Cagatay Dursun, Reham Fatima, Stefano Giorgetti, Carlos García-Girón, Jose Manuel Gonzalez, Matthew Hardy, Peter W. Harrison, Thibaut Hourlier, Zoe Hollis, Toby Hunt, Benjamin T. James, Yunzhe Jiang, Rory Johnson, Mike P. Kay, Julien Lagarde, Fergal J. Martin, Laura Martínez Gómez, Surag Nair, Pengyu Ni, Fernando Pozo, Vivek Ramalingam, Magali Ruffier, Bianca M. Schmitt, Jacob Meir Schreiber, Emily Steed, Marie-Marthe Suner, Dulika Sumathipala, Irina Sycheva, Barbara Uszczynska-Ratajczak, Elizabeth Wass, Yucheng T. Yang, Andrew D. Yates, Zahoor Zafrulla, Jyoti Choudhary, Mark B. Gerstein, Roderic Guigó, Tim J. P. Hubbard, Manolis Kellis, Anshul B Kundaje, Benedict Paten, Michael L. Tress, and Paul Flicek. GENCODE: reference annotation for the human and mouse genomes in 2023. Nucleic Acids Research, 51(D1):D942–D949, 11 2022.

[153] Heng Li, Bob Handsaker, Alec Wysoker, Tim Fennell, Jue Ruan, Nils Homer, Gabor Marth, Goncalo Abecasis, Richard Durbin, and 1000 Genome Project Data Processing Subgroup. The Sequence Alignment/Map format and SAMtools. Bioinformatics, 25(16):2078–2079, 06 2009.

[154] Alex Rubinsteyn, Tavi Nathanson, Julia Kodysh, Tim O’Donnell, Arun Ahuja, Jeff Hammerbacher, B. Arman Aksoy, Brent Pedersen-Bioinformatics, Valentin Grouès, and Isaac Hodes. hammerlab/pyensembl: Version 1.1.0, 2017. URL https://zenodo.org/record/822502.

[155] Sunghwan Kim, Paul A Thiessen, Tiejun Cheng, Bo Yu, and Evan E Bolton. An update on PUG-REST: RESTful interface for programmatic access to PubChem. Nucleic Acids Research, 46(W1):W563–W570, 2018.

[156] Sunghwan Kim, Paul A Thiessen, Evan E Bolton, Jie Chen, Gang Fu, Asta Gindulyte, Lianyi Han, Jane He, Siqian He, Benjamin A Shoemaker, Jiyao Wang, Bo Yu, Jian Zhang, and Stephen H Bryant. PubChem substance and compound databases. Nucleic acids research, 44 (D1):D1202–13, Jan 2016.

[157] David Weininger. SMILES, a chemical language and information system. 1. introduction to methodology and encoding rules. Journal of chemical information and computer sciences, 28(1):31–36, 1988.

[158] Evan E. Bolton, Yanli Wang, Paul A. Thiessen, and Stephen H. Bryant. PubChem: Integrated platform of small molecules and biological activities. Annual Reports in Computational Chemistry, 4:217–241, 2008.

[159] Craig Knox, Mike Wilson, Christen M Klinger, Mark Franklin, Eponine Oler, Alex Wilson, Allison Pon, Jordan Cox, Na Eun Lucy Chin, Seth A Strawbridge, Marysol Garcia-Patino, Ray Kruger, Aadhavya Sivakumaran, Selena Sanford, Rahil Doshi, Nitya Khetarpal, Omolola Fatokun, Daphnee Doucet, Ashley Zubkowski, Dorsa Yahya Rayat, Hayley Jackson, Karxena Harford, Afia Anjum, Mahi Zakir, Fei Wang, Siyang Tian, Brian Lee, Jaanus Liigand, Harrison Peters, Ruo Qi Rachel Wang, Tue Nguyen, Denise So, Matthew Sharp, Rodolfo da Silva, Cyrella Gabriel, Joshua Scantlebury, Marissa Jasinski, David Ackerman, Timothy Jewison, Tanvir Sajed, Vasuk Gautam, and David S Wishart. DrugBank 6.0: The DrugBank knowledgebase for 2024. Nucleic acids research, 52(D1):D1265–D1275, January 2024.

[160] Annalisa Buniello, Daniel Suveges, Carlos Cruz-Castillo, Manuel Bernal Llinares, Helena Cornu, Irene Lopez, Kirill Tsukanov, Juan María Roldán-Romero, Chintan Mehta, Luca Fumis, Graham McNeill, James D Hayhurst, Ricardo Esteban Martinez Osorio, Ehsan Barkhordari, Javier Ferrer, Miguel Carmona, Prashant Uniyal, Maria J Falaguera, Polina Rusina, Ines Smit, Jeremy Schwartzentruber, Tobi Alegbe, Vivien W Ho, Daniel Considine, Xiangyu Ge, Szymon Szyszkowski, Yakov Tsepilov, Maya Ghoussaini, Ian Dunham, David G Hulcoop, Ellen M McDonagh, and David Ochoa. Open Targets Platform: Facilitating therapeutic hypotheses building in drug discovery. Nucleic Acids Res., 53(D1):D1467–D1475, January 2025.

[161] Susanne Müller, Apirat Chaikuad, Nathanael S Gray, and Stefan Knapp. The ins and outs of selective kinase inhibitor development. Nat. Chem. Biol., 11(11):818–821, November 2015.

[162] Martin W Rowbottom, Raffaella Faraoni, Qi Chao, Brian T Campbell, Andiliy G Lai, Eduardo Setti, Maiko Ezawa, Kelly G Sprankle, Sunny Abraham, Lan Tran, Brian Struss, Michael Gibney, Robert C Armstrong, Ruwanthi N Gunawardane, Ronald R Nepomuceno, Ianina Valenta, Helen Hua, Michael F Gardner, Merryl D Cramer, Dana Gitnick, Darren E Insko, Julius L Apuy, Susan Jones-Bolin, Arup K Ghose, Torsten Herbertz, Mark A Ator, Bruce D Dorsey, Bruce Ruggeri, Michael Williams, Shripad Bhagwat, Joyce James, and Mark W Holladay. Identification of 1-(3-(6,7-dimethoxyquinazolin-4-yloxy)phenyl)-3-(5-(1,1,1-trifluoro-2-methylpropan-2-yl)isoxazol-3-yl)urea hydrochloride (CEP-32496), a highly potent and orally efficacious inhibitor of V-RAF murine sarcoma viral oncogene homologue B1 (BRAF) V600E. Journal of medicinal chemistry, 55(3):1082–1105, February 2012.

[163] WHO Drug Information. International nonproprietary names for pharmaceutical substances (inn): proposed inn list 131. WHO Drug Information, 38(2), 2024.

[164] Florent De Bruyne, Arnaud Ponçon, Joris Giai, Xavier Dode, David Darmon, Cyrille Colin, François Gueyffier, and Laurent Letrilliart. INN or brand name drug prescriptions: a multilevel, cross-sectional study in general practice. European Journal of Clinical Pharmacology, 75(2): 275–283, October 2018.

[165] Anita Bandrowski, Matthew Brush, Jeffrey S. Grethe, et al. The resource identification initiative: A cultural shift in publishing. F1000Research, 4:134, 2015.

[166] Christopher J. Mungall, Carlo Torniai, Georgios V. Gkoutos, Suzanna E. Lewis, and Melissa A. Haendel. Uberon, an integrative multi-species anatomy ontology. Genome Biology, 13(1): R5, 2012.

[167] Matthew D Hoffman, David M Blei, Chong Wang, and John Paisley. Stochastic variational inference. Journal of Machine Learning Research, 2013.

[168] Diederik P Kingma and Jimmy Ba. Adam: A method for stochastic optimization. arXiv preprint arXiv:1412.6980, 2014.

[169] David Rogers and Mathew Hahn. Extended-connectivity fingerprints. Journal of chemical information and modeling, 50(5):742–754, 2010.

